# Thirty years of fluconazole therapy selects an azole-resistant *Candida albicans* isolate with a pre-adapted physiological, metabolic and structural state

**DOI:** 10.64898/2026.09.19.752827

**Authors:** Claudia Parra-Giraldo, Julio Jesús Estrada-Valbuena, Yerly Vargas-Casanova, Irene Aldea, Luis Felipe Clemente, Concha Gil, Raquel Martínez-López, Lucía Monteoliva

## Abstract

Azole resistance in *Candida albicans* is traditionally attributed to alterations in drug targets and efflux mechanisms; however, how long-term antifungal exposure reshapes fungal physiology remains incompletely understood. Here, we characterize a fluconazole-resistant *C. albicans* clinical isolate (PUJ256) recovered from a patient with chronic mucocutaneous candidiasis after more than 30 years of continuous fluconazole therapy. Compared with the reference strain SC5314, the resistant isolate exhibited a clear fitness trade-off, with reduced filamentation and increased membrane vulnerability under basal conditions, yet enhanced fitness in the presence of fluconazole. Integration of phenotypic analyses with label-free quantitative proteomics revealed extensive metabolic remodelling, including coordinated regulation of central carbon metabolism, ergosterol biosynthesis and redox homeostasis. Notably, mitochondrial membrane potential was preserved in the resistant isolate under fluconazole stress, whereas the susceptible strain exhibited mitochondrial depolarization together with activation of MAPK signalling pathways. In addition, the resistant isolate displayed reduced susceptibility to phagocytosis under antifungal exposure, consistent with an increased capacity for persistence in the context of ongoing treatment. Collectively, our findings indicate that long-term azole resistance in *C. albicans* PUJ256 is associated with a stable, pre-adapted physiological state characterized by metabolic reprogramming and mitochondrial resilience. The prolonged antifungal exposure of this isolate provides a valuable opportunity to explore adaptative strategies that extend beyond canonical mechanisms, pointing to mitochondrial function and metabolic plasticity as potential targets for therapeutic intervention.

**Graphical abstract.:** 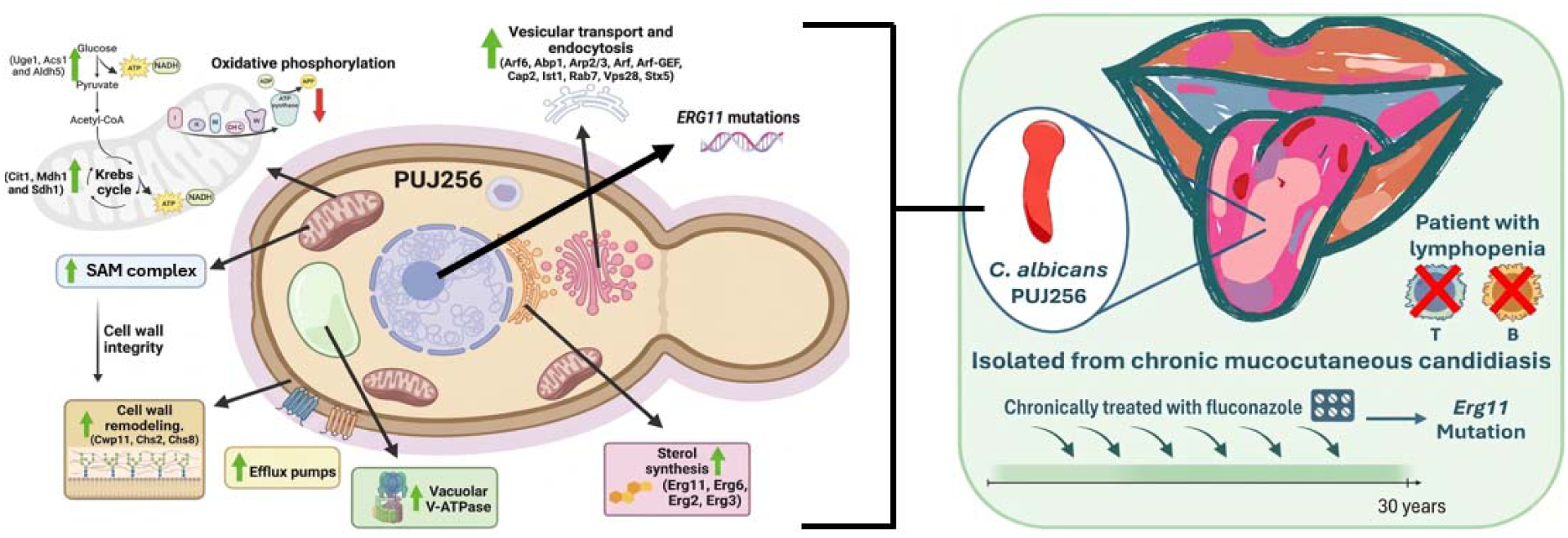

## Introduction

*Candida albicans* was first recognized as a cause of human disease in the 19^th^ century, with early clinical reports identifying it in chronic mucocutaneous lesions and systemic infections among hospitalized and immunocompromised patients (Parrot J., 1869). These observations established *C. albicans* as a clinically relevant opportunistic pathogen capable of persistent colonization and recurrent disease in diverse host environments. Over the latter 20^th^ and early 21^st^ centuries, *C. albicans* has emerged as one of the most prevalent causes of fungal infections worldwide, reflecting both its biological adaptability and evolutionary success in human hosts. Accordingly, *C. albicans* is now included in the World Health Organization (WHO) fungal priority pathogens list as a high-priority fungal pathogen, highlighting its major impact on global public health (paper WHO)

The widespread clinical use of azole antifungals, particularly fluconazole, revolutionized the management of candidiasis due to their favourable oral bioavailability, broad spectrum, and relatively low toxicity. Fluconazole became a first-line therapy for mucosal and systemic *Candida* infections beginning in the 1980s, leading to a decline in mortality from invasive candidiasis but simultaneously imposing a strong selective pressure on fungal populations. As a result, azole-resistant *C. albicans* isolates have become increasingly prevalent in clinical settings, especially among patients with long-term or recurrent antifungal exposure (Pfaller and Diekema, 2007; Andes and Safdar, 2012).

Azoles inhibit lanosterol 14α-demethylase (encoded by *ERG11*), a critical enzyme in the ergosterol biosynthesis pathway, thereby disrupting membrane integrity and interfering with fungal growth. Canonical mechanisms of azole resistance in *C. albicans* include point mutations in *ERG11* that reduce drug binding, upregulation of ATP-binding cassette (ABC) and major facilitator superfamily (MFS) efflux pumps, and adaptive changes in membrane sterol composition. While these mechanisms explain azole tolerance at the molecular level, they do not fully capture the broader physiological and fitness changes observed in many resistant clinical isolates (White et al., 1998; Sanglard et al., 2002).

Beyond direct effects on membrane biosynthesis, fluconazole and other azoles exert pleiotropic influences on fungal cellular physiology, often involving mitochondrial function, redox balance, and stress signalling pathways. Mitochondria are central hubs for cellular metabolism, coordinating ATP production, regulation of reactive oxygen species (ROS), and initiation of programmed cell death pathways. Disruption of mitochondrial function has been associated with altered antifungal susceptibility and changes in virulence-associated traits, such as morphogenesis and biofilm formation, suggesting that mitochondrial homeostasis contributes to the adaptive landscape of resistance (Shingu-Vazquez and Traven, 2011; Calderone and Clancy, 2014).

In addition, central carbon metabolism, including glycolysis and the tricarboxylic acid (TCA) cycle, has been implicated in modulating responses to azole stress. Metabolic flexibility may support cellular survival by maintaining energy homeostasis and mitigating oxidative stress during prolonged drug exposure. Resistant isolates often exhibit metabolic signatures distinct from susceptible strains, suggesting that reprogramming of core pathways complements canonical resistance mechanisms and sustains fungal fitness under drug pressure (Pereira et al., 2014; Akins et al., 2005).

Despite these insights, most mechanistic studies of azole resistance have focused on short-term drug exposure or laboratory-derived strains, leaving a gap in understanding how *C. albicans* adapts physiologically over decades of antifungal therapy. Clinical isolates from patients with chronic candidiasis and long-standing fluconazole treatment provide a unique opportunity to investigate sustained adaptations beyond classical resistance mechanisms. Importantly, antifungal therapy occurs in the context of ongoing host immune surveillance. While fluconazole imposes fungistatic stress in susceptible populations, its physiological impact in resistant backgrounds remains less clearly defined. In such contexts, drug exposure may not solely represent inhibitory pressure but could contribute to physiological remodelling of metabolic, mitochondrial, and cell-surface programs that influence fungal persistence during host interaction.

In this study, we investigated the fluconazole-resistant *C. albicans* isolate PUJ256 recovered from a patient with chronic mucocutaneous candidiasis after more than 30 years of continuous fluconazole therapy. Although the molecular mechanisms underlying azole resistance have been extensively characterized, much less is known about the physiological state adopted by resistant isolates after years or decades of continuous antifungal exposure. Through integrative phenotypic assays, quantitative proteomics, and functional analyses of mitochondrial activity and stress signalling, we show that long-term azole resistance is characterized by metabolic reprogramming and preserved mitochondrial integrity rather than exacerbated activation of classic stress pathways. These findings highlight adaptive strategies that support fungal persistence under sustained antifungal pressure and expand our understanding of resistance evolution beyond canonical mechanisms.

## Materials and Methods

### Cells and culture conditions

*Candida albicans* PUJ256, corresponding to isolate 4 from a clinical resistance isolate described by Ceballos-Garzón et al 2020, was used for the phenotypic and proteomic analysis. The control strain employed in this study was the wild-type *Candida albicans* strain SC5314. These strains were maintained on solid YPD medium (2% D-glucose, 2% peptone, 1% Yeast Extract and 2% agar) and incubated at 37 °C for one day. Before experiments, cells were grown on liquid-YPD medium at 30°C with rotatory shaking (180 rpm) until exponential phase was reached (optical density, 0.8 ± 0.1). Murine macrophages (RAW 264.7), derived from an Abelson murine leukemia virus induced tumor, were obtained from ATCC (TIB-71). Macrophages were maintained in RPMI-1640 medium (Sigma-Aldrich) supplemented with 10% fetal bovine serum (FBS) (Gibson) and 100 μg/ml penicillin/streptomycin and incubated at 37°C in a humidified atmosphere with 5% CO□.

### *ERG11* PCR and sequence analysis

The amplification of the ERG11 gene was carried out by amplifying two overlapping fragments: the first fragment covered positions 83 to 746 of the gene, and the second covered positions 639 to 1554. Primer sequences were obtained from Menéndez-Manjón et al., 2022. PCR amplifications were performed in a final reaction volume of 25 µL. Each reaction contained 1× MyTaq HS DNA polymerase (Bioline), forward and reverse primers (0.2 µl of final concentration), and 4 µL of DNA template. Thermal cycling conditions consisted of an initial denaturation step at 95 °C for 1 min, followed by 30 cycles of denaturation at 95 °C for 15 s, annealing at 57 °C for 30 s, and extension at 72 °C for 30 s. A final extension step was performed at 72 °C for 5 min. PCR products were purified using ExoSAP-IT™ PCR Product Cleanup Reagent (Thermo Fisher Scientific, USA), following the manufacturer’s instructions. Subsequently, 3 µL of each primer (forward and reverse) were added to separate tubes for sequencing reactions. Chromatograms were analyzed and consensus sequences were generated using SeqTrace software version 0.9.0 (Stucky, 2012). Consensus nucleotide sequences were translated into amino acid sequences and aligned with the reference ERG11 sequence of Candida albicans (Ca22chr5A_C_albicans_SC5314) obtained from the Candida Genome Database (Skrzypek MS et al., 2017) using MEGA version 12 (Tamura et al., 2021).

### Efflux of Rhodamine 6G

The assay was implemented according to a previously published protocol by Vargas-Casanova et al., 2024. Briefly, yeasts cells were initially grown in 10 mL of yeast peptone dextrose (YPD) broth at 35 °C for 20 under shaking (150 rpm). After, 2.5 mL of the suspension was added to 22.5 mL of PBS (1–5×10^7^ cells/mL) and incubated at 37 °C for 2 h. Subsequently, the cells were centrifuged (3000× g for 5 min) and washed three times with sterile distilled water. The resulting pellet was suspended in a solution of Rhodamine 6G (R6G) 83697 (Sigma-Aldrich St. Louis, Missouri, USA) (20 µM), followed by the addition of fluconazole at sub-inhibitory concentrations. The volume was adjusted to 25 mL with PBS. After incubation, cells were collected (3000×g, 5 min, 4 °C), washed twice with sterile distilled water, and resuspended in 25 mL of cold PBS (4 °C). For the efflux assay, 5 mL of the suspension was transferred to a new tube and supplemented with 500 µL glucose (final concentration: 2 mM). A negative control was prepared by replacing glucose with PBS. At 2 and 4 hours, 400 µL aliquots were collected and centrifuged (10,000×g, 1 min, 4 °C). Subsequently, 100 µL of the supernatant was transferred to a 96-well plate protected from light. Fluorescence intensity, corresponding to extracellular R6G, was measured using a Varioskan LUX spectrofluorometer (Thermo Scientific™, Waltham, MA, USA) at excitation and emission wavelengths of 530 and 560 nm, respectively. R6G concentrations were determined using a previously established calibration curve, enabling conversion of fluorescence intensity into concentration values. All experiments were performed in triplicate.

### Growth curves

For growth curves, 10 µL of *C. albicans* PUJ256 and SC5314 culture (OD_600_ = 0.04) were added to 180 µL of the YPD medium and YPD with fluconazole (1.1 μg/mL) and incubated for 24 h at 37 °C in SPECTROstar Nano (*BMG Labtech*).

### Fluconazole exposure and hyphal size measurement

Cells were prepared as suspensions with a density of 1 x 10^6^ cells/mL into plates with RPMI-1640 medium. *C. albicans* PUJ256 and SC5314 strains were incubated for 4 hours in RPMI-1640 medium with 1.1 μg/mL of fluconazole at 37°C. After that, a 1/1000 dilution was performed and 10 μL was inoculated on a YPD-agar plate. It was evenly spread and incubated for 24 hours at 37°C. Then, the CFU were counted. Images were also taken using the LasX microscope (*Leica Microsystems*), and 100 hyphae were measured under each condition with ImageJ (*National Institutes of Health*).

### In vitro competition experiments

A pre-culture of the strains was prepared. Cells were prepared as suspensions with a density of 5 x 10^5^ cells/mL into plates with YPD medium. *RFP*-labeled derivative of the wild type strain SC5314 and PUJ256 were mixed in a 1:1 ratio and were coincubated for 8 hours at 30 °C. After that, a 1/1000 dilution was performed and 10 μL was inoculated on a YPD-agar plate. It was evenly spread and incubated for 24 hours at 37°C. The percentage of red colonies (*RFP*-labelled SC5314) and white colonies (PUJ256) was determined after storage of the plates at 4 °C for four days, which increased the fluorescence intensity of the *RFP*-expressing colonies.

### Filamentation assays

A pre-culture of the strains was prepared. Cells were then collected by 10 min of centrifugation at 3,500 rpm in an-Eppendorf 5810R centrifuge, the supernatant discarded, and cells washed with PBS and collected again by 3 min of centrifugation at 5,000 rpm in a Heraeus Fresco 21 microcentrifuge (Thermo Scientific). The pellet was resuspended in milliQ water. The suspension was diluted to reach a final absorbance of 0.4. Drop growth assay was then performed by placing 4 μL drops on YPD-agar, YPD-agar + 10% Fetal Bovine Serum (FBS), Spider (1% mannitol, 1% nutrient broth, 0.2% K_2_HPO_4_, and 2% agar, pH 7.2), Spider + fluconazole 1.1 μg/mL, Sabouraud (4% dextrose, 1% peptone, and 2% agar, pH 5.6), Sabouraud + 0.04% SDS, and Sabouraud + 0.04% SDS + fluconazole 0.55 μg/mL. The plates were incubated at 37°C for 24 hours and then at 4°C for 72 hours.

### Chitin content

A pre-culture of the strains was prepared. Cells were washed with PBS and collected again by 3 min of centrifugation at 5,000 rpm in a Heraeus Fresco 21 microcentrifuge (Thermo Scientific). The pellet was resuspended in milliQ water. Cells were stained with 25 μg ml^−1^ calcofluor white M2R (CFW) (Sigma-Aldrich) for 10 minutes at 4 °C. All samples were examined by fluorescence microscopy using a LasX microscope (Leica Microsystems). Chitin content was estimate by measuring CFW fluorescence with FLUOstar Galaxy (*BMG Labtech*).

### Membrane permeabilization assays

A pre-culture of the strains was prepared. To observe the effect of SDS in liquid medium and measure the percentage of cell death (propidium iodide (PI) positive), strains were resuspended to reach an absorbance of 0.4 and incubated for 24 hours at 37°C in Sabouraud medium and Sabouraud + 0.04% SDS. Viability was assessed by PI staining (1.25 µg/mL). The percentage of dead cells was determined by counting 100 cells. Images were captured using a LasX microscope (Leica Microsystems).

### Biofilm formation assay

Biofilm formation capacity was determined based on a protocol described by Fattouh et al. Briefly, a single colony from each isolate was cultured overnight in 5 mL of potato dextrose broth (PDB) at 30 °C with shaking at 100 rpm. A suspension containing 10 *C. albicans* cells/mL in a total volume of 200 µL was used to inoculate wells of a 96-well plate that had been pre-treated overnight with 5% fetal bovine serum at 4 °C. The plate was incubated for 3 h at 37 °C with shaking at 75 rpm. Following incubation, the wells were washed once with 1× phosphate-buffered saline (PBS) and 0.2 mL of fresh potato dextrose broth was added. Plates were then incubated for 48 h at 37 °C with shaking at 75 rpm of PDB, PDB + fluconazole (1.1 µg/mL), YPD and YPD + fluconazole (1.1 µg/mL). Wells were washed once with 1× PBS to remove planktonic cells. Biofilms were fixed by adding 0.2 mL of 99% methanol for 15 min, after which methanol was removed and wells were air-dried for 20 min. Biofilms were then stained with 0.2 mL of 0.2% crystal violet for 20 min, followed by five washes with sterile water. To solubilize the dye, 0.2 mL of 33% acetic acid was added, and optical density was measured at 595 nm using the SPECTROstar Nano (*BMG Labtech*).

### *Candida albicans*–RAW264.7 macrophage interaction

The culture conditions for both yeast strains and macrophages were maintained as previously described. Yeast and macrophages were prepared as suspensions at a density of 1 × 10 cells/mL in RPMI-1640 medium (Sigma-Aldrich) and seeded in plates at a multiplicity of infection (MOI) of 1:1. The strains were co-incubated with RAW 264.7 macrophages for 4 hours at 37 °C in a humidified 5% CO atmosphere, in the presence or absence of fluconazole (1.1 μg/mL). As controls, yeasts were incubated under the same conditions but without macrophages. To assess fungicidal activity, 1:100 dilutions were performed and 100 μL aliquots were plated onto YPD-agar, followed by incubation at 37 °C for 24 hours to determine CFUs. To evaluate phagocytic activity, a differential staining protocol was established to quantify the phagocytic process. *C. albicans* cells were pre-labeled with Oregon Green 488 (1 μM) (Molecular Probes) for 20 min at 30 °C in darkness with gentle agitation. After washing with PBS-glycine (100 mM), yeast cells were resuspended at the desired density and co-cultured with RAW264.7 macrophages (1 × 10 cells/mL) at a MOI of 1:1. Cells were then washed with cold PBS, fixed with 4% paraformaldehyde for 30 min, and counterstained with calcofluor white M2R (2.5 μM) (Sigma-Aldrich) for 10 minutes to differentiate internalized yeast from adherent ones. Images were acquired using a LasX microscope (*Leica Microsystems*) equipped with FITC (excitation/emission 480/535) and UV filters (excitation/emission 365/397), and 600 cells per condition were counted using ImageJ (National Institutes of Health). The phagocytosis percentage was calculated as follows: % phagocytic cells = (calcofluor white-stained cells / Oregon Green-stained cells) × 100.

### Cell disruption and protein extract quantification

Cells were prepared as suspensions with a density of 1 x 10^6^ cells/mL into plates with RPMI-1640 medium. *C. albicans* PUJ256 and SC5314 strains were incubated for 4 hours in RPMI-1640 medium with 1.1 μg/mL of fluconazole at 37°C. Control and treated cells were washed three times with PBS. The cell pellet was resuspended in lysis buffer (50 mM Tris-HCl [pH 7.5], 1 mM EDTA, 1 mM DTT, 150 mM NaCl, 10% protease inhibitors [Thermo Scientific, Waltham, MA, USA], and 5 mM PMSF). Cell extracts were obtained by mechanical disruption using a Fast-Prep system (Bio101, Savant, Thermo Fisher, Waltham, MA, USA), with 5 cycles of 30 seconds, applying glass beads (0.5 to 0.75 mm diameter). The samples were then centrifuged for 20 minutes at 13,000 rpm to separate the protein extracts from cell debris, and protein concentrations were measured using a Bradford assay.

### LC-MS/MS

After cell lysis, peptide digestion was performed using 50 micrograms of protein extracts (iST kit, PREOMICS). The samples were denatured, reduced, and alkylated; subsequently, they were digested using a trypsin/LysC mix, and the resulting peptides were purified using a reverse-phase LC-MS column. The final peptide concentration in the samples was quantified by fluorometry using a Qubit4 system (Thermo Scientific). Finally, 0.4 μg peptides were loaded for mass spectrometric analysis on Evosep One (Evosep) coupled to a Tims TOF Pro 2 (Bruker). Peptides were separated on a C18 resin analytical column Bruker Daltonics performance (PepSep C18, 15cm x 150 µm ID, Bruker) using an 88 min gradient. Data were obtained by data-dependent acquisition in positive mode. Each MS scan (350 - 1700 Da) was divided in 10 ramps of ion mobility (between 0.6 and 1,6 V·s/cm2), and in each ramp, the 10 most intense precursors (charges 2-5) were selected for their high collision energy dissociation fragmentation.

### Protein quantification

Raw data were processed using FragPipe v22 pipeline. The database selected was the Candida Genome Database Assembly 21 (A21-s02-m09-r12)) with 6279 sequences. Quantitation was performed using the default LFQ-MBR settings with slight modifications. Precursor mass tolerance was set to-20, 20 ppm and fragment mass tolerance was set to 20 ppm, in silico digestion was performed with strict trypsin allowing for 2 missed cleavages and peptide lengths of 5-40 aminoacids, and peptide mass range set to 700-5,000 kDa. Variable modifications searched included methionine oxidation and N-terminal methionine loss. Cysteine carbamidomethylation was set as a static modification. Peptide and protein identifications were validated using Philosopher with Percolator and ProteinProphet, applying a 1% false discovery rate (FDR). Finally, the quantification module was performed using IonQuant with Match between runs option selected. Finally, results were normalized to the total peptide abundance to equalize signal intensity across samples. The mass spectrometry proteomics data have been deposited to the ProteomeXchange Consortium via the PRIDE partner repository (Perez-Riverol et al., 2025) with the dataset identifier PXD077083.

### GO enrichment analysis and protein clustering

GO enrichment analysis was performed using the GO Term Finder and GO Slim Mapper tools from the Candida Genome Database (CGD), based on biological process, molecular function, and cellular component. Protein network clustering was conducted using STRING v.12.0 software. The obtained proteins were classified according to Uniprot. For data integration, the FungiDB and KEGG databases were used.

### Mitochondrial function

Mitochondrial membrane potential measurement was used to determine changes in mitochondrial activity of the PUJ256 and SC5314 strains after treatment with fluconazole (1.1 μg/mL) for 4 hours. Cells were washed, resuspended in PBS, and the fluorescent reagent JC-1 (Invitrogen®) was added at a final concentration of 0.5 μL/mL, followed by incubation for 20 minutes. After washing, cell fluorescence was analyzed using a confocal laser microscope (FluoView-1200, Olympus) with ImageJ software (National Institutes of Health).

### MAPK detection

To validate the proteomic data, immunodetection of the phosphorylated MAPK proteins p-Hog1, p-Cek1, and p-Mkc1 was performed by Western blot. Tunicamycin (2.5 μg/mL) was used as a positive control for 2 hours. Samples (100 μg of protein per sample) were heated at 99 °C for 5 minutes with loading buffer. Protein separation was performed by SDS-PAGE electrophoresis for 1 hour at 150 V using 10% gels. Proteins were electrotransferred to a nitrocellulose membrane (GE Healthcare) for 60 minutes at 100 V. Membranes were washed with distilled water and stained with 15 mL of Ponceau Red (Pierce™ 24580) (0.1 g/mL) for 5 minutes. Blots were blocked with 10% powdered milk in PBS with 0.1% Tween 20 (PBS-T) for 120 minutes. The presence of the target proteins was detected using specific primary antibodies: anti-p-Hog1 (1:500, BioRad) and anti-p-Mkc1 (1:1000, BioRad) in 5% powdered milk in PBS-T overnight at 4 °C. After washing with PBS-T, blots were incubated with Alexa Fluor Plus 800 secondary antibody diluted in 5% powdered milk in PBS-T for 1 hour in the dark. Signals were detected using the ChemiDoc MP imaging system (BioRad). Bands were densitometrically quantified using ImageJ (National Institutes of Health). Total protein loading control was verified by Ponceau Red staining.

## Statistical analysis

Graphs and statistical analyses were performed using GraphPad Prism 8. Data represent the mean ± standard deviation of at least three independent biological replicates for both strains under control and fluconazole-treated conditions. Statistical significance was assessed using Student’s *t*-test or one-way ANOVA, as appropriate. *P* values < 0.05 were considered statistically significant (*p* < 0.05, p < 0.01, *p* < 0.001, p < 0.0001).

## Results

### Genetic characterization of the classical azole resistance mechanism mediated by *ERG11*

As a first step towards understanding the molecular basis of azole resistance in the clinical isolate PUJ256, the coding sequence of *ERG11*, which encodes the target enzyme of azole antifungals (lanosterol 14α-demethylase), was analysed and compared with that of the reference strain SC5314. Five amino acid substitutions were identified in PUJ256 (S263L, E266D, V402F, S405F and V488I), compared to the susceptible reference strain SC5314 (**Table 1**). Among these substitutions, only S405F has been experimentally demonstrated to reduce azole susceptibility when introduced into a susceptible genetic background. In contrast, E266D and V488I are commonly reported polymorphisms that are also found in azole-susceptible isolates, while the contribution of S263L and V402F to resistance remains unclear. (**Table 1**; **Figure 1**).

**Figure 1.**
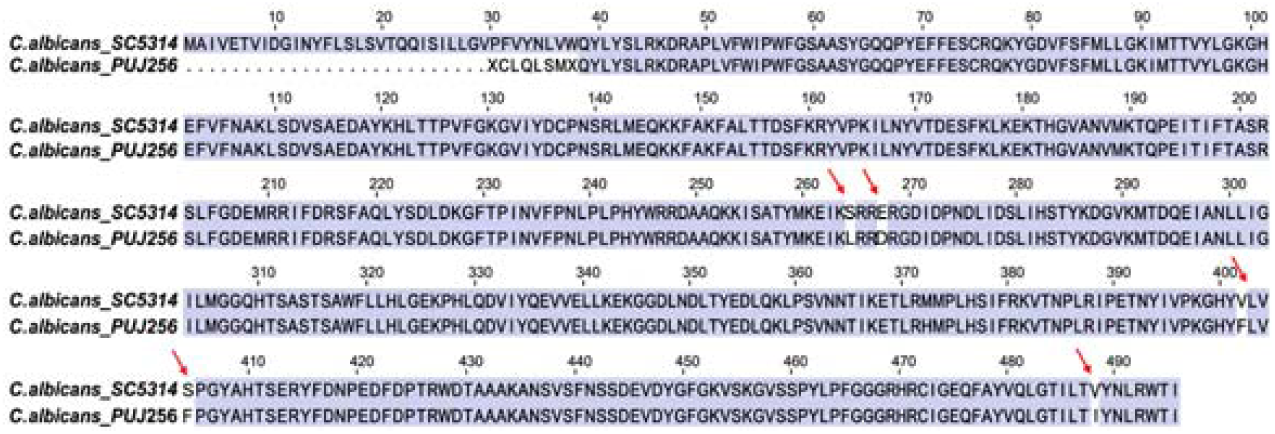
Alignment of *C. albicans* SC5314 and *C. albicans* PUJ256. Amino acid differences between the strains are displayed without colour.

**Table 1.** Mutations found in the strain *C. albicans* PUJ256.

| Position in *ERG11* | Reference | Mutation |
| --- | --- | --- |
| 263 | S | L |
| 266 | E | D |
| 402 | V | F |
| 405 | S | F |
| 488 | V | I |

### Functional analysis of efflux activity

To determine whether active drug efflux contributes to the resistant phenotype of PUJ256, Rhodamine 6G (R6G) extrusion assays were performed following exposure to subinhibitory concentrations of fluconazole. Because R6G is actively exported by ATP-dependent ABC transporters, extracellular fluorescence was used as a functional indicator of efflux activity.

In the susceptible strain SC5314, fluconazole pretreatment did not stimulate R6G extrusion at either 2 or 4 h, with extracellular R6G levels remaining similar to or slightly lower than those observed under untreated conditions (**Figure 2**). In contrast, PUJ256 exhibited a significant increase in R6G efflux following fluconazole exposure at both time points (P < 0.0001), with the highest extrusion observed after 4 h of treatment. These results indicate that fluconazole induces a sustained efflux response in the resistant isolate but not in the susceptible reference strain.

**Figure 2.**
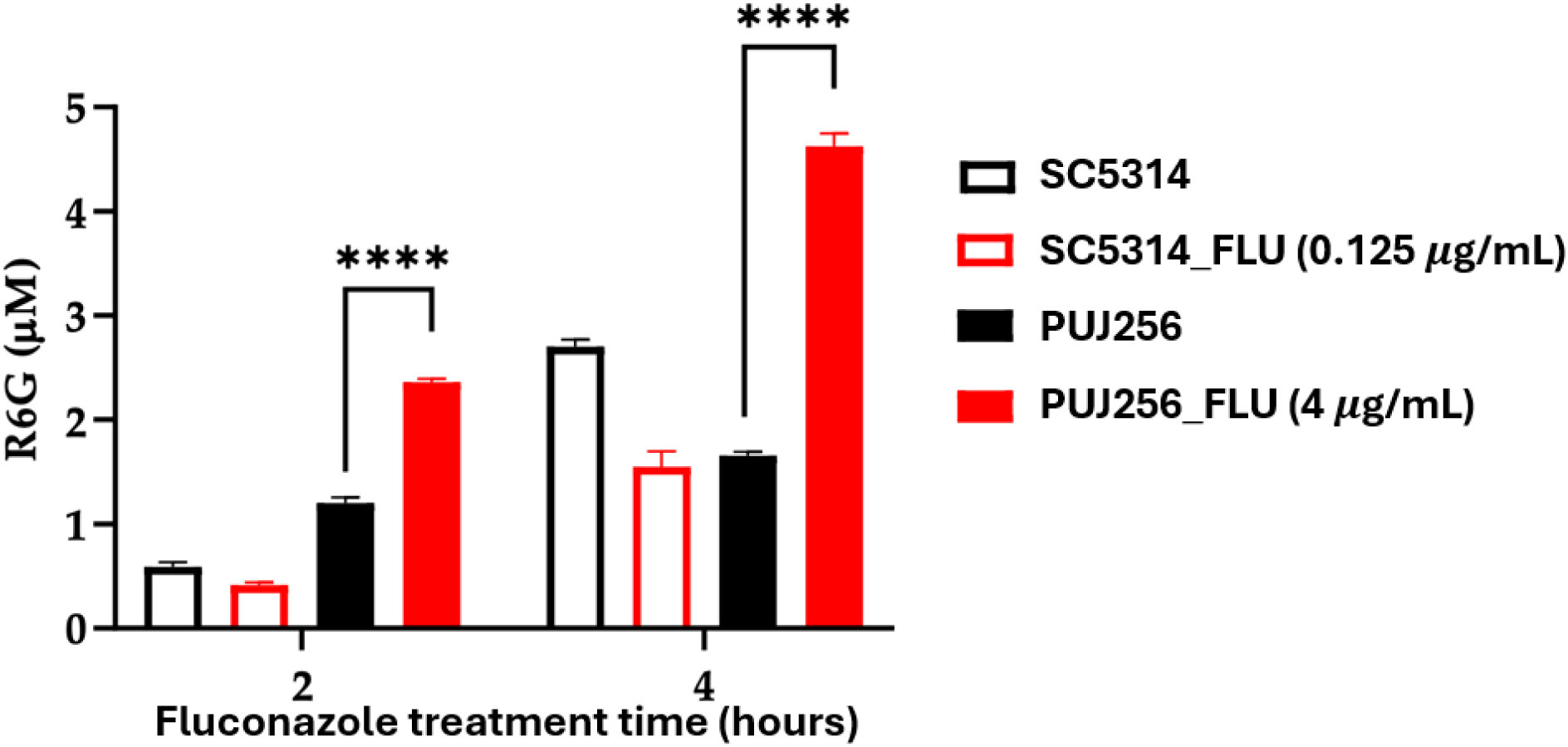
Functional analysis of efflux activity in *Candida albicans* SC5314 and the azole-resistant clinical isolate PUJ256. Cells were exposed to subinhibitory concentrations of fluconazole (0.125× MIC) for 2 or 4 h. Following loading with rhodamine 6G (R6G) and metabolic reactivation with glucose, extracellular R6G concentration was determined as a functional measure of ATP-dependent efflux activity. Data are presented as mean ± SD from three independent experiments. Statistical significance was assessed using two-way ANOVA followed by Šídák’s multiple comparisons test. **P < 0.0001.

### Growth, viability and fitness of the azole-resistant isolate PUJ256 compared with the reference strain SC5314

To comprehensively evaluate the biological cost associated with long-term azole resistance, we assessed the growth, viability and fitness of PUJ256 under both basal and fluconazole-treated conditions. Here, fitness was defined as the ability of each strain to maintain growth, viability and competitive capacity under fluconazole pressure, as assessed by growth kinetics, viability assays and mixed-strain competition experiments. Growth kinetics were evaluated for both *C. albicans* strains by monitoring optical density at 600 nm over a 24-h period in YPD medium. Experiments were conducted using a fluconazole concentration of 1.1 μg/mL, which was predetermined to induce significant growth inhibition without achieving full fungicidal activity in SC5314 (**Figure 3A**). Complementary cell viability assays in RPMI-1640 medium supported these findings; the reference strain exhibited a substantial reduction in viability, reaching approximately 52.2% of the untreated control after 4 hours of exposure (**Figure 3B**). In contrast, no significant changes in viability were detected for the PUJ256 strain under the same conditions, consistent with its fluconazole-resistant phenotype. Competition assays in YPD medium between the azole-sensitive strain SC5314 and the resistant strain PUJ256 revealed a treatment-dependent fitness shift affecting population dominance over an 8-hour period (**Figure 3C**). Under control conditions, SC5314 maintained a competitive advantage, accounting for approximately 75% of the total population. Exposure to 0.22 μg/mL of fluconazole inverted this ratio, with the resistant strain PUJ256 increasing to ∼68% of the total CFUs. At a higher concentration of 1.1 μg/mL, the resistant strain PUJ256 outcompeted SC5314 (**Figure 3C**) demonstrating that fluconazole exposure shifts the competitive balance in favour of the resistant isolate.

**Figure 3.**
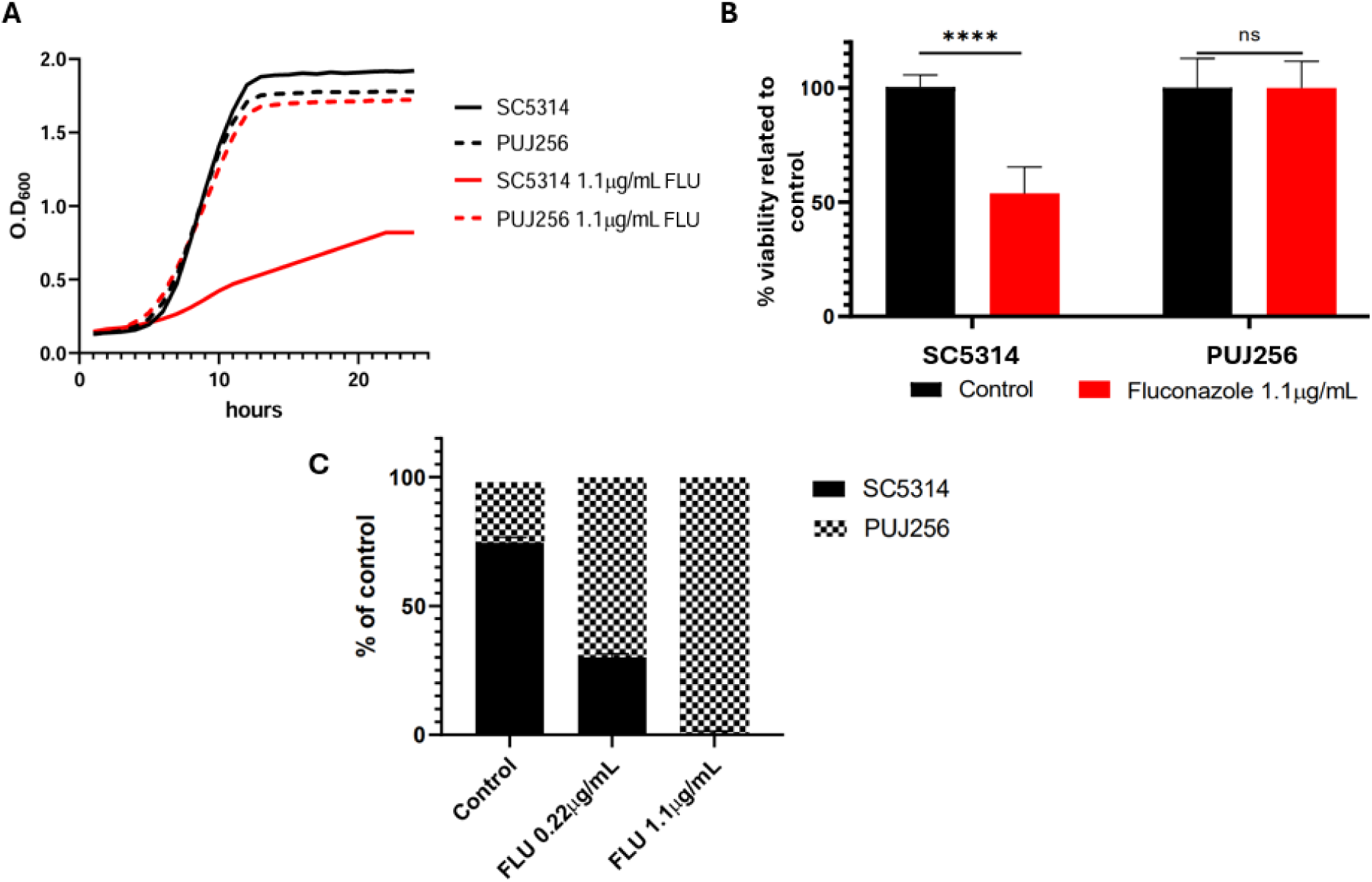
Fluconazole susceptibility profile and competitive fitness of *C. albicans* clinical isolates SC5314 and PUJ256. **(A)** Representative growth curves in YPD medium. SC5314 (solid black line) and PUJ256 (dashed black line) exhibit similar growth kinetics in the absence of treatment (Control). Treatment with 1.1 μg/mL of fluconazole (red lines) results in substantial growth inhibition of SC5314 (solid red line) but not PUJ256 (dashed red line). **(B)** Relative viability after fluconazole exposure (1.1 µg/mL), expressed as percentage of untreated control, after.4 hours at 37 °C in RPMI-1640 medium. **(C)** Competitive fitness of SC5314 and PUJ256 under fluconazole pressure for 8 hours at 30 °C in YPD medium. Data represent mean ± SD, ANOVA *test*, n =3. ns: not significant; FLU: fluconazole.

### Filamentation and plate invasion ability of PUJ256 compared with SC5314

The comparison of the morphological transition ability between strains was assessed. The clinical isolate PUJ256 exhibited a significantly smaller hyphal size after 4 hours of incubation at 37 °C in RPMI-1640 medium compared to SC5314 (**Figure 4A**). Upon fluconazole exposure (1.1 μg/mL) SC5314 showed 78% reduction marked reduction in hyphal length, decreasing from 75.5 6.7 μm to 16.6 3.1 μm. In contrast, PUJ256 was minimally affected by the antifungal (7% reduction), from 27.9 4.4 μm to 25.9 6.5 μm (**Figure 4B**). These results indicate that although PUJ256 displays impaired filamentation under basal conditions, its morphology is largely insensitive to fluconazole, whereas SC5314 undergoes a drastic filamentation defect upon treatment. To evaluate filamentation and agar invasion, a spot growth assay was performed on different solid media that promote these processes (YPD, YPD supplemented with 10% fetal bovine serum (FBS), Spider, and Spider supplemented with fluconazole (1.1 μg/mL)). On both YPD and YPD supplemented with 10% FBS, SC5314 displayed robust filamentation, resulting in characteristic irregular and rough colonies (**Figure 4C**). Conversely, PUJ256 failed to filament under these conditions, maintaining smooth, creamy colonies typical of yeast-phase growth (**Figure 4C**). On Spider medium, SC5314 underwent filamentation, whereas PUJ256 did not (**Figure 4C-D**). Notably, on Spider medium supplemented with fluconazole (1.1 μg/mL), the invasive growth of SC5314 was completely abrogated, whereas PUJ256 displayed growth and colony morphology comparable to its performance on drug-free medium, highlighting its ability to maintain its fitness despite antifungal pressure (**Figure 4C**).

**Figure 4.**
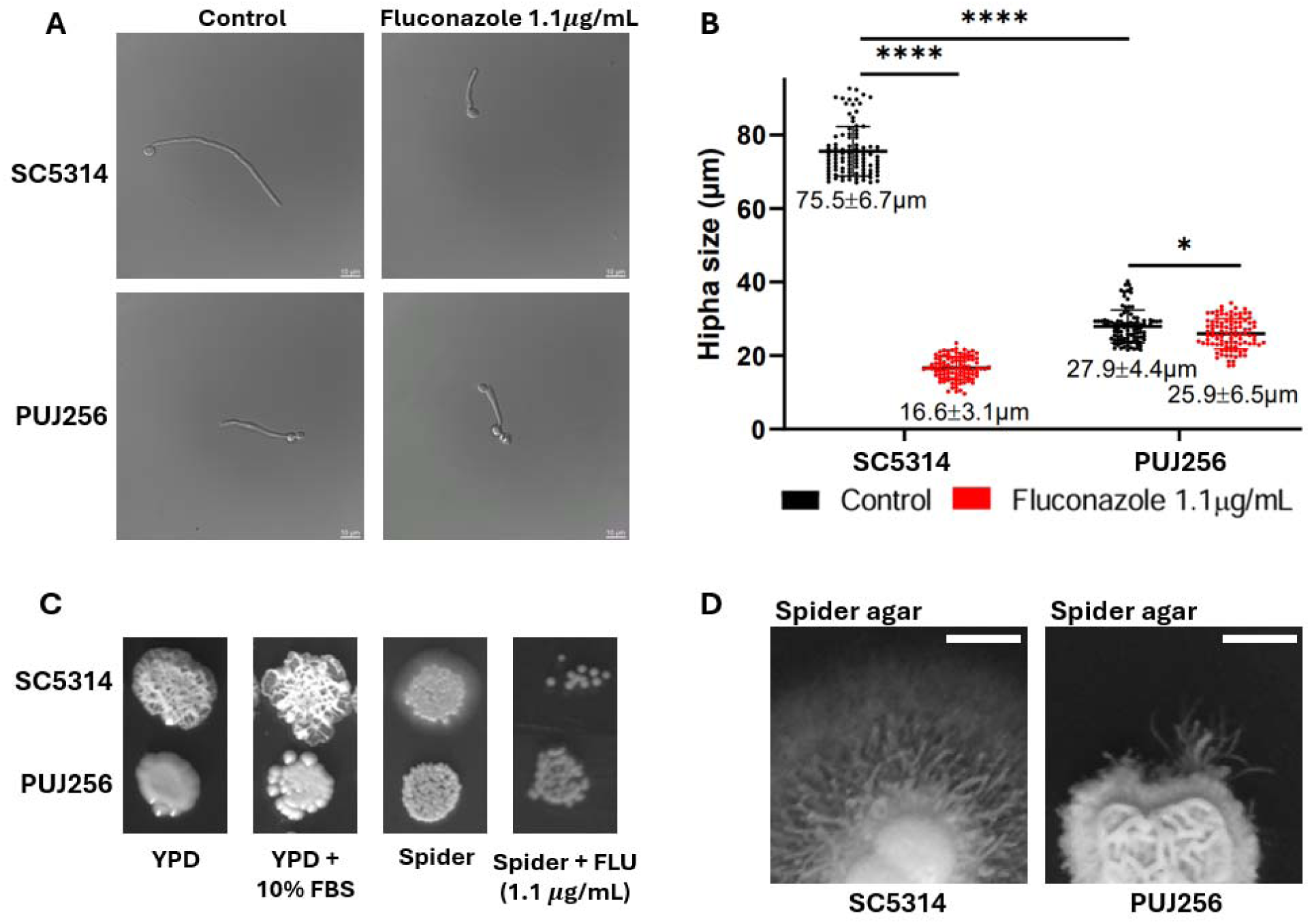
Filamentation and plate invasion capacity of PUJ256 compared with SC5314. **(A)** Bright-field microscopy of SC5314 and PUJ256 in RPMI-1640 medium. Both strains were incubated for 4 hours at 37°C to induce filamentation in the absence (Control) or presence of 1.1 µg/mL of fluconazole Scale bar: 10 µm. **(B)** Comparison of hyphal length in SC5314 and PUJ256 in the presence and absence of fluconazole. (1.1 µg/mL). Data represent mean ± SD, ANOVA *test*, n = 100. **(C)** Spot growth assay of SC5314 and PUJ256 on YPD, YPD + 10% FBS, Spider medium and Spider medium + fluconazole (1.1 µg/mL). **(D)** Colony morphology of SC5314 and PUJ256 grown on Spider medium showing agar invasion. Scale bars represent 1 cm.

To further characterize the structural adaptations in the clinical isolate, we quantified chitin content using Calcofluor White (CFW) staining followed by fluorescence intensity measurement (FIM) and microscopy (**Figure 5**). Quantitative analysis revealed that PUJ256 possesses a significantly higher basal chitin content compared to the SC5314 reference strain (**Figure 5A**). Fluorescence microscopy confirmed these findings, showing a more intense and robust blue signal along the cell periphery and at the bud scars of PUJ256 yeast cells relative to SC5314 (**Figure 5B**). This enrichment in chitin suggests a constitutive remodelling of the cell wall architecture in the long-term evolved resistant isolate, a feature frequently associated with increased physical robustness and altered recognition by immune cells.

**Figure 5.**
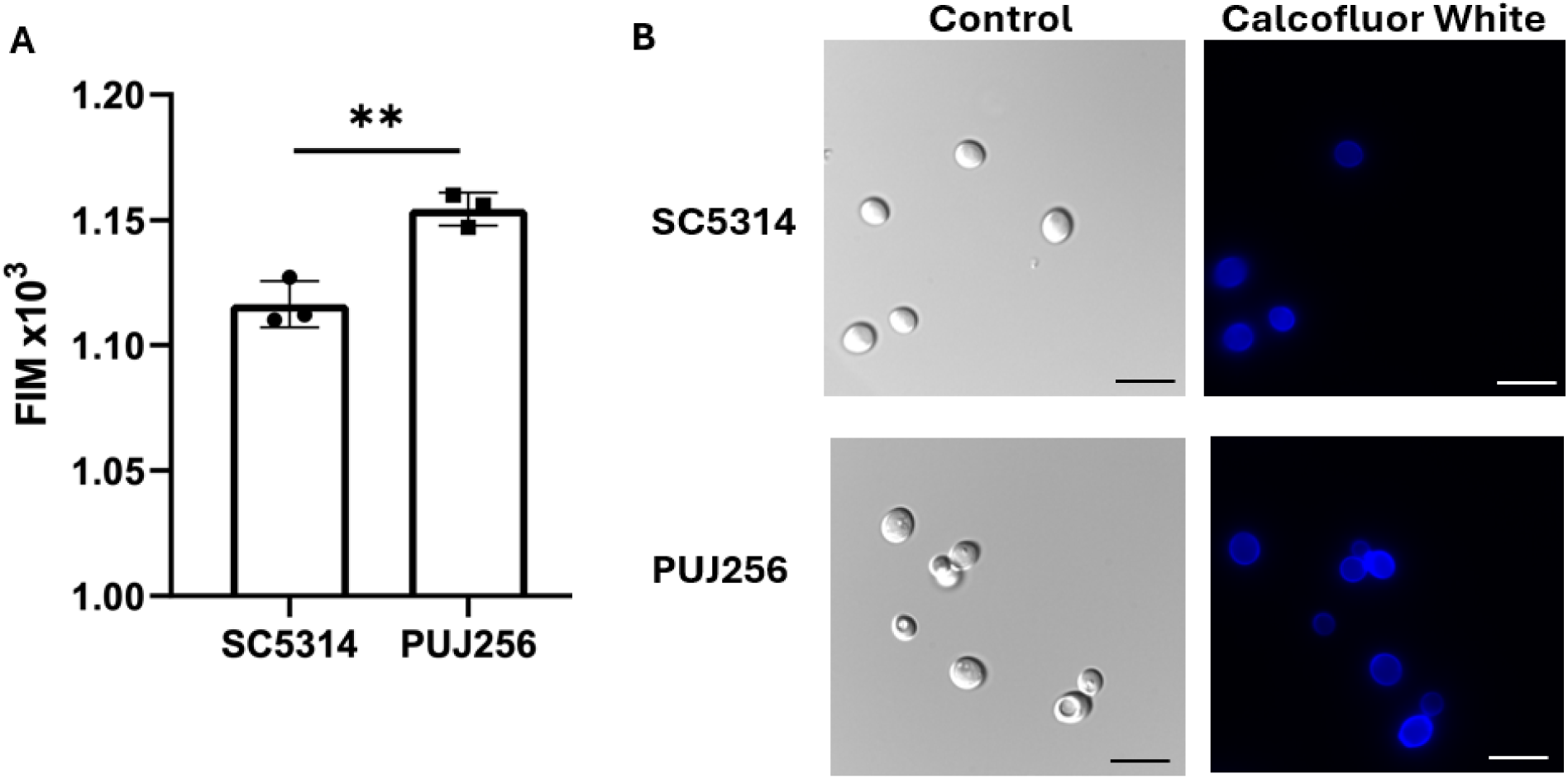
Comparative analysis of cell wall chitin content between *C. albicans* SC5314 and PUJ256. **(A)** Quantitative assessment of chitin levels represented as Fluorescence Intensity Measurement (FIM x 10^3^). Data represent mean ± SD, Student’s t-test, n = 100 cells. **(B)** Representative bright-field (left) and fluorescence (right) microscopy images of yeast cells stained with Calcofluor White (CFW). Scale bars: 10 μm.

### Membrane integrity of PUJ256 compared with SC5314

To assess plasma membrane integrity cells were exposed to the detergent SDS at 0.04%, a known membrane-disrupting agent, for 24 hours at 30 °C. Growth was assessed by spot assays (**Figure 6A**) and cell viability was quantified by propidium iodide (PI) staining (**Figure 6B**). The resistant isolate showed markedly increased sensitivity to SDS compared to SC5314, with impaired growth and PI viability (∼57% at 0.04% SDS) indicating compromised membrane integrity. In contrast, SC5314 at this SDS concentration maintained higher viability and growth. Neither strain grew in Sabouraud medium supplemented with 0.04% SDS and fluconazole (1.1 µg/mL), likely reflecting excessive combined cellular stress (**Figure 6C-D**).

**Figure 6.**
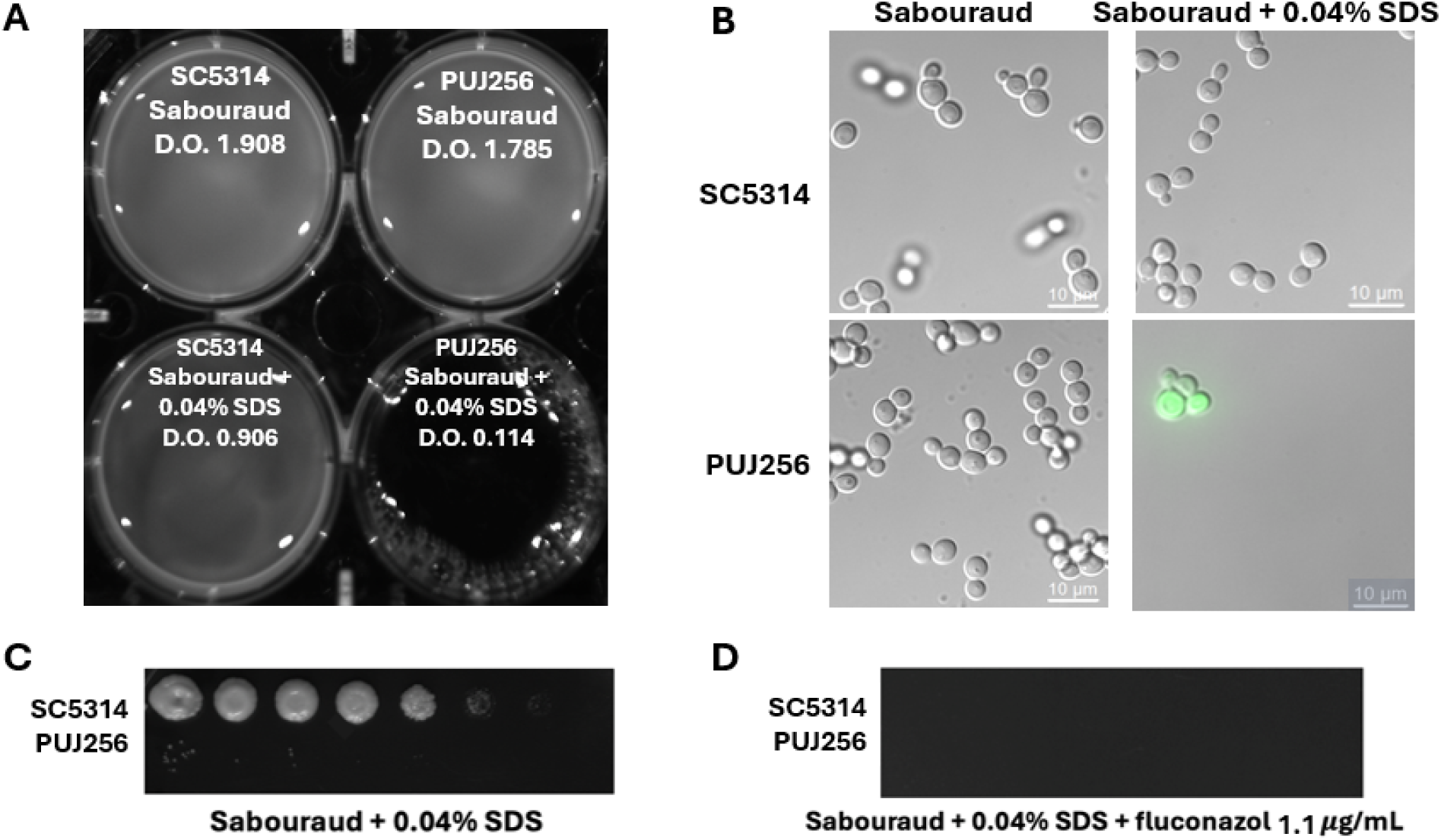
Effect of SDS detergent on *C. albicans* SC5314 and PUJ256. **(A)** Photograph of the wells corresponding to each strain and condition, along with their optical density values after 24 hours of incubation at 30°C. **(B)** Bright-field microscopy images of both strains under control and SDS-treated conditions. **(C)** Spot assay of both strains on Sabouraud-agar supplemented with 0.04% SDS and **(D)** Sabouraud-agar supplemented with 0.04% SDS + fluconazole (0.55 µg/mL). Scale bar: 10 µm.

### Comparative interaction of SC5314 and PUJ256 with RAW264.7 macrophages

We evaluated both, RAW264.7 macrophage phagocytic and fungicidal activity against both strains after 4 hours. RAW264.7 macrophages exhibited similar killing efficiency against both strains under basal conditions in RPMI-1640 medium (**Figure 7A**). However, in the presence of fluconazole (1.1 μg/mL), PUJ256 exhibited significantly higher survival following exposure to the candidacidal activity of RAW264.7 macrophages than SC5314, demonstrating increased resistance of this isolate under antifungal stress despite its lower basal filamentation capacity (**Figure 7A**). Regarding phagocytosis assays, PUJ256 was more efficiently internalized than SC5314 in the absence of the drug (**Figure 7B**), consistent with its predominantly yeast morphology. Nevertheless, fluconazole treatment reversed this trend, causing a significant decrease in the percentage of phagocytosed PUJ256 cells compared to its untreated control, whereas the drug increased the internalization rate in SC5314, findings that were confirmed by light microscopy analysis (**Figure 7C**).

**Figure 7.**
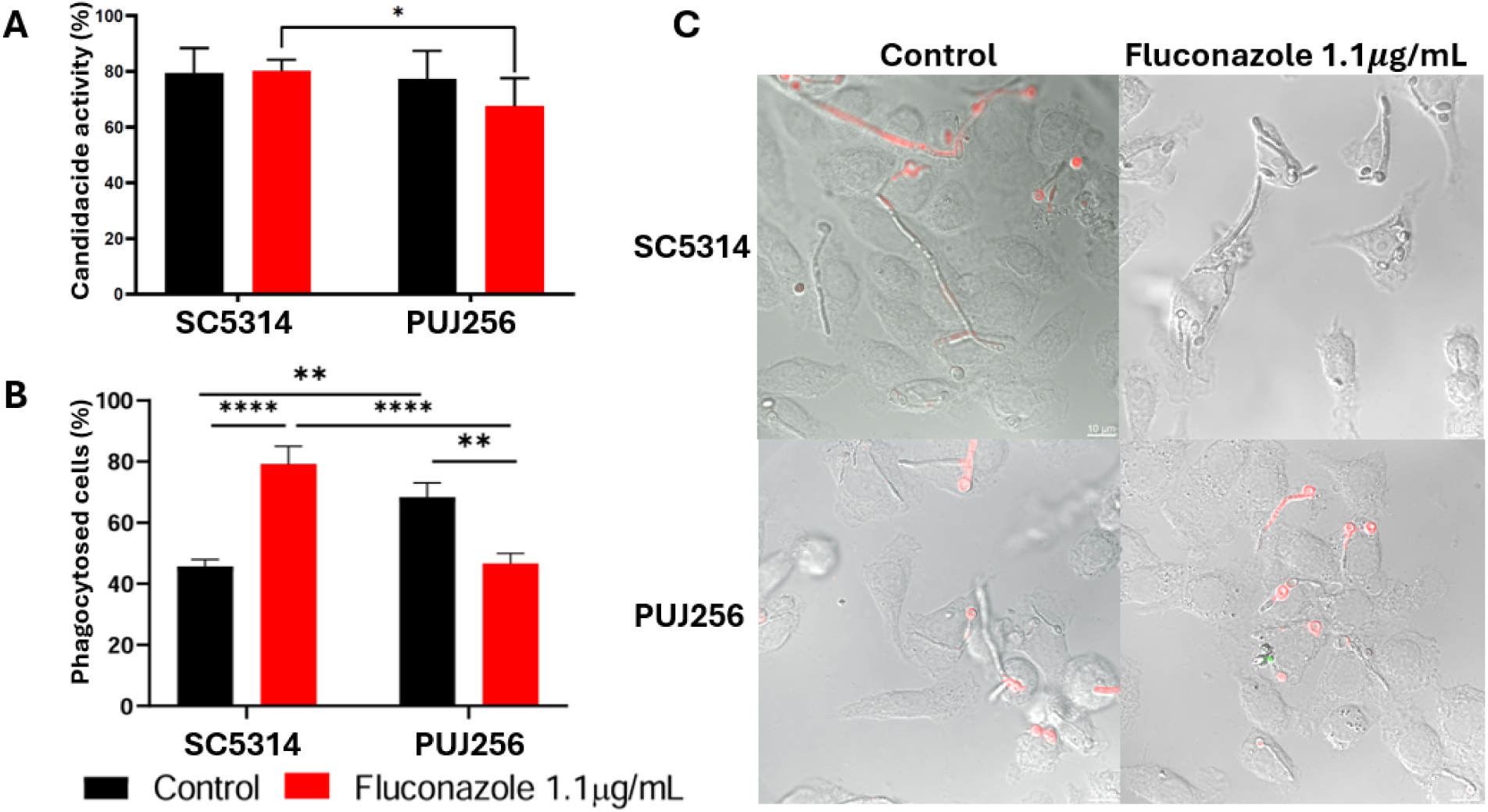
Susceptibility of SC5314 and PUJ256 to RAW 264.7 macrophages killing and phagocytosis in the presence and absence of fluconazole (1.1 µg/mL). **(A)** Percentage of macrophage candidacidal activity after 4 hours of interaction with SC5314 and PUJ256 under control conditions or in the presence of fluconazole (1. 1 µg/mL). **(B)** Percentage of phagocytosed cells after 4 hours of interaction under the same conditions. **(C)** Representative images of RAW 264.7 macrophages interacting with both strains under the same conditions. Data represent mean ± SD, ANOVA *test*, n = 3.

### Biofilm formation in PUJ256 compared with SC5314

To determine whether the fluconazole-associated resistance phenotypes observed in planktonic cells and host-interactions extended to surface-associated persistence traits, we assessed the biofilm formation potential of both *C. albicans* strains under nutrient-poor (PDB) and nutrient-rich (YPD) conditions (**Figure 8**). Quantitative analysis of biofilm biomass via crystal violet staining revealed that in PDB there were no statistically significant differences in biofilm formation capacity between the SC5314 and PUJ256 strains (**Figure 8A**). However, the addition of fluconazole (1.1 μg/mL) to PDB medium induced a dramatic, significant increase in biofilm production in PUJ256, while having no effect on SC5314 (**Figure 8A**). This response was also observed under nutrient-rich conditions in YPD medium, where SC5314 formed significantly more robust biofilms than PUJ256 (**Figure 8A**) with microscopic images in **Figure 8B** showing a more intricate, hyphal-rich structure for SC5314. Yet, upon fluconazole exposure, PUJ256 showed a statistically significant elevation in biofilm biomass in YPD, whereas no significant change was detected for SC5314 in this condition (**Figure 8A**).

**Figure 8.**
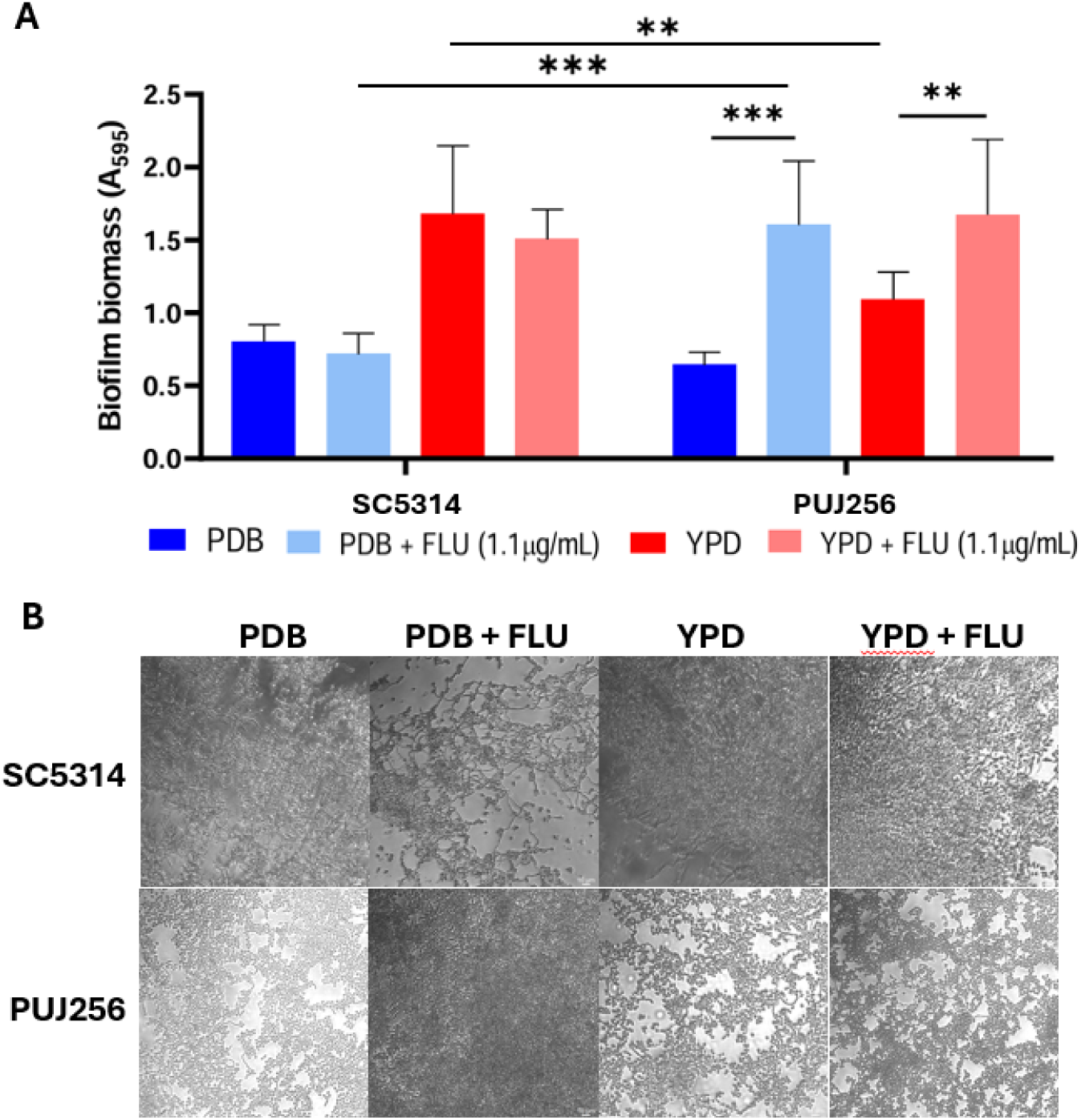
Biofilm formation of SC5314 and PUJ256 under different growth conditions and fluconazole exposure. **(A)** Quantification of biofilm biomass measured as optical density at 595 nm (OD_595_) for all isolates grown in PDB or YPD, with or without fluconazole (FLU, 1.1 µg/mL). Data represent mean ± SD, ANOVA, n = 3. **(B)** Representative phase-contrast microscopy images of biofilms formed by SC5314 and PUJ256 under the same conditions. Scale bars: 10 µm.

### Comparative proteomic analysis of PUJ256 and SC5314 in the presence and absence of fluconazole

To gain insight into the proteomic changes associated with fluconazole resistance, we compared the protein expression profiles of the azole-resistant isolate PUJ256 and SC5314 under fluconazole-treated and untreated conditions. A label-free proteomic approach was used to analyse total proteins from cytoplasmic extracts of both strains grown for 4 hours in RPMI-1640 medium supplemented with 10% FBS and 100 μg/mL penicillin/streptomycin, in the presence or absence of fluconazole (1.1 μg/mL). Following protein identification by mass spectrometry, the dataset was filtered to include only proteins detected in at least three out of four biological replicates and identified with a minimum of two unique peptides. Relative protein abundance between strains and conditions was then assessed, generating abundance ratios and a list of quantified proteins meeting reliable statistical criteria based on q-values. **Table 2** summarizes the total number of proteins quantified in the different pairwise comparisons, including comparisons between strains, within each strain with and without fluconazole, and between strains under fluconazole treatment. The number of proteins showing significant positive or negative log fold-change values is indicated for each comparison.

**Table 2.** Number of quantified proteins and proteins showing significant differences in abundance (positive or negative log_2_FC; p<0.05) across the four pairwise comparisons. FLU: Fluconazole. The number of proteins exclusively identified in each condition appears in parentheses.

| Comparative | Total quantified proteins | Total differentially abundant proteins | Increased (log<sub>2</sub>FC>0) | Decreased (log<sub>2</sub>FC<0) |
| --- | --- | --- | --- | --- |
| PUJ256 / SC5314 | 1896 | 461 | 298 (196) | 163 (95) |
| PUJ256_FLU / SC5314_FLU | 1860 | 514 | 253 (99) | 261 (135) |
| SC5314_FLU / SC5314 | 1774 | 352 | 218 (132) | 134 (83) |
| PUJ256_FLU / PUJ256 | 1874 | 257 | 86 (68) | 171 (155) |

Across all pairwise comparisons, a similar number of proteins was identified, ranging from approximately 1,700 to 1,900 quantified proteins (**Table 2**). Under basal conditions, 461 proteins showed significant differences in abundance between PUJ256 and SC5314 (298 increased and 163 decreased in PUJ256). Following fluconazole exposure, the proteomic divergence between the two strains increased to 514 differentially abundant proteins (253 increased and 261 decreased in PUJ256). Fluconazole treatment induced more extensive proteome remodelling in SC5314 (352 proteins) than in PUJ256 (257 proteins) (**Table 2**). The complete global proteomic datasets, including volcano plots for all pairwise comparisons, are presented in **Supplementary tables 1-4** and **Supplementary figures 1-4.**

### Comparative functional enrichment analysis of the PUJ256 and SC5314 proteomes

When comparing the proteomes of both strains in the absence of fluconazole, the first notable observation is the enrichment of proteins localized to the mitochondria in the resistant isolate (**Figure 9A**), including components of the electron transport chain (Cox1, Cox2, Qcr8) and the tricarboxylic acid cycle (Cit1, Mdh1, Mdh1-3) (**Table 3**) showing an enrichment in proteins related with mitochondrion organization process and in mitochondrion cellular compartment (**Figure 9A**). Proteins related to vesicular trafficking and secretion, such as Arf3, Vps28, Ist1, Arc19, Sft1 and Ypt72 were also more abundant (**Table 3**) in PUJ256 isolate exhibiting an enrichment in proteins related with intracellular protein transport. The resistant isolate also displayed higher abundance of proteins involved in ergosterol biosynthesis (Erg11, Erg2, Erg3, Erg6) and sterol regulation (Ebp7) (**Table 3**). Regarding the comparison of the proteome from both strains under fluconazole exposure, a more pronounced proteomic shift was observed in SC5314 (**Figure 9B**). Among the 261 proteins significantly more abundant in this strain, the enriched biological processes comprised peroxisome organization, apoptosis, transmembrane transport, and cell wall and membrane organization, being these proteins mainly localized to the plasma membrane and extracellular regions.

**Figure 9.**
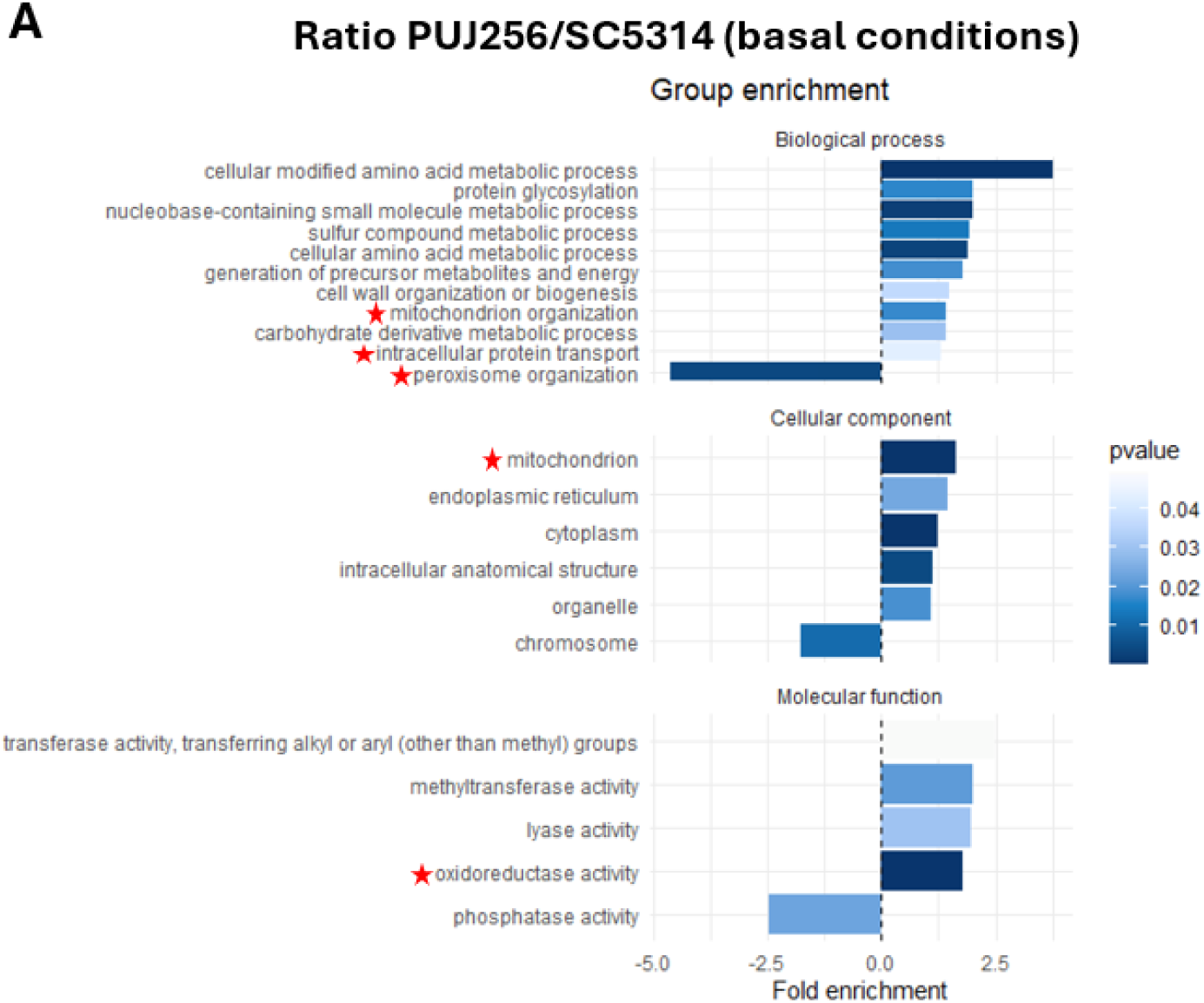
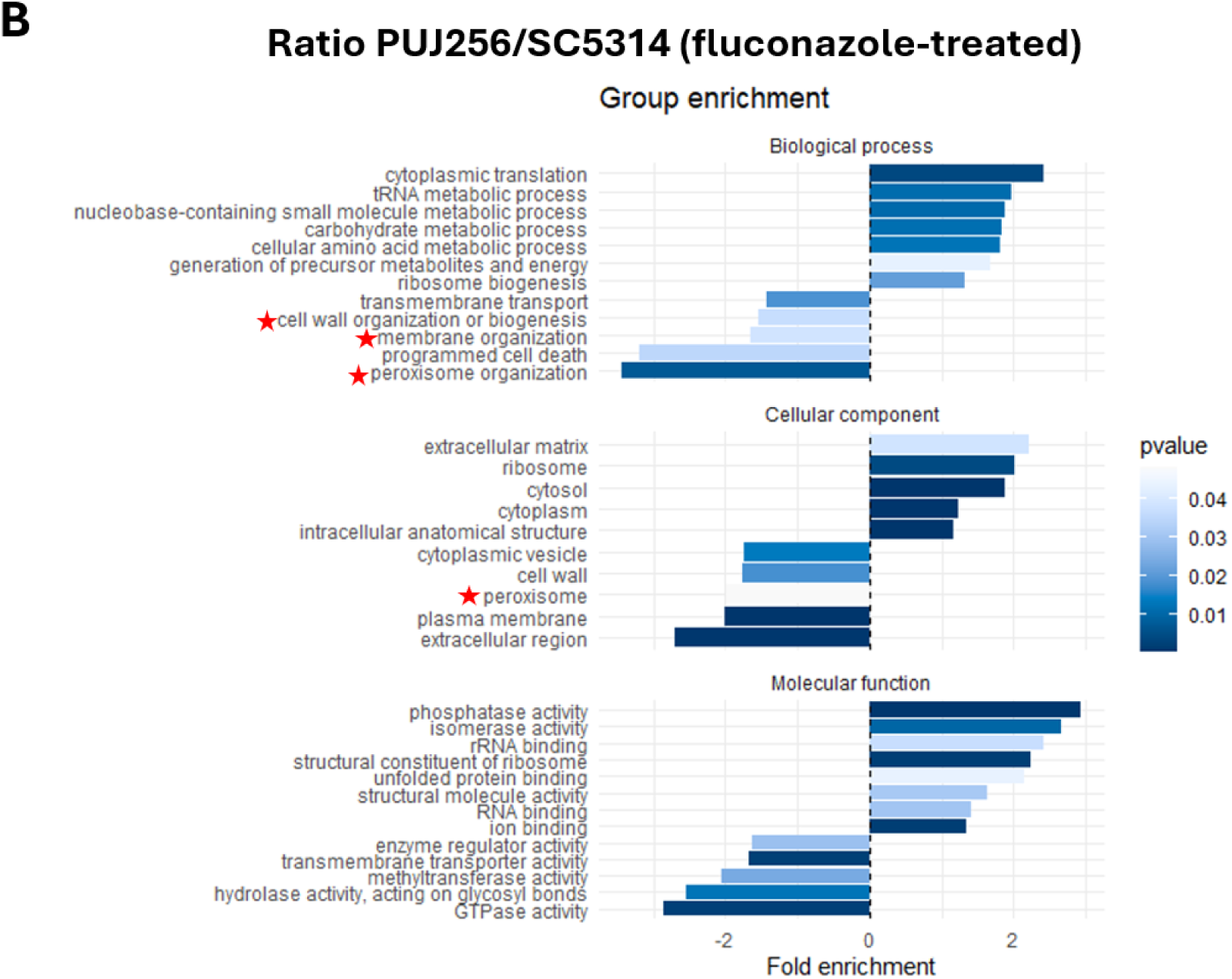
Gene Ontology (GO) functional enrichment analysis of differentially abundant proteins identified between PUJ256 and SC5314 under basal **(A)** and fluconazole-treated (1.1 μg/mL) **(B)** conditions. Biological process, cellular component and molecular function categories are shown. Positive fold enrichment values indicate GO terms enriched among proteins more abundant in PUJ256, whereas negative values indicate GO terms enriched among proteins more abundant in SC5314. The color intensity of the terms indicates the significance of the enrichment, representing *p*-values.

**Table 3.**
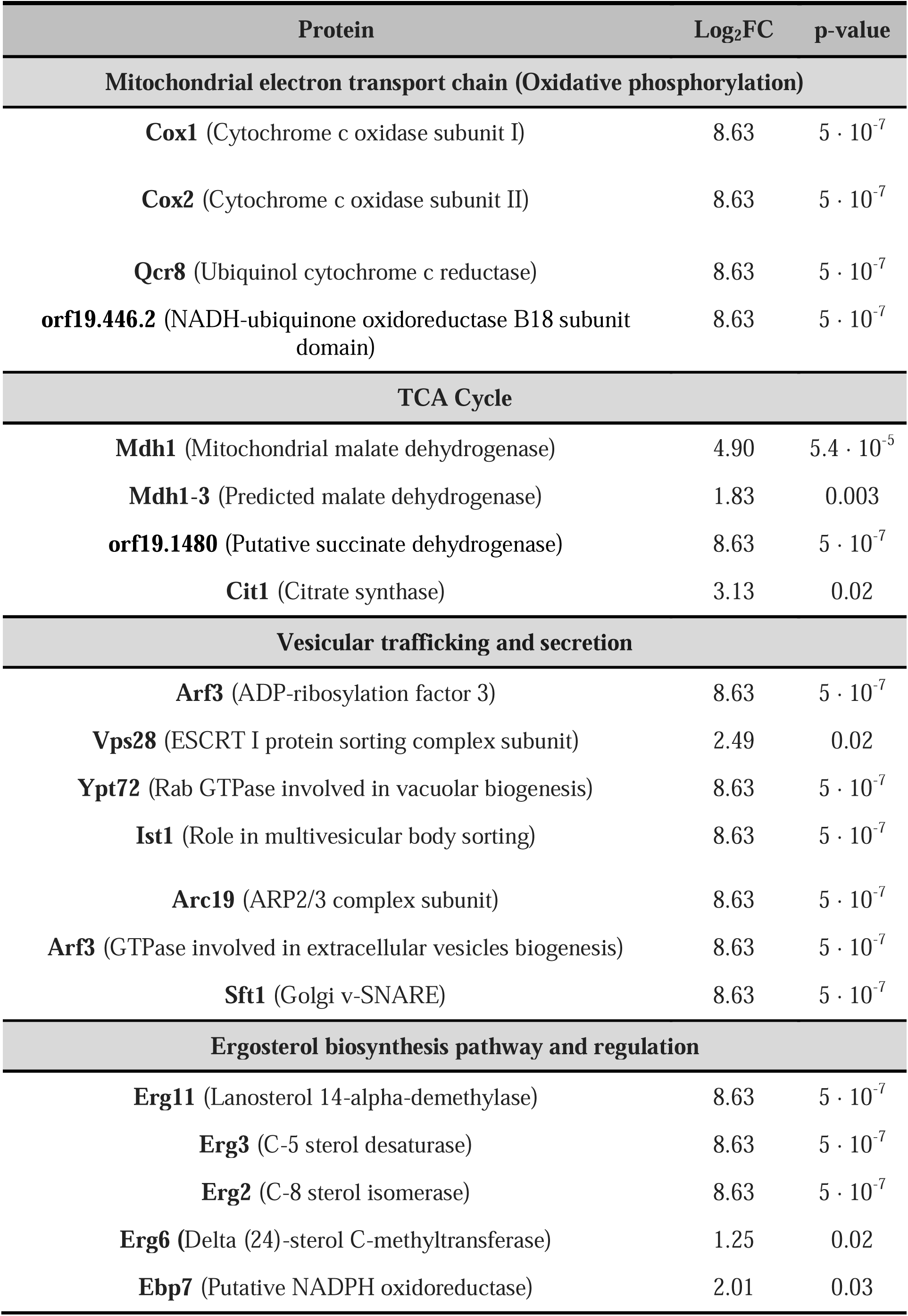
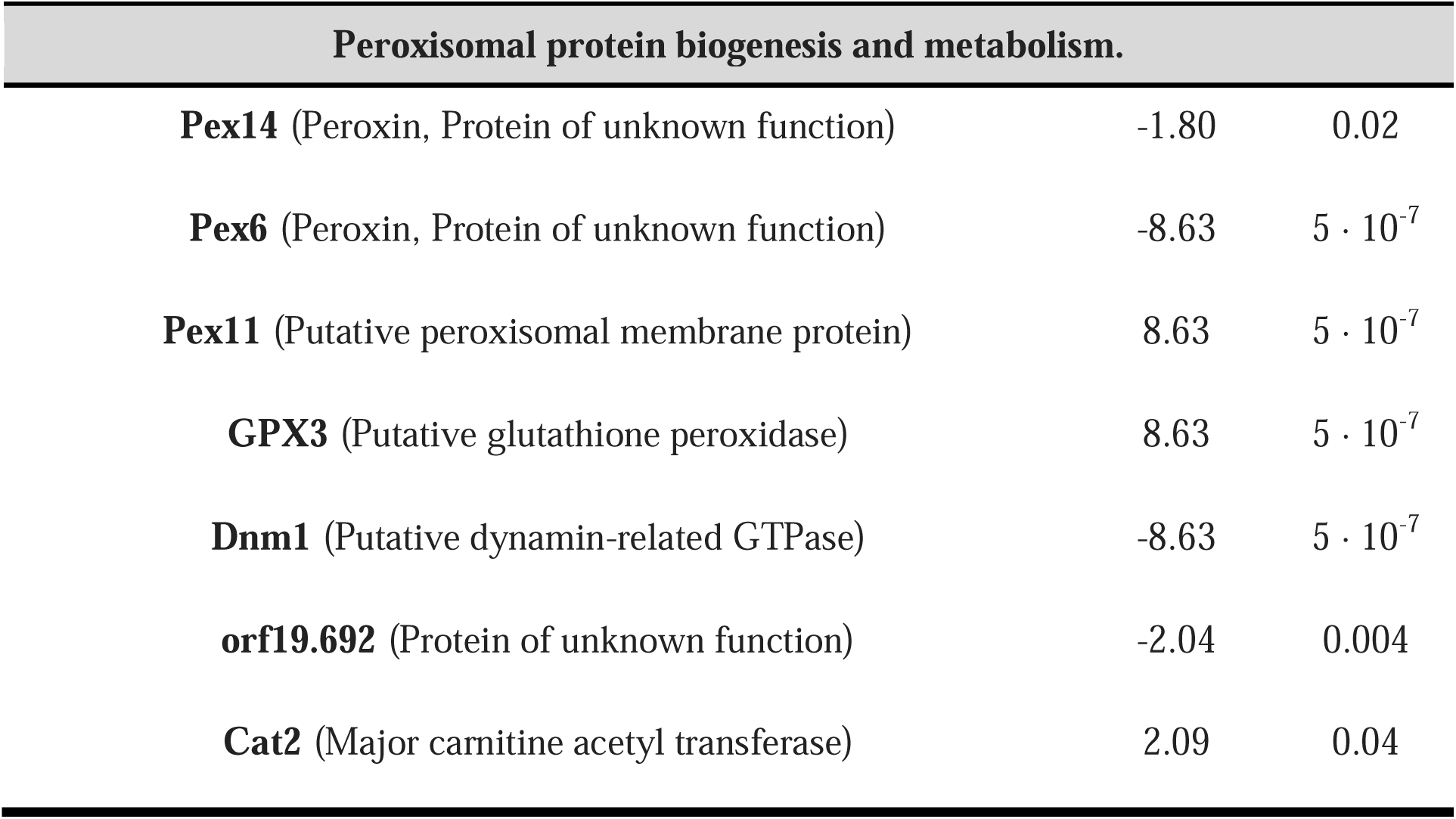
Log2 fold change (PUJ256/SC5314) of selected proteins associated with the biological processes and cellular compartments enriched in the GO analysis shown in Figure 9 under basal conditions. Positive values indicate higher abundance in PUJ256, whereas negative values indicate higher abundance in SC5314. A log_2_FC (PUJ256/SC5314) of 8.63 indicates that this protein was detected exclusively in PUJ256, and a log_2_FC (PUJ256/SC5314) of-8.63 indicates that this protein was detected exclusively in SC5314.

| Protein | Log<sub>2</sub>FC | p-value |
| --- | --- | --- |
| **Mitochondrial electron transport chain (Oxidative phosphorylation)** |  |  |
| **Cox1** (Cytochrome c oxidase subunit I) | 8.63 | $5 \cdot 10^{-7}$ |
| **Cox2** (Cytochrome c oxidase subunit II) | 8.63 | $5 \cdot 10^{-7}$ |
| **Qcr8** (Ubiquinol cytochrome c reductase) | 8.63 | $5 \cdot 10^{-7}$ |
| **orf19.446.2** (NADH-ubiquinone oxidoreductase B18 subunit domain) | 8.63 | $5 \cdot 10^{-7}$ |
| **TCA Cycle** |  |  |
| **Mdh1** (Mitochondrial malate dehydrogenase) | 4.90 | $5.4 \cdot 10^{-5}$ |
| **Mdh1-3** (Predicted malate dehydrogenase) | 1.83 | 0.003 |
| **orf19.1480** (Putative succinate dehydrogenase) | 8.63 | $5 \cdot 10^{-7}$ |
| **Cit1** (Citrate synthase) | 3.13 | 0.02 |
| **Vesicular trafficking and secretion** |  |  |
| **Arf3** (ADP-ribosylation factor 3) | 8.63 | $5 \cdot 10^{-7}$ |
| **Vps28** (ESCRT I protein sorting complex subunit) | 2.49 | 0.02 |
| **Ypt72** (Rab GTPase involved in vacuolar biogenesis) | 8.63 | $5 \cdot 10^{-7}$ |
| **Ist1** (Role in multivesicular body sorting) | 8.63 | $5 \cdot 10^{-7}$ |
| **Arc19** (ARP2/3 complex subunit) | 8.63 | $5 \cdot 10^{-7}$ |
| **Arf3** (GTPase involved in extracellular vesicles biogenesis) | 8.63 | $5 \cdot 10^{-7}$ |
| **Sft1** (Golgi v-SNARE) | 8.63 | $5 \cdot 10^{-7}$ |
| **Ergosterol biosynthesis pathway and regulation** |  |  |
| **Erg11** (Lanosterol 14-α-demethylase) | 8.63 | $5 \cdot 10^{-7}$ |
| **Erg3** (C-5 sterol desaturase) | 8.63 | $5 \cdot 10^{-7}$ |
| **Erg2** (C-8 sterol isomerase) | 8.63 | $5 \cdot 10^{-7}$ |
| **Erg6** (Delta (24)-sterol C-methyltransferase) | 1.25 | 0.02 |
| **Ebp7** (Putative NADPH oxidoreductase) | 2.01 | 0.03 |
| Peroxisomal protein biogenesis and metabolism. |  |  |
| **Pex14** (Peroxin, Protein of unknown function) | -1.80 | 0.02 |
| **Pex6** (Peroxin, Protein of unknown function) | -8.63 | $5 \cdot 10^{-7}$ |
| **Pex11** (Putative peroxisomal membrane protein) | 8.63 | $5 \cdot 10^{-7}$ |
| **GPX3** (Putative glutathione peroxidase) | 8.63 | $5 \cdot 10^{-7}$ |
| **Dnm1** (Putative dynamin-related GTPase) | -8.63 | $5 \cdot 10^{-7}$ |
| **orf19.692** (Protein of unknown function) | -2.04 | 0.004 |
| **Cat2** (Major carnitine acetyl transferase) | 2.09 | 0.04 |

When assessing the effect of fluconazole exposure within each strain (**Figure 10**), distinct proteomic changes were observed. In SC5314, proteins that increased in abundance following fluconazole treatment were significantly enriched in biological processes related to cell wall biogenesis and organization, as well as other extracellular and cell wall-associated functions (**Table 4**). Consistent with these enrichments, the predominant cellular localizations of these proteins were the extracellular region, plasma membrane, and endoplasmic reticulum (Figure 10a). Fluconazole treatment also increased the abundance of proteins involved in ergosterol biosynthesis and membrane homeostasis (**Table 4**), together with proteins associated with vesicular trafficking and secretion, and several stress response and signalling proteins. Conversely, proteins that decreased in abundance following fluconazole treatment in SC5314 were predominantly involved in translation, ribosome biogenesis, cell cycle progression, and DNA replication.

**Figure 10.**
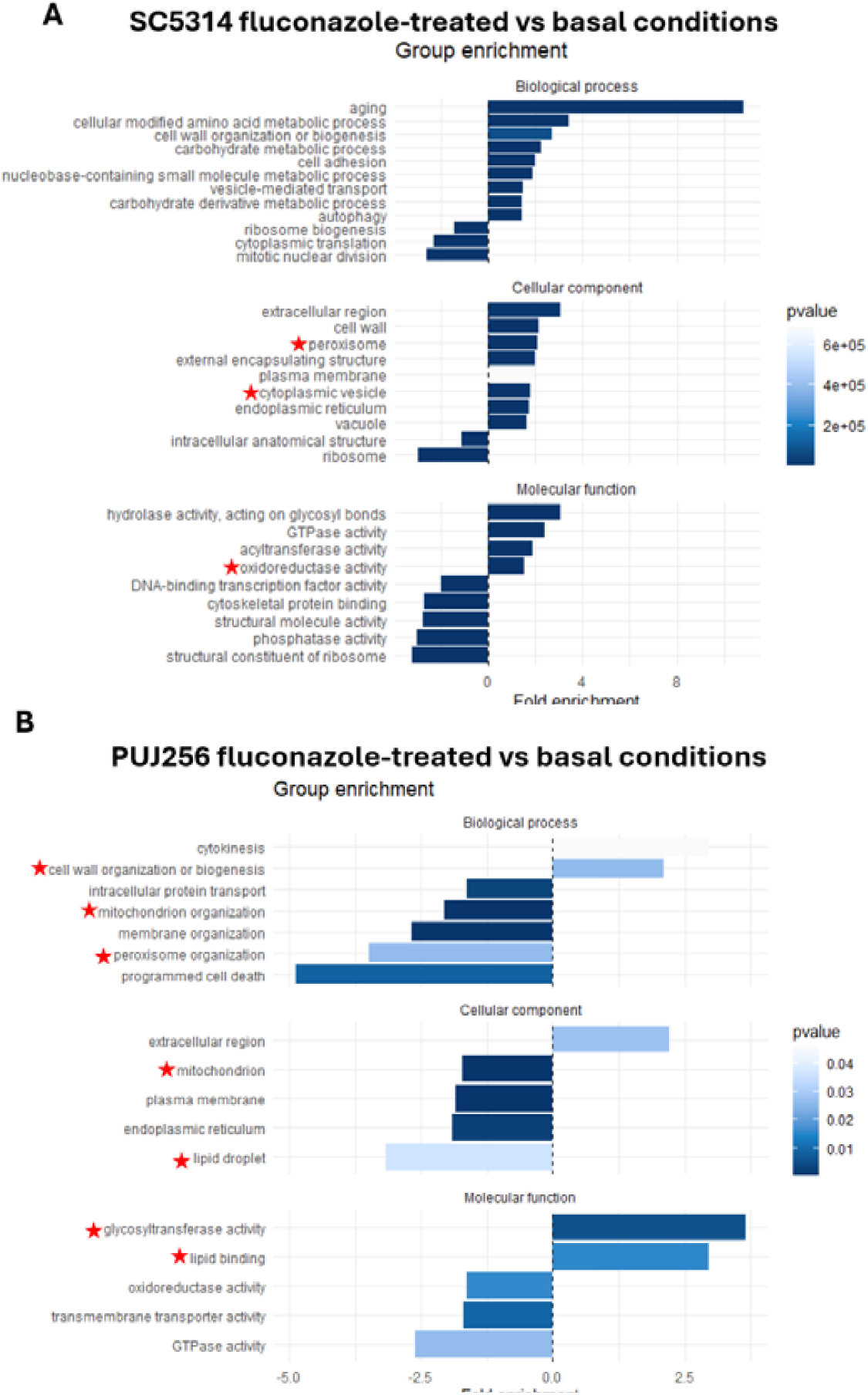
GO functional enrichment analysis of differentially abundant proteins in SC5314 **(A)** and PUJ256 **(B)** following fluconazole treatment (1.1 μg/mL) compared with basal conditions. Biological process, cellular component and molecular function categories are shown. Positive fold enrichment values indicate GO terms enriched among proteins with increased abundance following fluconazole treatment, whereas negative values indicate GO terms enriched among proteins more abundant under basal conditions. The colour intensity of the terms indicates the significance of the enrichment, representing *p*-values.

**Table 4.** Table 4. Log2 fold change (SC5314_FLU/SC5314) of selected proteins associated with the biological processes and cellular compartments enriched in the GO analysis shown in Figure 10. Positive values indicate higher abundance following fluconazole treatment (1.1 _μ_g/mL). A log_2_FC (SC5314_FLU/SC5314) of 8.63 indicates that this protein was detected exclusively in SC5314_FLU.

| Protein | Log<sub>2</sub>FC | p-value |
| --- | --- | --- |
| **Cell wall remodelling** |  |  |
| **Mnn9** (Protein of N-linked outer-chain mannan biosynthesis) | 8.63 | $5 \cdot 10^{-7}$ |
| **Mnt2** (Alpha-1,2-mannosyl transferase) | 8.63 | $5 \cdot 10^{-7}$ |
| **Kre6** (Essential beta-1,6-glucan synthase subunit) | 3.17 | 0.0009 |
| **Crh11** (GPI-anchored cell wall transglycosylase) | 2.00 | 0.003 |
| **Bgl2** (Cell wall 1,3-beta-glucosyltransferase) | 1.33 | 0.01 |
| **Phr2** (Glycosidase) | 2.65 | 0.0002 |
| **Xog1** (Exo-1,3-beta-glucanase) | 5.42 | $1.7 \cdot 10^{-5}$ |
| **Ergosterol biosynthesis pathway and membrane remodulation** |  |  |
| **Erg11** (Lanosterol 14-alpha-demethylase) | 8.63 | $5 \cdot 10^{-7}$ |
| **Erg3** (C-5 sterol desaturase) | 8.63 | $5 \cdot 10^{-7}$ |
| **Erg6** (Delta (24)-sterol C-methyltransferase) | 2.50 | $9 \cdot 10^{-4}$ |
| **Sur2** (Putative ceramide hydroxylase) | 8.63 | $5 \cdot 10^{-7}$ |
| **Are2** (Acyl CoA:sterol acyltransferase) | 1.51 | 0.02 |
| **Vesicular trafficking and secretion** |  |  |
| **Exo84** (Predicted subunit of the exocyst complex) | 8.63 | $5 \cdot 10^{-7}$ |
| **Sec3** (Predicted subunit of the exocyst complex) | 1.62 | 0.04 |
| **Vps21** (Late endosomal Rab small monomeric GTPase) | 2.19 | 0.01 |
| **Trs33** (Putative TRAPP complex subunit) | 8.63 | $5 \cdot 10^{-7}$ |
| **Transcriptional and stress regulation** |  |  |
| **Ddr48** (Immunogenic stress-associated protein) | 3.80 | 0.003 |
| **Gtt12** (Protein of unknown function) | 8.63 | $5 \cdot 10^{-7}$ |
| **Snf1** (Essential protein) | 8.63 | $5 \cdot 10^{-7}$ |
| **Gcn5** (Putative histone acetyltransferase) | 8.63 | $5 \cdot 10^{-7}$ |
| **Spt7** (Putative SAGA transcriptional regulatory complex subunit) | 8.63 | $5 \cdot 10^{-7}$ |
| **Ccr4** (Component of the Ccr4-Pop2 mRNA deadenylase) | 8.63 | $5 \cdot 10^{-7}$ |
| **Bur2** (Cyclin for the Sgv1p (Bur1p) protein kinase) | 8.63 | $5 \cdot 10^{-7}$ |

A similar analysis was performed for the clinical isolate PUJ256. Among the 86 proteins that increased significantly in abundance following fluconazole exposure, the most enriched molecular functions were glycosyltransferase activity and lipid assembly, particularly sterol biosynthesis (**Figure 10B**). These proteins were mainly localized to the extracellular region and the phagophore assembly site involved in autophagosome formation (**Figure 10B**; **Table 5**). Conversely, the 171 proteins that decreased in abundance following fluconazole exposure were enriched in biological processes related to programmed cell death, membrane organization, and mitochondrial organization, and were primarily localized to the plasma membrane, mitochondria, and endoplasmic reticulum (**Figure 10B**; **Table 5**).

**Table 5.** Table 4. Log2 fold change (PUJ256_FLU/PUJ256) of selected proteins associated with the biological processes and cellular compartments enriched in the GO analysis shown in Figure 10. Positive values indicate higher abundance following fluconazole treatment (1.1 _μ_g/mL), whereas negative values indicate higher abundance under basal conditions. A log_2_FC (PUJ256_FLU/PUJ256) of 8.63 indicates that this protein was detected exclusively in PUJ256_FLU, and a log_2_FC (PUJ256_FLU/PUJ256) of-8.63 indicates that this protein was detected exclusively in PUJ256.

| Protein | Log<sub>2</sub>FC | p-value |
| --- | --- | --- |
| **Cell wall remodelling** |  |  |
| **Chs2** (Chitin synthase) | 8.63 | $5 \cdot 10^{-7}$ |
| **Chs8** (Chitin synthase) | 8.63 | $5 \cdot 10^{-7}$ |
| **Scw11** (Cell wall protein) | 1.39 | 0.03 |
| **Wsc4** (Putative cell wall integrity and stress response subunit 4 precursor) | 8.63 | $5 \cdot 10^{-7}$ |
| **Xog1** (Exo-1,3-beta-glucanase) | 1.95 | 0.004 |
| **Sphingolipid biosynthesis** |  |  |
| **MIT1** (Mannosylinositol phosphorylceramide synthase) | 8.63 | $5 \cdot 10^{-7}$ |
| **Mitochondrial protein import and respiratory function** |  |  |
| **Cox2** (Subunit II of cytochrome c oxidase) | -8.63 | $5 \cdot 10^{-7}$ |
| **Qcr8** (Putative ubiquinol cytochrome c reductase) | -8.63 | $5 \cdot 10^{-7}$ |
| **Sdh1** (Putative mitochondrial succinate dehydrogenase) | -8.63 | $5 \cdot 10^{-7}$ |
| **Sco1** (Putative copper transporter) | -8.63 | $5 \cdot 10^{-7}$ |
| **Tom40** (Protein involved in mitochondrial protein import) | -8.63 | $5 \cdot 10^{-7}$ |
| **Tom7** (Predicted component of the TOM (translocase of outer membrane) complex) | -8.63 | $5 \cdot 10^{-7}$ |
| **Pam18** (Predicted component of the presequence translocase-associated import motor (PAM complex)) | -8.63 | $5 \cdot 10^{-7}$ |

### Mitochondrial function

Given that the proteomic analysis revealed an upregulation of proteins involved in cellular respiration in PUJ256 (**Figure 11B**), we investigated whether mitochondrial homeostasis contributes to the resistant phenotype by assessing mitochondrial membrane potential using the JC-1fluorogenic probe, where the red/green fluorescence ratio is used as a measure of mitochondrial polarization (**Figure 11A**). Under basal conditions, the susceptible reference strain SC5314 exhibited significantly higher mitochondrial polarization compared to the clinical isolate PUJ256. However, fluconazole exposure (1.1 μg/mL) significantly decrease mitochondrial polarization in SC5314, evidenced by a decline in the red/green ratio **(Figure 11A**). In contrast, under basal conditions PUJ256 displayed a significantly lower mitochondrial membrane potential than SC5314. Notably, this difference was not maintained under fluconazole exposure; rather, PUJ256 preserved mitochondrial polarization and even showed a trend towards increased membrane potential, in contrast to the marked depolarization observed in SC5314.

**Figure 11.**
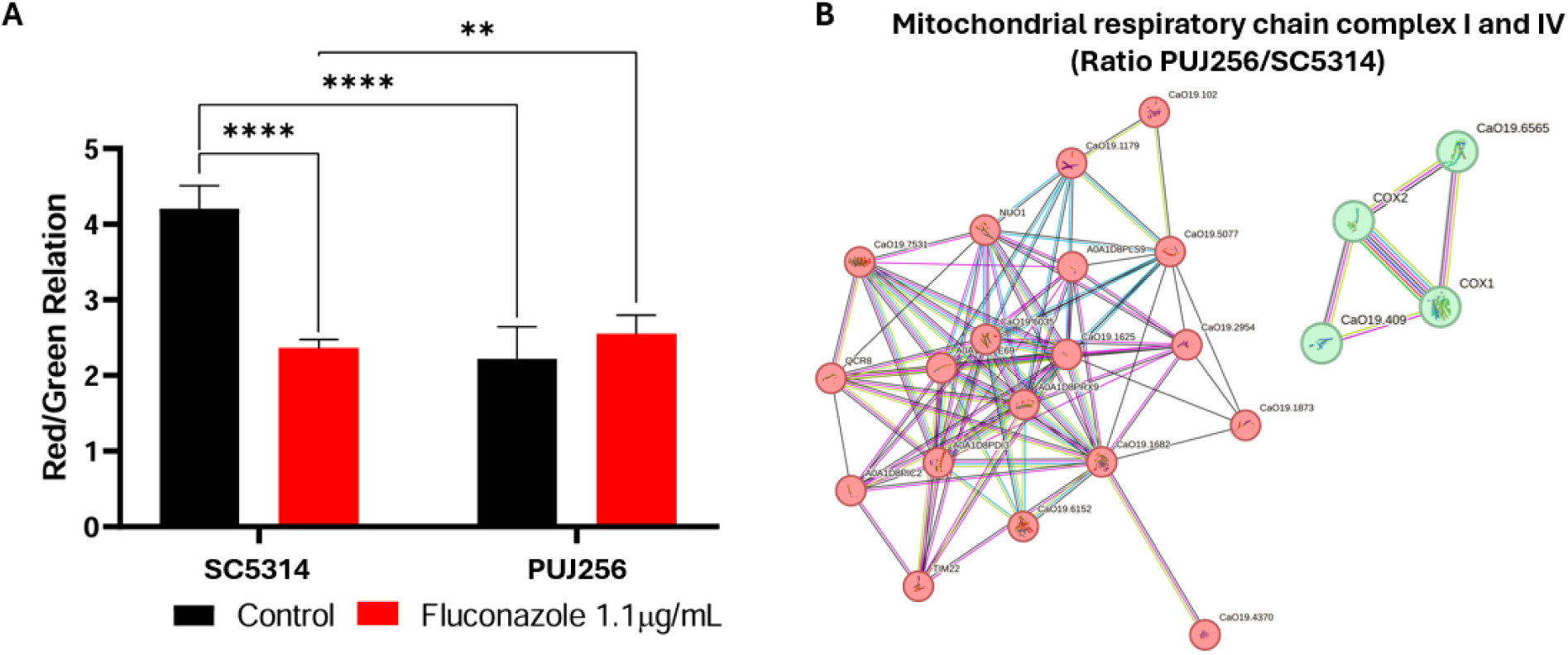
(A) Assessment of mitochondrial membrane potential after 4 hours of fluconazole exposure using JC-1 staining. Quantification of the red/green fluorescence ratio in SC5314 and PUJ256 grown in the absence (control) or presence of fluconazole (1.1 µg/mL). Data represent mean ± SD (n = 3); statistical significance was determined by ANOVA. **(B)** Protein-protein interaction network of proteins related to mitochondrial respiratory chain complex I and IV using STRING software.

### MAPK pathways detection

Given the extensive remodelling of cell wall, membrane and redox-associated processes revealed by the proteomic analysis, we next examined whether these adaptive responses were associated with differential activation of stress-responsive MAPK pathways by assessing the phosphorylation of Mkc1, Hog1 and Cek1 in SC5314 and PUJ256, the terminal MAPKs in the cell wall integrity, filamentation and high osmolarity (HOG) pathways following fluconazole exposure. Both strains exhibited increased phosphorylation of Mkc1 and Cek1 after 30 minutes of incubation with the drug, consistent with activation of the CWI and filamentation pathways. In contrast, Hog1 phosphorylation was detected earlier, after 10 minutes of exposure, indicating a rapid stress response. Higher levels of phosphorylated Mkc1 and Cek1 were observed in the fluconazole-susceptible strain compared to the azole-resistant isolate, whereas Hog1 activation followed a distinct pattern, in agreement with the proteomic data (**Figure 12**).

**Figure 12.**
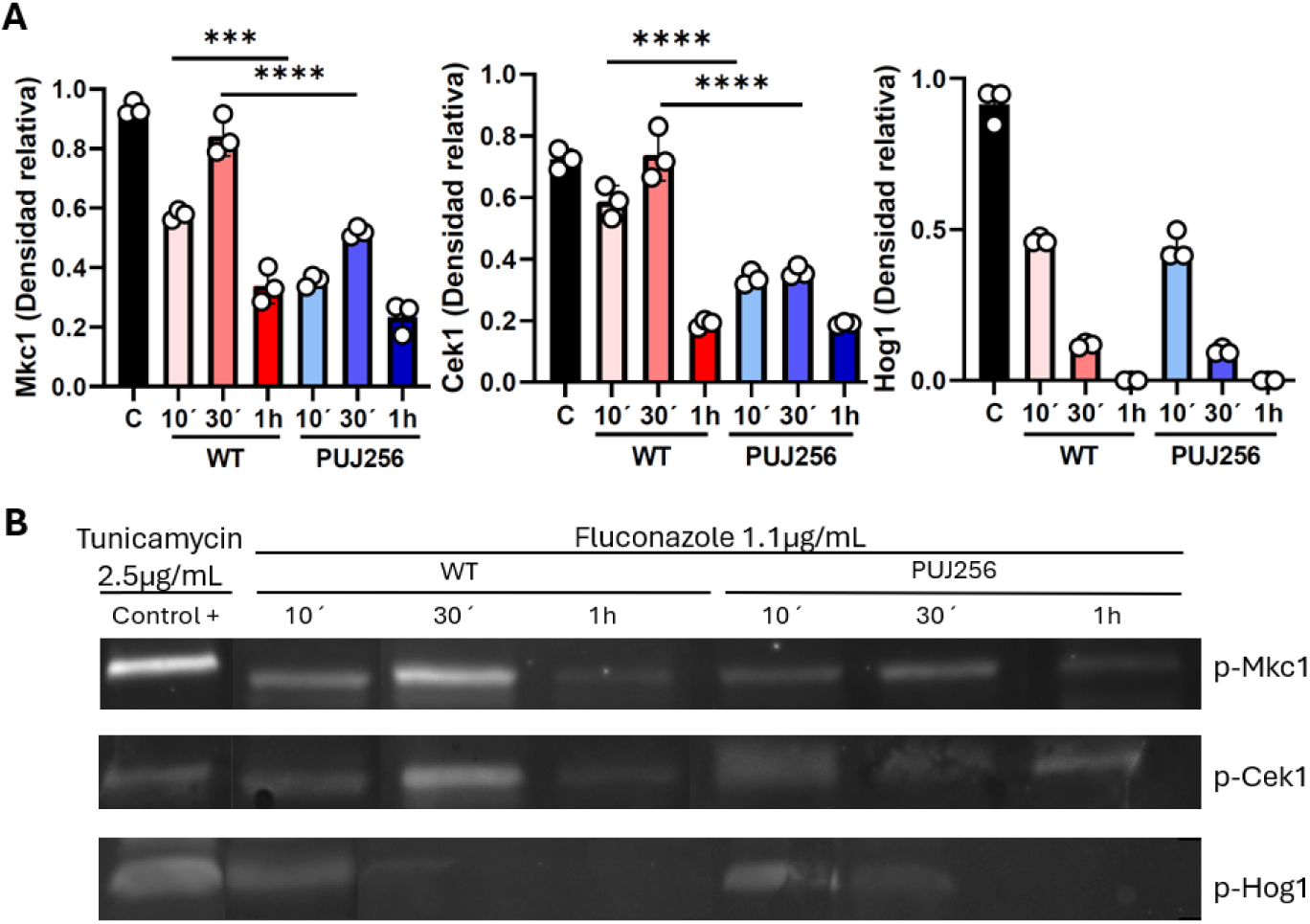
Activation of MAPK signalling pathways in response to fluconazole exposure. **(A)** Quantification of phosphorylated Mkc1, Cek1, and Hog1 in SC5314 (WT) and PUJ256 following treatment with fluconazole (1.1 µg/mL) for the indicated times. Data represent mean ± SD (n = 3); statistical significance was determined by ANOVA. **(B)** Representative Western blot images showing p-Mkc1, p-Cek1, and p-Hog1 under the indicated conditions. Tunicamycin (2.5 µg/mL) was used as a positive control for pathway activation. Total protein loading was assessed by Ponceau Red staining.

**Figure 13.**
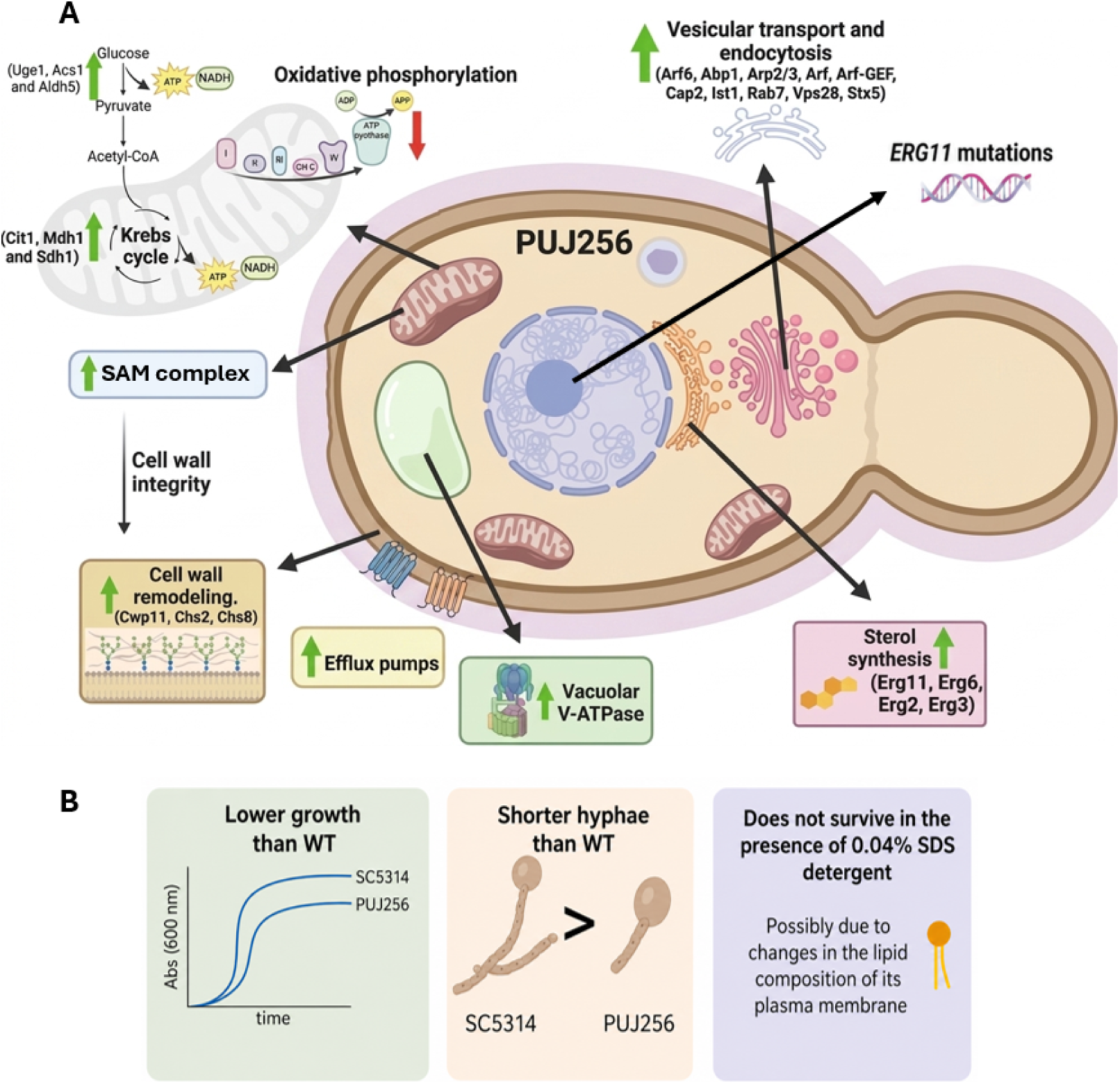
Schematic representation of the adaptive cellular and metabolic responses in PUJ256 compared to the reference strain SC5314 under fluconazole treatment.

## Discussion

*Candida albicans* is a highly adaptable opportunistic pathogen whose prolonged exposure to azole antifungals can drive the emergence of resistance to high drug concentrations (Singh et al., 2012). In this study, we analysed a clinical isolate (PUJ256) recovered after more than 30 years of continuous fluconazole therapy, providing a unique model to investigate the physiological basis and potential clinical consequences of long-term azole resistance (Cárdenas et al., 2020; Ceballos-Garzón et al., 2020).

Mutations in *ERG11* constitute one of the best-characterized mechanisms of azole resistance in *C. albicans* (Flowers et al., 2014; Healey et al., 2018). Sequence analysis of PUJ256 identified five amino acid substitutions located within or adjacent to previously described functional hotspot regions of Erg11 (**Figure 1**, **Table 1**) (Xiang et al., 2013). However, among these substitutions, only S405F has been experimentally demonstrated to directly reduce azole susceptibility, whereas E266D and V488I are frequently reported as naturally occurring polymorphisms, and the contribution of S263L and V402F remains unclear (Flowers et al., 2015). PUJ256 also displayed enhanced fluconazole-induced efflux activity, indicating that active drug extrusion constitutes an additional contributor to its resistant phenotype (Sanglard et al., 1995; Cowen et al., 2014). Nevertheless, these classical mechanisms alone are unlikely to account for the extensive phenotypic and proteomic remodeling observed in this isolate. Instead, our findings support the view that long-term azole resistance results from the integration of canonical resistance mechanisms with broader physiological adaptations acquired during prolonged antifungal exposure (Healey et al., 2018).

Our competition assays revealed a clear environment-dependent fitness trade-off, in which the azole-sensitive strain outcompeted the resistant isolate in the absence of fluconazole, whereas the opposite trend was observed under drug exposure (**Figure 3C**). This pattern is consistent with previous experimental evolution and in vitro studies in *C. albicans* (Popp et al., 2017), where sensitive strains exhibited higher basal fitness, and by Cowen et al., 2001, who demonstrated that resistance confers a crucial selective advantage under antifungal pressure despite associated biological costs. Notably, the presence of a detectable fitness cost in our resistant isolate suggests that compensatory adaptations, commonly reported to mitigate these costs over time, may be incomplete or absent in this strain (Cowen et al., 2001).

Fluconazole has been reported to inhibit filamentation in *C. albicans* (Kontoyiannis et al., 2001), and our quantitative analysis confirms that hyphal elongation is markedly impaired in the susceptible strain under antifungal pressure (**Figure 4A**). Interestingly, the resistant isolate did not exhibit a marked reduction in hyphal length under fluconazole pressure while the antifungal caused a nearly fivefold reduction in the SC5314 (**Figure 4B**). This indicates that, in the resistant isolate, morphogenetic responses become less sensitive to antifungal stress, suggesting a partial uncoupling between filamentation and drug susceptibility during long-term adaptation. In line with this, SC5314 displayed activation of MAPK signalling pathways (**Figure 12**) and a reduction in phosphatase-associated functions upon fluconazole exposure (**Figure 10A**), consistent with a dynamic stress response. In contrast, PUJ256 showed lower MAPK activation and no significant changes in phosphatase-related functions (**Figure 10B**), supporting a model in which long-term adaptation leads to a more stable, pre-adapted physiological state that reduces reliance on inducible signalling responses. This phenotype has been previously associated with intrinsic alterations in lipid composition and mitochondrial function in resistant strains, which may affect morphogenetic signalling pathways (Singh et al., 2012). Under basal conditions, PUJ256 also exhibited lower mitochondrial membrane polarization compared with SC5314, as indicated by JC-1 staining (**Figure 11A**), suggesting altered basal bioenergetic status in the resistant isolate. Consistent with this phenotype, comparative proteomic analysis revealed significant remodeling of organelles involved in energy metabolism. In addition to the enrichment of mitochondrial-associated proteins (**Figure 9A**), proteins involved in peroxisome organization were less abundant in PUJ256 than in SC5314 (**Table 3**). Since peroxisomes and mitochondria cooperate in fatty acid metabolism, redox homeostasis and acetyl-CoA trafficking, these changes suggest a coordinated reorganization of energy metabolism rather than an isolated alteration of mitochondrial function. However, upon fluconazole exposure, the mitochondrial membrane potential of SC5314 was significantly impaired, whereas it was maintained, and even increased, in PUJ256 (**Figure 11A**). Interestingly, GO enrichment analysis also revealed a reduction in proteins associated with peroxisome organization following fluconazole exposure in PUJ256 (**Figure 10B**), suggesting that long-term azole adaptation is accompanied by sustained remodeling of the peroxisomal compartment. Together with the preservation of mitochondrial membrane potential, these observations support the existence of a constitutively reprogrammed metabolic state that minimizes the need for extensive stress-induced organelle remodeling upon antifungal challenge. In fact, when comparing the proteomes from SC5314 and PUJ256 strains, an enrichment in the proteins associated with mitochondria was also observed in the resistant isolate (**Figure 9A**). At the metabolic level, PUJ256 displayed increased abundance of proteins related to glycolysis and the tricarboxylic acid cycle (**Table 3**), possibly promoting NADH generation and supporting regeneration of NAD through complex I activity, consistent with the increased abundance of proteins from this complex (**Figure 11B**). Although proteins belonging to respiratory complexes II and III were also more abundant in the resistant isolate (**Supplementary Tables 1 and 4**), this did not correlate with higher mitochondrial membrane polarization under basal conditions (**Figure 11A**), suggesting that mitochondrial membrane potential is determined not only by respiratory protein abundance but also by electron transport efficiency and coupling. However, it is remarkable that upon fluconazole exposure, the mitochondrial membrane potential of the SC5314 was significantly impaired whereas it was preserved, and even enhanced, in the resistant isolate under antifungal stress (**Figure 11A**). Furthermore, the increase abundance of Sam51 (**Supplementary table 1**), a core component of the SAM complex, further supports the involvement of mitochondrial remodeling in the resistant phenotype. The SAM complex mediates phospholipid trafficking between the endoplasmic reticulum and mitochondria and is essential for mitochondrial membrane organization and cell wall integrity (Shingu-Vazquez and Traven, 2011).

Interestingly, although hyphal length in the resistant isolate did not change upon fluconazole exposure (**Figure 4B**), we observed a significant increase in its ability to form biofilm compared with basal conditions (**Figure 8**) while no significant change was observed in the SC5314. This phenotype can be partially explained by our proteomic data (**Figure 10B**), which revealed a stronger enrichment of glycosyltransferase activity and lipid binding functions in the resistant isolate upon fluconazole exposure compared with SC5314. Such enrichment may facilitate enhanced biofilm formation in the resistant isolate.

Membrane-associated trade-offs were evident in the resistant isolate, which exhibited increased sensitivity to SDS compared to the wild-type strain (**Figure 6C**). This phenotype is consistent with altered plasma membrane lipid composition. Reduced phosphatidylglycerol levels have been described to increase susceptibility to membrane-disrupting agents (Singh et al., 2012). Proteomic analyses revealed that fluconazole resistance in PUJ256 is strongly associated with remodelling of ergosterol biosynthesis and central metabolism. Multiple enzymes of the ergosterol and sphingolipids pathway (Erg2, Erg3, Erg6, Erg11 and Ebp7), under the control of the transcription factor Upc2, were more abundant in the resistant isolate than in the wild-type strain (**Table 3**), consistent with their established role in azole resistance (de Backer et al., 2001; Flowers et al., 2012). In particular, the increased abundance of Erg6 suggests compensatory production of C-24 alkylated sterols that may preserve membrane integrity in the presence of fluconazole but could also be responsible for the increased sensitivity to SDS detergent. These findings support the notion that azole resistance entails compensatory physiological costs that reshape membrane properties while preserving viability under antifungal pressure.

Proteomic profiling also revealed a constitutively higher abundance of multiple V-ATPase subunits in the resistant isolate compared with SC5314 under basal conditions, a pattern that persisted following fluconazole exposure. Since fluconazole-induced ergosterol depletion destabilizes the lipid rafts required for V-ATPase assembly and coupling efficiency (Zhang et al., 2010), this increased abundance may represent a compensatory response to maintain vacuolar function under conditions of membrane perturbation. By expanding the pool of available subunits, the resistant strain could partially offset impaired proton pumping and maintain intracellular homeostasis under antifungal stress. Consequently, this protein shift could function as an adaptive buffer, maintaining the electrochemical gradients necessary to drive Ca^2+^/H^+^ exchange and prevent cytosolic acidification, thereby sustaining cellular viability under chronic antifungal pressure.

The increased survival of PUJ256 to macrophage interaction in the presence of fluconazole (**Figure 3B**) appears to be closely associated with its reduced phagocytosis under the same conditions (**Figure 7B**), suggesting that decreased uptake rather than enhanced resistance to intracellular killing underlies this phenotype. Notably, this effect is condition-dependent, as under basal conditions PUJ256 was more efficiently phagocytosed than SC5314. This increased uptake in the absence of fluconazole may be related, at least in part, to its altered morphology, as PUJ256 exhibits shorter and defective hyphae. Morphological transitions are known to influence macrophage interactions, and although the relationship is complex, differences in hyphal architecture can affect recognition and engulfment (Lewis et al., 2012). Previous studies showed that *Cek1* mutants with impaired filamentation are more readily phagocytosed than its wild type strain (Román et al., 2016).

Under fluconazole exposure, proteomic analysis revealed increased abundance of several proteins associated with cell wall organization and remodeling in PUJ256, including the chitin synthase Chs2 and Chs8, the cell wall integrity-related proteins Ecm7, Scw11 and Wsc4, as well as Xog1 and the phospholipase Plb1 (**Table 5, Supplementary table 3**). These proteins are involved in cell wall biogenesis, remodeling and secretion, processes that can influence the composition and architecture of the fungal surface. Although fluconazole induced cell wall remodeling in both SC5314 and PUJ256, proteins associated with these processes were consistently more abundant in the resistant isolate. Interestingly, this occurred despite a markedly weaker activation of MAPK signalling in PUJ256 than in SC5314 following fluconazole exposure. These observations suggest that the resistant isolate is constitutively primed for cell wall remodelling, thereby reducing its dependence on a robust inducible cell wall integrity signalling response. Antifungal-induced cell wall remodeling has been shown to alter the exposure of immunogenic components such as β-glucan, thereby modulating recognition by macrophage receptors such as Dectin-1 (Wheeler and Fink, 2006; Walker et al., 2008; Gow et al., 2012). In this context, the observed changes may contribute to reduced macrophage recognition and uptake of PUJ256 specifically under antifungal stress.

The integration of phenotypic, proteomic, and mitochondrial analyses indicates that long-term azole resistance in *C. albicans* PUJ256 is supported by metabolic reprogramming and mitochondrial resilience that preserve cellular homeostasis under antifungal pressure. Rather than relying on exaggerated stress activation, the resistant isolate adopts a physiologically flexible state that promotes fitness and persistence during chronic fluconazole exposure. These insights highlight mitochondrial function and metabolic remodeling as central components of azole resistance and potential targets for therapeutic intervention.

