## Supplementary figures and images for "Thirty years of fluconazole therapy selects an azole-resistant *Candida albicans* isolate with a pre-adapted physiological, metabolic and structural state"

### Supplementary figure 1

a)

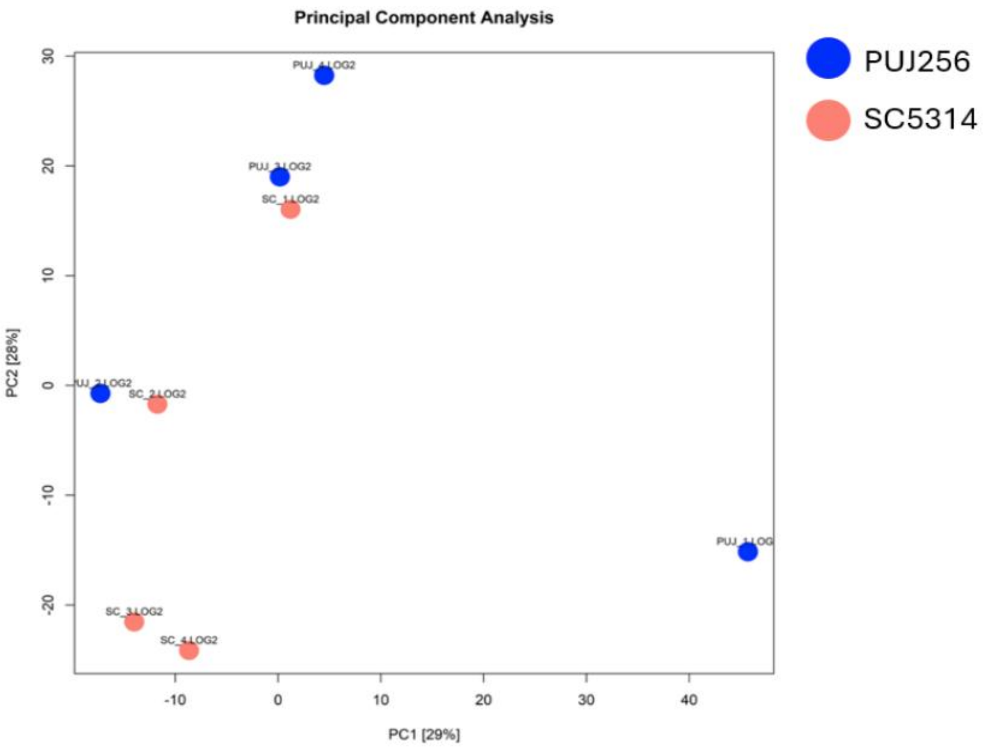


b)

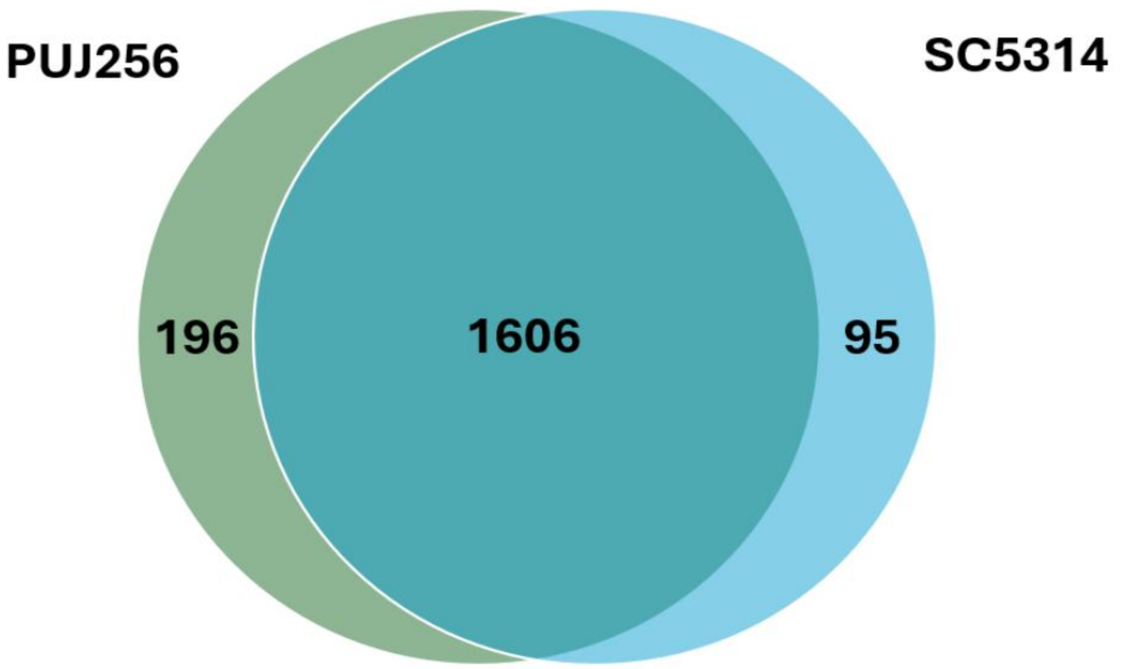


c)

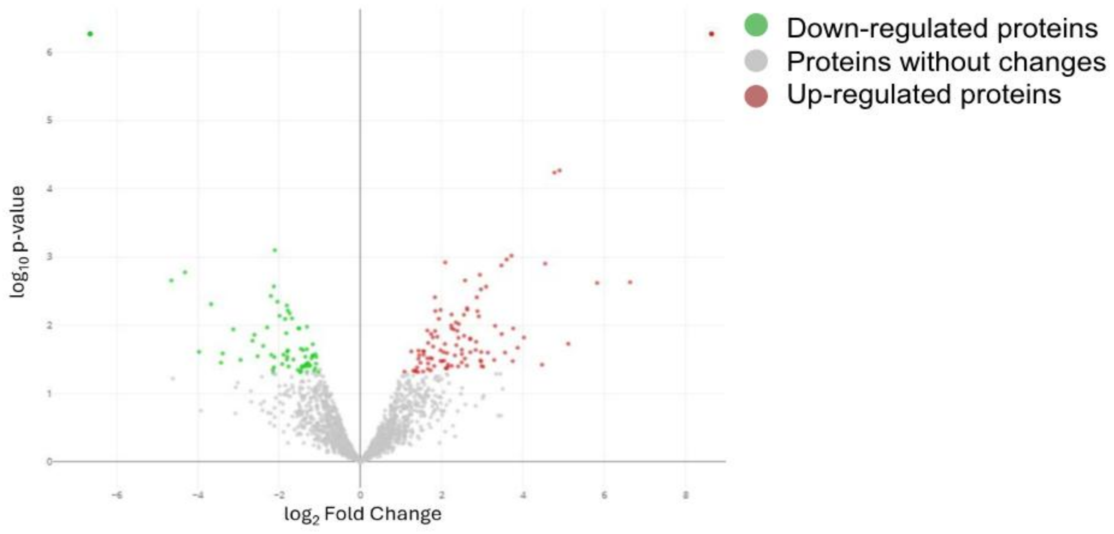


d)

| Condition | Quantified | Increased | Decreased |
|---|---|---|---|
| PUJ256 / SC5314 | 1896 | 298 | 163 |

e)

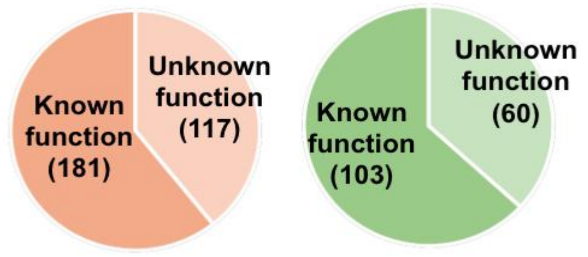

### Supplementary figure 2

a)

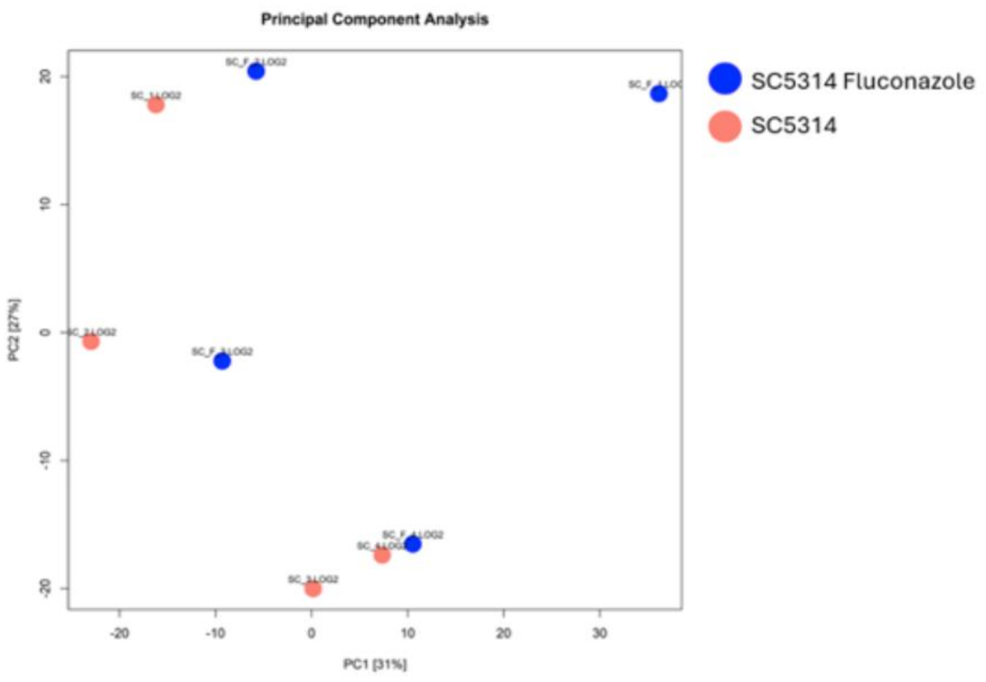


b)

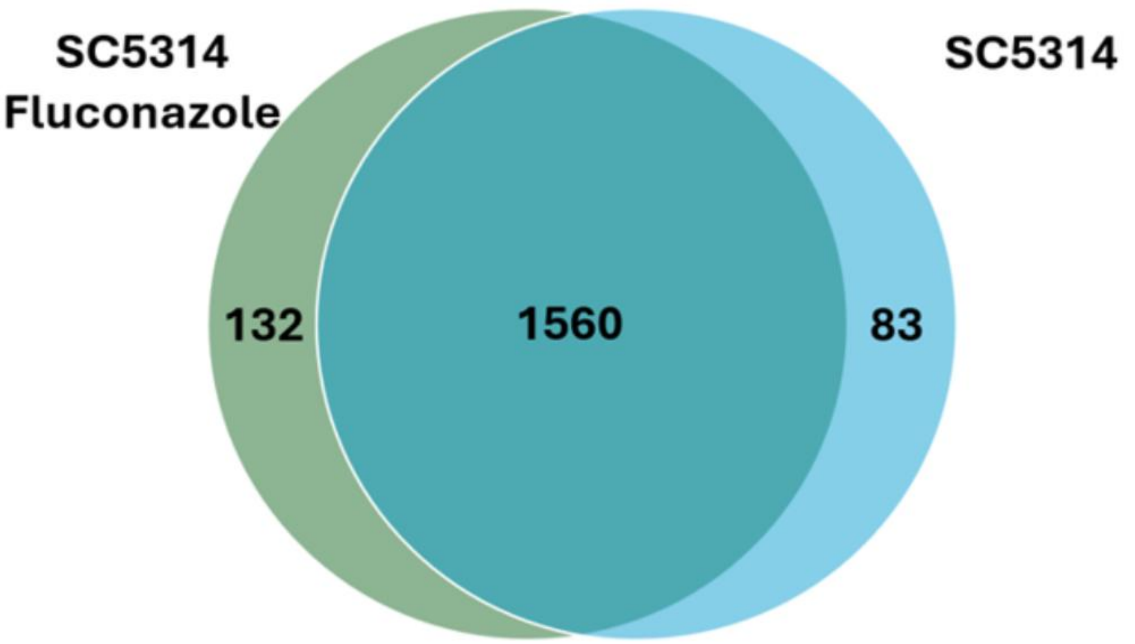


c)

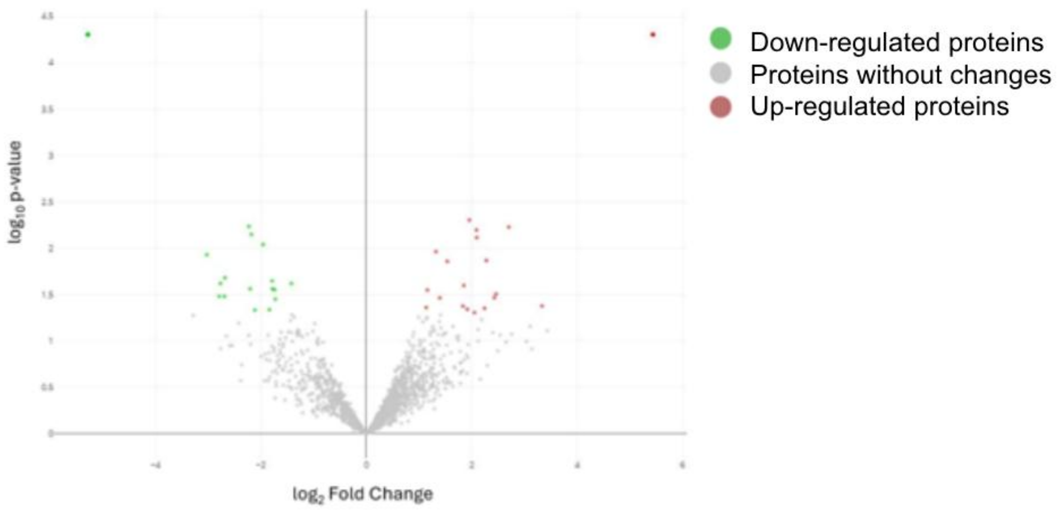


d)

| Condition | Quantified | Increased | Decreased |
|---|---|---|---|
| SC5314 Fluco / SC5314 | 1774 | 218 | 134 |

e)

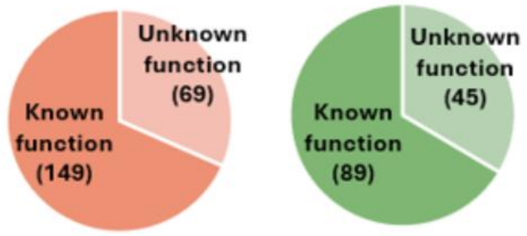

### Supplementary figure 3

a)

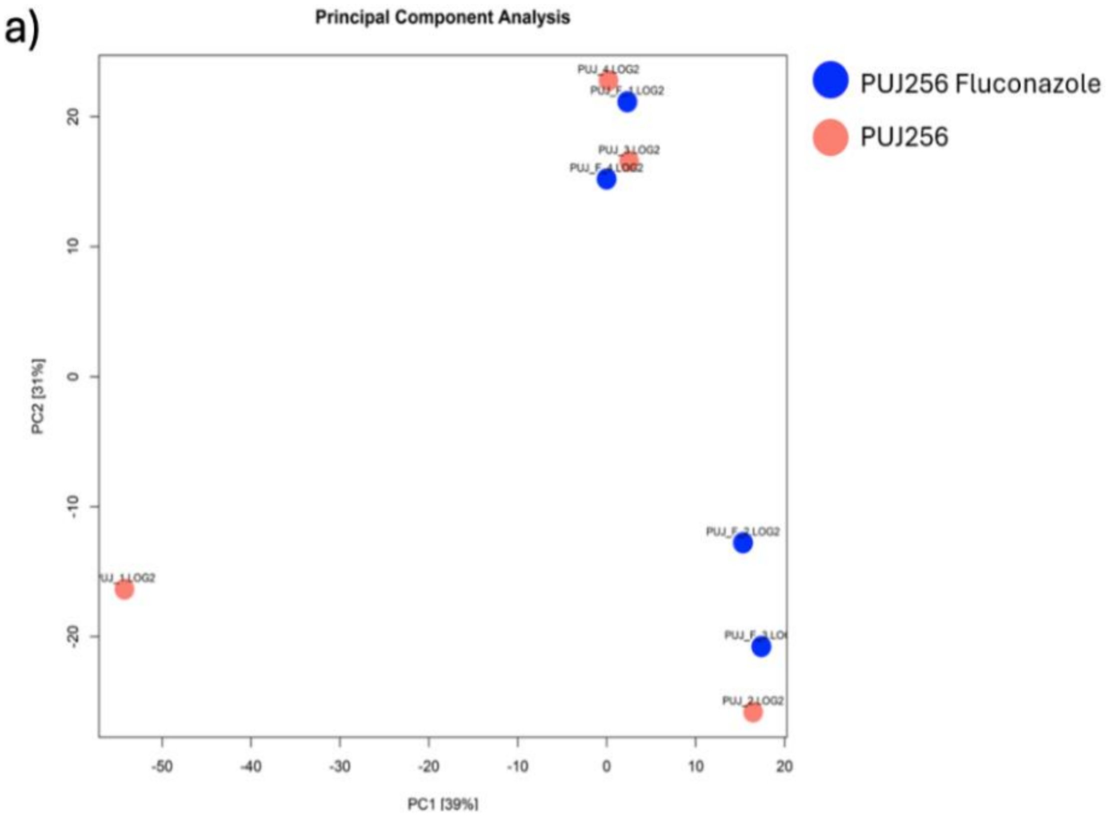


b)

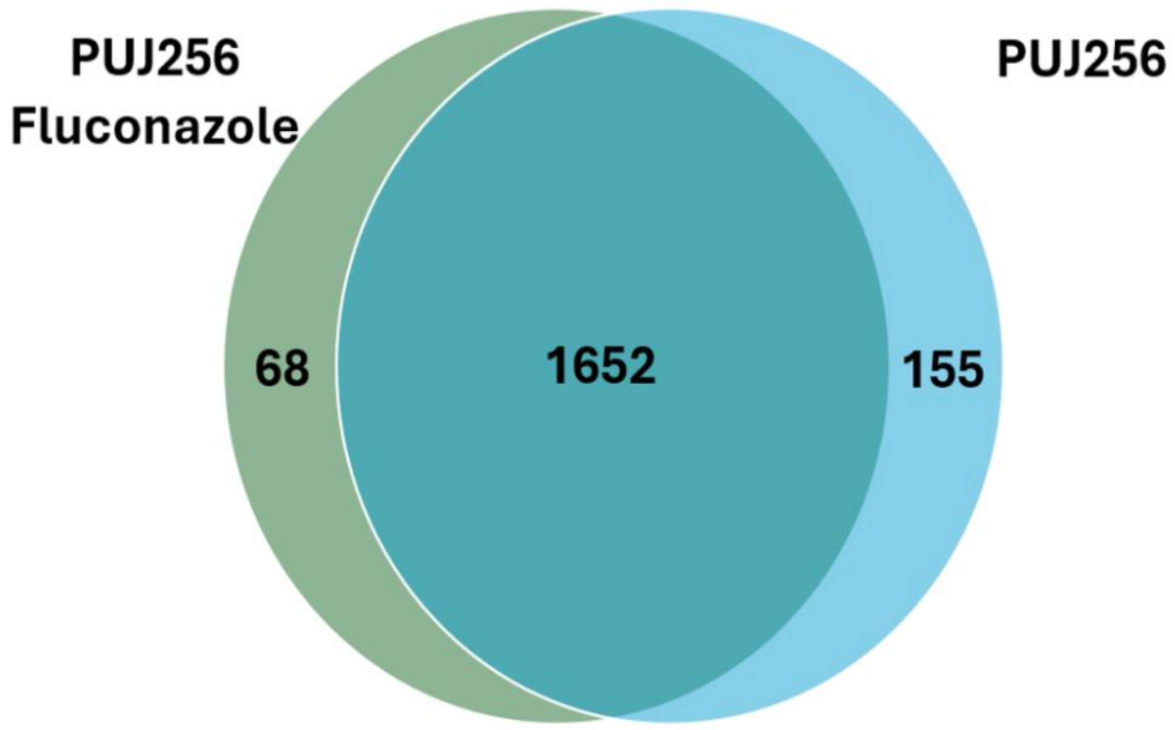


c)

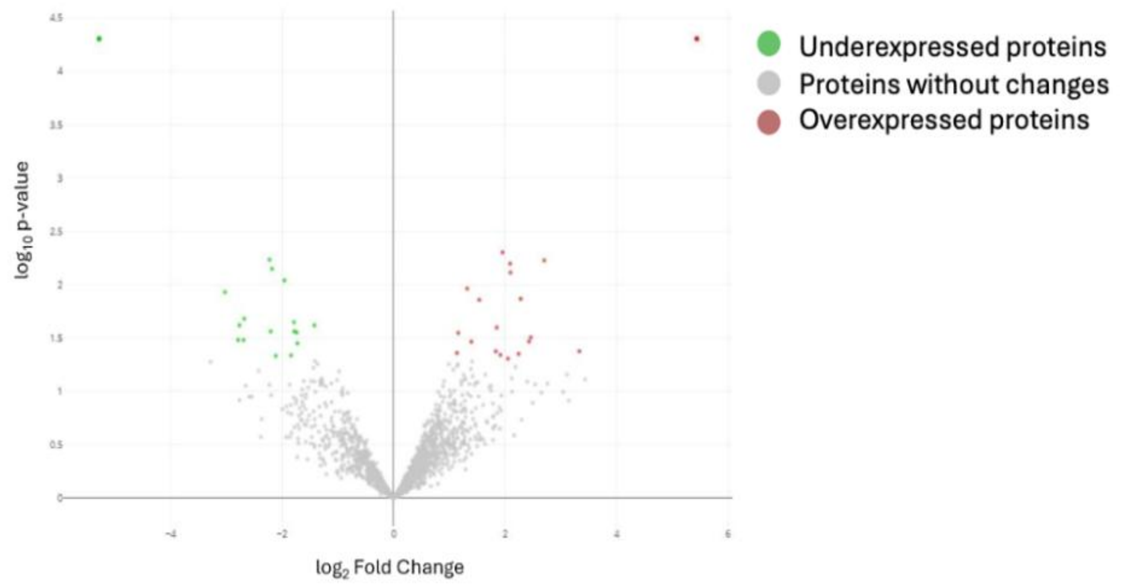


d)

| Condition | Quantified | Increased | Decreased |
|---|---|---|---|
| PUJ256 Fluco / PUJ256 | 1874 | 86 | 171 |

e)

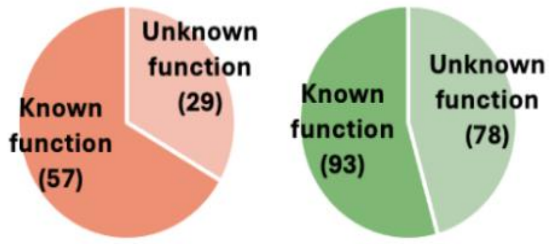

### Supplementary figure 4

a)

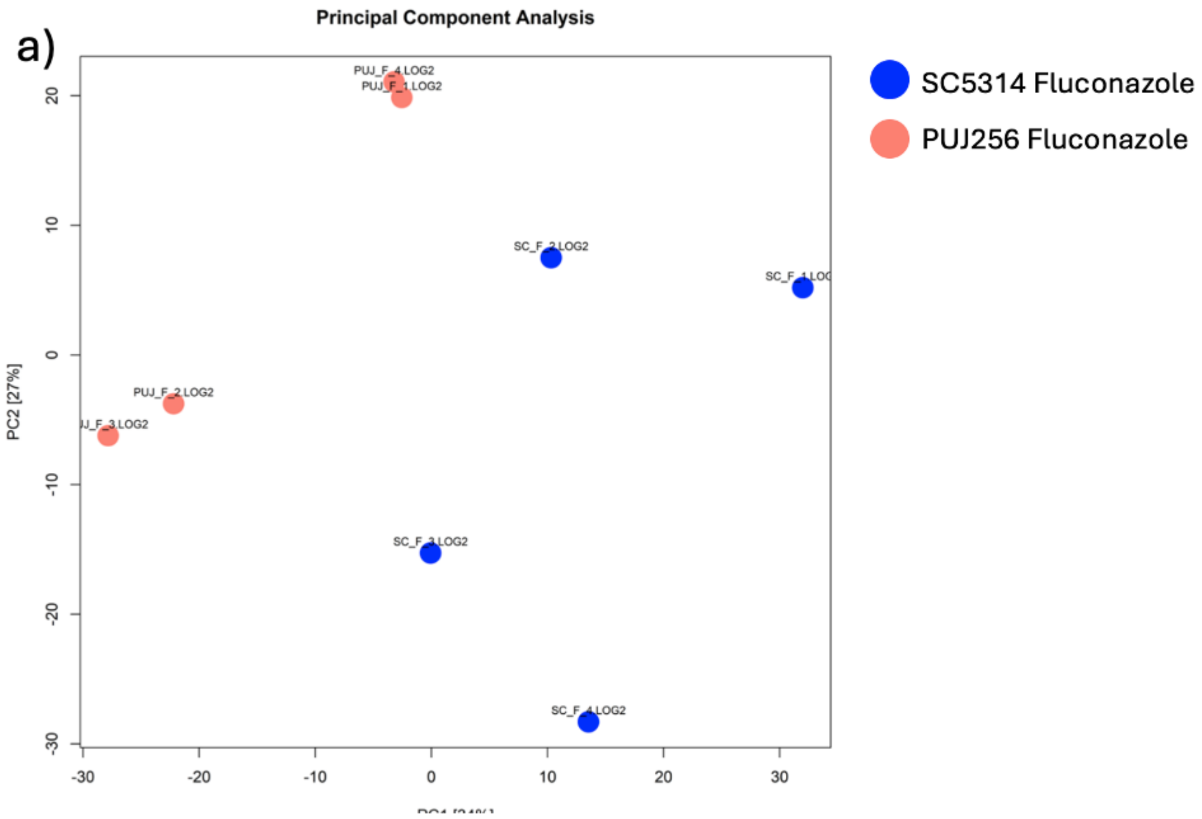


b)

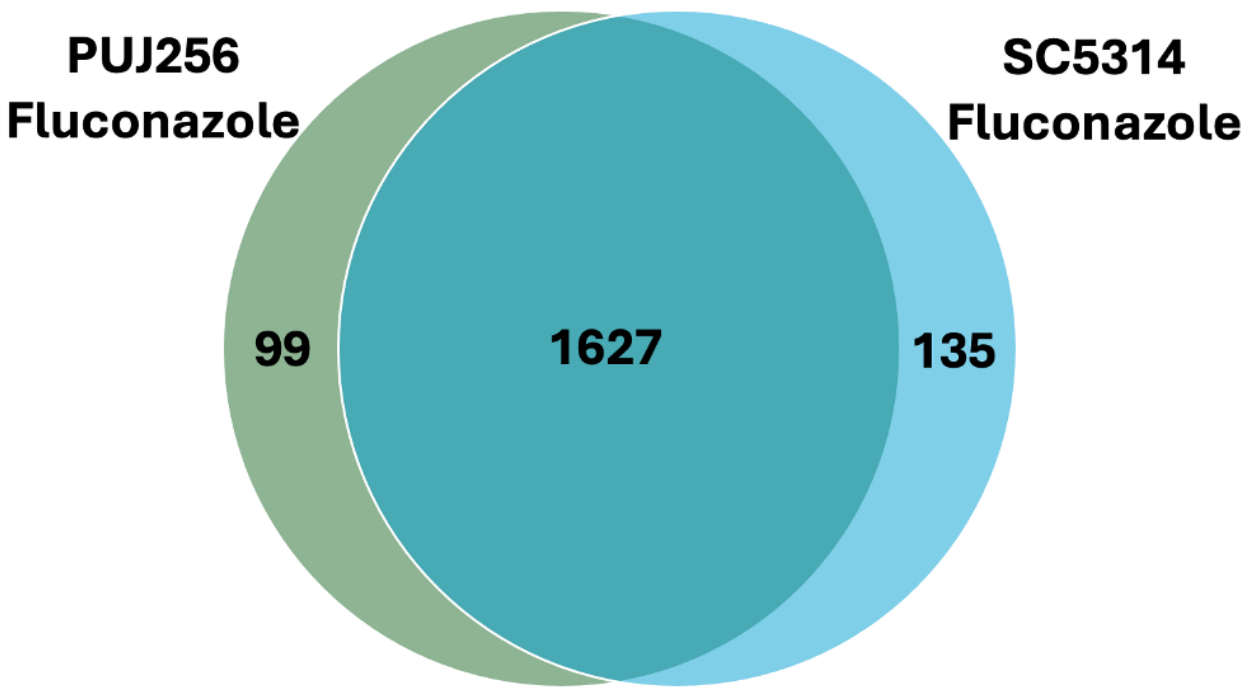


c)

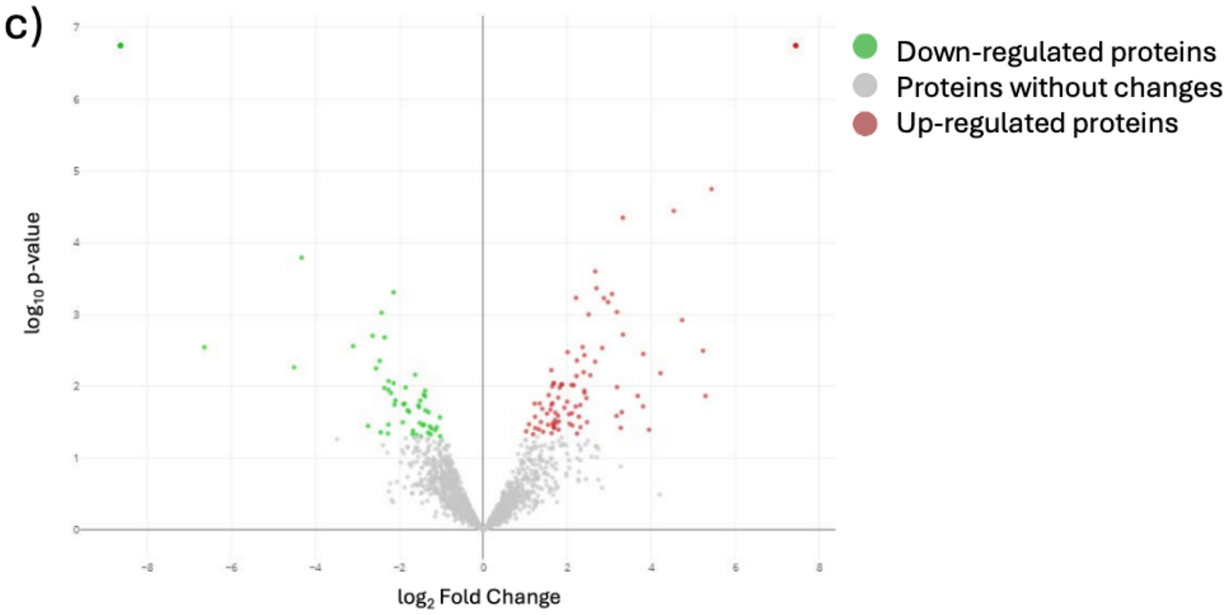


d)

| Condition | Quantified | Increased | Decreased |
|---|---|---|---|
| PUJ256 Fluco / SC5314 Fluco | 1860 | 253 | 261 |

e)

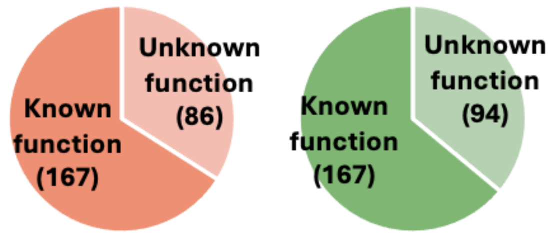
