## Supplemental Tables 1, 2, 3 and 4 for "Thirty years of fluconazole therapy selects an azole-resistant *Candida albicans* isolate with a pre-adapted physiological, metabolic and structural state"

Supplementary Material

### 1 Supplementary Tables

| **Supplementary table 1. List of proteins that significantly changed their abundance pvalue <0.05 between PUJ256 and SC5314** | | | | |
| --- | --- | --- | --- | --- |
| **Protein** | | **log2FC** | | **p.value** |
| ACS1 | | 8.632132565 | | 5.43834E-07 |
| ADE4 | | 8.632132565 | | 5.43834E-07 |
| AFP99 | | 8.632132565 | | 5.43834E-07 |
| ANP1 | | 8.632132565 | | 5.43834E-07 |
| APM1 | | 8.632132565 | | 5.43834E-07 |
| ARC19 | | 8.632132565 | | 5.43834E-07 |
| ARF3 | | 8.632132565 | | 5.43834E-07 |
| ATO2 | | 8.632132565 | | 5.43834E-07 |
| BNA4 | | 8.632132565 | | 5.43834E-07 |
| CCE1 | | 8.632132565 | | 5.43834E-07 |
| CCT7 | | 8.632132565 | | 5.43834E-07 |
| CEF1 | | 8.632132565 | | 5.43834E-07 |
| COX1 | | 8.632132565 | | 5.43834E-07 |
| COX2 | | 8.632132565 | | 5.43834E-07 |
| CTN3 | | 8.632132565 | | 5.43834E-07 |
| CYR1 | | 8.632132565 | | 5.43834E-07 |
| DAO1 | | 8.632132565 | | 5.43834E-07 |
| DRG1 | | 8.632132565 | | 5.43834E-07 |
| ECM21 | | 8.632132565 | | 5.43834E-07 |
| EPL1 | | 8.632132565 | | 5.43834E-07 |
| ERG11 | | 8.632132565 | | 5.43834E-07 |
| ERG2 | | 8.632132565 | | 5.43834E-07 |
| ERG3 | | 8.632132565 | | 5.43834E-07 |
| FLC2 | | 8.632132565 | | 5.43834E-07 |
| FMP45 | | 8.632132565 | | 5.43834E-07 |
| FOL1 | | 8.632132565 | | 5.43834E-07 |
| GCN5 | | 8.632132565 | | 5.43834E-07 |
| GPX3 | | 8.632132565 | | 5.43834E-07 |
| GTT12 | | 8.632132565 | | 5.43834E-07 |
| HPA2 | | 8.632132565 | | 5.43834E-07 |
| ISN1 | | 8.632132565 | | 5.43834E-07 |
| IST1 | | 8.632132565 | | 5.43834E-07 |
| MED20 | | 8.632132565 | | 5.43834E-07 |
| MET3 | | 8.632132565 | | 5.43834E-07 |
| MNN9 | | 8.632132565 | | 5.43834E-07 |
| MNT2 | | 8.632132565 | | 5.43834E-07 |
| NAG6 | | 8.632132565 | | 5.43834E-07 |
| NEP1 | | 8.632132565 | | 5.43834E-07 |
| OCA1 | | 8.632132565 | | 5.43834E-07 |
| ORC1 | | 8.632132565 | | 5.43834E-07 |
| orf19.102 | | 8.632132565 | | 5.43834E-07 |
| orf19.1034 | | 8.632132565 | | 5.43834E-07 |
| orf19.1054 | | 8.632132565 | | 5.43834E-07 |
| orf19.1057 | | 8.632132565 | | 5.43834E-07 |
| orf19.1113 | | 8.632132565 | | 5.43834E-07 |
| orf19.1114 | | 8.632132565 | | 5.43834E-07 |
| orf19.1179 | | 8.632132565 | | 5.43834E-07 |
| orf19.1417 | | 8.632132565 | | 5.43834E-07 |
| orf19.1480 | | 8.632132565 | | 5.43834E-07 |
| orf19.1584 | | 8.632132565 | | 5.43834E-07 |
| orf19.1589.1 | | 8.632132565 | | 5.43834E-07 |
| orf19.1608 | | 8.632132565 | | 5.43834E-07 |
| orf19.1618.1 | | 8.632132565 | | 5.43834E-07 |
| orf19.1625 | | 8.632132565 | | 5.43834E-07 |
| orf19.1682 | | 8.632132565 | | 5.43834E-07 |
| orf19.1761 | | 8.632132565 | | 5.43834E-07 |
| orf19.1772 | | 8.632132565 | | 5.43834E-07 |
| orf19.1856 | | 8.632132565 | | 5.43834E-07 |
| orf19.1873 | | 8.632132565 | | 5.43834E-07 |
| orf19.1888 | | 8.632132565 | | 5.43834E-07 |
| orf19.190 | | 8.632132565 | | 5.43834E-07 |
| orf19.1940 | | 8.632132565 | | 5.43834E-07 |
| orf19.2030 | | 8.632132565 | | 5.43834E-07 |
| orf19.216.1 | | 8.632132565 | | 5.43834E-07 |
| orf19.2228 | | 8.632132565 | | 5.43834E-07 |
| orf19.2318.1 | | 8.632132565 | | 5.43834E-07 |
| orf19.2346 | | 8.632132565 | | 5.43834E-07 |
| orf19.2400 | | 8.632132565 | | 5.43834E-07 |
| orf19.2472.1 | | 8.632132565 | | 5.43834E-07 |
| orf19.2473 | | 8.632132565 | | 5.43834E-07 |
| orf19.2485 | | 8.632132565 | | 5.43834E-07 |
| orf19.2564 | | 8.632132565 | | 5.43834E-07 |
| orf19.2673 | | 8.632132565 | | 5.43834E-07 |
| orf19.2749 | | 8.632132565 | | 5.43834E-07 |
| orf19.279 | | 8.632132565 | | 5.43834E-07 |
| orf19.2848 | | 8.632132565 | | 5.43834E-07 |
| orf19.2863 | | 8.632132565 | | 5.43834E-07 |
| orf19.2889 | | 8.632132565 | | 5.43834E-07 |
| orf19.2954 | | 8.632132565 | | 5.43834E-07 |
| orf19.2963 | | 8.632132565 | | 5.43834E-07 |
| orf19.2995 | | 8.632132565 | | 5.43834E-07 |
| orf19.3030 | | 8.632132565 | | 5.43834E-07 |
| orf19.3045 | | 8.632132565 | | 5.43834E-07 |
| orf19.3057 | | 8.632132565 | | 5.43834E-07 |
| orf19.3128 | | 8.632132565 | | 5.43834E-07 |
| orf19.3135 | | 8.632132565 | | 5.43834E-07 |
| orf19.3163 | | 8.632132565 | | 5.43834E-07 |
| orf19.3366.1 | | 8.632132565 | | 5.43834E-07 |
| orf19.3483 | | 8.632132565 | | 5.43834E-07 |
| orf19.3552 | | 8.632132565 | | 5.43834E-07 |
| orf19.3659 | | 8.632132565 | | 5.43834E-07 |
| orf19.3679 | | 8.632132565 | | 5.43834E-07 |
| orf19.3684 | | 8.632132565 | | 5.43834E-07 |
| orf19.3782.2 | | 8.632132565 | | 5.43834E-07 |
| orf19.3804 | | 8.632132565 | | 5.43834E-07 |
| orf19.3810 | | 8.632132565 | | 5.43834E-07 |
| orf19.3843 | | 8.632132565 | | 5.43834E-07 |
| orf19.3965 | | 8.632132565 | | 5.43834E-07 |
| orf19.3991 | | 8.632132565 | | 5.43834E-07 |
| orf19.4086 | | 8.632132565 | | 5.43834E-07 |
| orf19.409 | | 8.632132565 | | 5.43834E-07 |
| orf19.4132 | | 8.632132565 | | 5.43834E-07 |
| orf19.4262 | | 8.632132565 | | 5.43834E-07 |
| orf19.4358 | | 8.632132565 | | 5.43834E-07 |
| orf19.4365 | | 8.632132565 | | 5.43834E-07 |
| orf19.4370 | | 8.632132565 | | 5.43834E-07 |
| orf19.4382 | | 8.632132565 | | 5.43834E-07 |
| orf19.446.2 | | 8.632132565 | | 5.43834E-07 |
| orf19.4522 | | 8.632132565 | | 5.43834E-07 |
| orf19.4612 | | 8.632132565 | | 5.43834E-07 |
| orf19.4659 | | 8.632132565 | | 5.43834E-07 |
| orf19.491 | | 8.632132565 | | 5.43834E-07 |
| orf19.4914 | | 8.632132565 | | 5.43834E-07 |
| orf19.4947 | | 8.632132565 | | 5.43834E-07 |
| orf19.5079.1 | | 8.632132565 | | 5.43834E-07 |
| orf19.5134 | | 8.632132565 | | 5.43834E-07 |
| orf19.5238 | | 8.632132565 | | 5.43834E-07 |
| orf19.5249 | | 8.632132565 | | 5.43834E-07 |
| orf19.5411 | | 8.632132565 | | 5.43834E-07 |
| orf19.5418 | | 8.632132565 | | 5.43834E-07 |
| orf19.556 | | 8.632132565 | | 5.43834E-07 |
| orf19.5566 | | 8.632132565 | | 5.43834E-07 |
| orf19.5680 | | 8.632132565 | | 5.43834E-07 |
| orf19.5727 | | 8.632132565 | | 5.43834E-07 |
| orf19.5828 | | 8.632132565 | | 5.43834E-07 |
| orf19.5935 | | 8.632132565 | | 5.43834E-07 |
| orf19.6027 | | 8.632132565 | | 5.43834E-07 |
| orf19.6035 | | 8.632132565 | | 5.43834E-07 |
| orf19.6039 | | 8.632132565 | | 5.43834E-07 |
| orf19.6062 | | 8.632132565 | | 5.43834E-07 |
| orf19.6075 | | 8.632132565 | | 5.43834E-07 |
| orf19.6152 | | 8.632132565 | | 5.43834E-07 |
| orf19.6189 | | 8.632132565 | | 5.43834E-07 |
| orf19.6198.1 | | 8.632132565 | | 5.43834E-07 |
| orf19.6211 | | 8.632132565 | | 5.43834E-07 |
| orf19.6247.1 | | 8.632132565 | | 5.43834E-07 |
| orf19.6341 | | 8.632132565 | | 5.43834E-07 |
| orf19.642 | | 8.632132565 | | 5.43834E-07 |
| orf19.6474 | | 8.632132565 | | 5.43834E-07 |
| orf19.6552 | | 8.632132565 | | 5.43834E-07 |
| orf19.6565 | | 8.632132565 | | 5.43834E-07 |
| orf19.6607 | | 8.632132565 | | 5.43834E-07 |
| orf19.6639 | | 8.632132565 | | 5.43834E-07 |
| orf19.6732 | | 8.632132565 | | 5.43834E-07 |
| orf19.6769 | | 8.632132565 | | 5.43834E-07 |
| orf19.6788 | | 8.632132565 | | 5.43834E-07 |
| orf19.6939 | | 8.632132565 | | 5.43834E-07 |
| orf19.7067 | | 8.632132565 | | 5.43834E-07 |
| orf19.7088 | | 8.632132565 | | 5.43834E-07 |
| orf19.7183 | | 8.632132565 | | 5.43834E-07 |
| orf19.7202 | | 8.632132565 | | 5.43834E-07 |
| orf19.7260 | | 8.632132565 | | 5.43834E-07 |
| orf19.729 | | 8.632132565 | | 5.43834E-07 |
| orf19.7326 | | 8.632132565 | | 5.43834E-07 |
| orf19.7344 | | 8.632132565 | | 5.43834E-07 |
| orf19.7345 | | 8.632132565 | | 5.43834E-07 |
| orf19.7478 | | 8.632132565 | | 5.43834E-07 |
| orf19.7566 | | 8.632132565 | | 5.43834E-07 |
| orf19.757 | | 8.632132565 | | 5.43834E-07 |
| orf19.7604 | | 8.632132565 | | 5.43834E-07 |
| PAM16 | | 8.632132565 | | 5.43834E-07 |
| PAM18 | | 8.632132565 | | 5.43834E-07 |
| PDR17 | | 8.632132565 | | 5.43834E-07 |
| PEX11 | | 8.632132565 | | 5.43834E-07 |
| PKH3 | | 8.632132565 | | 5.43834E-07 |
| PMT1 | | 8.632132565 | | 5.43834E-07 |
| PRS5 | | 8.632132565 | | 5.43834E-07 |
| QCR8 | | 8.632132565 | | 5.43834E-07 |
| RAC1 | | 8.632132565 | | 5.43834E-07 |
| RFC2 | | 8.632132565 | | 5.43834E-07 |
| RHO2 | | 8.632132565 | | 5.43834E-07 |
| RIX7 | | 8.632132565 | | 5.43834E-07 |
| RMP1 | | 8.632132565 | | 5.43834E-07 |
| RNH35 | | 8.632132565 | | 5.43834E-07 |
| RRP42 | | 8.632132565 | | 5.43834E-07 |
| RTA2 | | 8.632132565 | | 5.43834E-07 |
| SAM4 | | 8.632132565 | | 5.43834E-07 |
| SAM51 | | 8.632132565 | | 5.43834E-07 |
| SAP2 | | 8.632132565 | | 5.43834E-07 |
| SAP4 | | 8.632132565 | | 5.43834E-07 |
| SCO1 | | 8.632132565 | | 5.43834E-07 |
| SFT1 | | 8.632132565 | | 5.43834E-07 |
| SLN1 | | 8.632132565 | | 5.43834E-07 |
| SNF1 | | 8.632132565 | | 5.43834E-07 |
| SPC2 | | 8.632132565 | | 5.43834E-07 |
| SPT7 | | 8.632132565 | | 5.43834E-07 |
| SRP54 | | 8.632132565 | | 5.43834E-07 |
| SUN41 | | 8.632132565 | | 5.43834E-07 |
| SUR2 | | 8.632132565 | | 5.43834E-07 |
| TIM22 | | 8.632132565 | | 5.43834E-07 |
| TOM7 | | 8.632132565 | | 5.43834E-07 |
| UBP1 | | 8.632132565 | | 5.43834E-07 |
| VMA5 | | 8.632132565 | | 5.43834E-07 |
| YIM1 | | 8.632132565 | | 5.43834E-07 |
| YPT72 | | 8.632132565 | | 5.43834E-07 |
| YTA6 | | 8.632132565 | | 5.43834E-07 |
| SER2 | | 6.632132565 | | 0.002350022 |
| orf19.6306 | | 5.822661904 | | 0.002413891 |
| GAL10 | | 5.113936159 | | 0.018788533 |
| MDH1 | | 4.899232191 | | 5.43834E-05 |
| LEU1 | | 4.772226385 | | 5.8248E-05 |
| HMT1 | | 4.543722233 | | 0.001257142 |
| GEA2 | | 4.46559307 | | 0.0379769 |
| DAP2 | | 4.020118279 | | 0.01517559 |
| orf19.2737 | | 3.867131306 | | 0.021426588 |
| orf19.3307 | | 3.756636181 | | 0.011197401 |
| HYR1 | | 3.746590095 | | 0.033671846 |
| ECM38 | | 3.715138231 | | 0.000959736 |
| orf19.7531 | | 3.596095479 | | 0.00109001 |
| AGO1 | | 3.55596599 | | 0.025224196 |
| orf19.5987 | | 3.474780459 | | 0.013496935 |
| OYE23 | | 3.467733111 | | 0.001333915 |
| orf19.639 | | 3.312732076 | | 0.010265489 |
| CYB2 | | 3.289586489 | | 0.032412386 |
| CIT1 | | 3.13458811 | | 0.025243913 |
| IPK2 | | 3.09093239 | | 0.002731259 |
| MTR10 | | 3.017590189 | | 0.040542779 |
| orf19.2278 | | 2.979695481 | | 0.03980444 |
| orf19.2965 | | 2.969565447 | | 0.024720977 |
| orf19.3787 | | 2.966607144 | | 0.034080288 |
| orf19.6604 | | 2.963141907 | | 0.002998924 |
| orf19.2853 | | 2.952819262 | | 0.03275488 |
| PHA2 | | 2.939068569 | | 0.001828703 |
| orf19.6658 | | 2.911441194 | | 0.007438616 |
| ACS2 | | 2.876776967 | | 0.006239563 |
| MEX67 | | 2.861211895 | | 0.003911355 |
| ARP9 | | 2.846602546 | | 0.017527148 |
| orf19.3292 | | 2.833288976 | | 0.023077532 |
| orf19.3312 | | 2.703606588 | | 0.024885164 |
| orf19.5322 | | 2.702300921 | | 0.01565773 |
| BUD23 | | 2.699625399 | | 0.016104248 |
| orf19.6271 | | 2.627183901 | | 0.005951739 |
| HIS7 | | 2.621898988 | | 0.005618932 |
| orf19.5077 | | 2.616720913 | | 0.039099457 |
| orf19.512 | | 2.574645423 | | 0.002210638 |
| orf19.2607 | | 2.568342897 | | 0.030840593 |
| orf19.4850 | | 2.550005731 | | 0.014267459 |
| GDA1 | | 2.545244445 | | 0.007032074 |
| VPS28 | | 2.492504518 | | 0.026466774 |
| MAE1 | | 2.491612374 | | 0.022854497 |
| GFA1 | | 2.489800052 | | 0.035065782 |
| orf19.4904 | | 2.417972728 | | 0.009606866 |
| orf19.5783 | | 2.409721823 | | 0.01536631 |
| orf19.3235 | | 2.404794669 | | 0.04009998 |
| PTC1 | | 2.383688187 | | 0.012114329 |
| LEU4 | | 2.358563781 | | 0.019640295 |
| orf19.1355 | | 2.346893286 | | 0.009103555 |
| orf19.86 | | 2.322230029 | | 0.027734289 |
| orf19.5885 | | 2.296388066 | | 0.011524519 |
| orf19.5541 | | 2.243205215 | | 0.03945321 |
| APN1 | | 2.242139167 | | 0.011125757 |
| BMT1 | | 2.240961674 | | 0.006979003 |
| orf19.1026 | | 2.234353671 | | 0.010109259 |
| SOF1 | | 2.154107799 | | 0.037692998 |
| RBK1 | | 2.146157972 | | 0.040953816 |
| BOI2 | | 2.12795736 | | 0.02562363 |
| BMS1 | | 2.118247541 | | 0.043045591 |
| CAT2 | | 2.096920224 | | 0.042666953 |
| orf19.7357 | | 2.085816024 | | 0.001205611 |
| orf19.2041 | | 2.076503905 | | 0.018884876 |
| orf19.6054 | | 2.057799803 | | 0.033093347 |
| ARO4 | | 2.045912443 | | 0.024835417 |
| EBP7 | | 2.01435389 | | 0.033764005 |
| orf19.4888 | | 1.971643771 | | 0.005986228 |
| MIS11 | | 1.970323471 | | 0.033497327 |
| orf19.5090 | | 1.967305631 | | 0.023274328 |
| orf19.1267.1 | | 1.92681478 | | 0.008076206 |
| orf19.7199 | | 1.889735843 | | 0.014800249 |
| ARO2 | | 1.842702849 | | 0.006212813 |
| orf19.4520 | | 1.836223396 | | 0.012173304 |
| MDH1-3 | | 1.830402933 | | 0.003909744 |
| UGA11 | | 1.814976038 | | 0.039626419 |
| orf19.504 | | 1.787709065 | | 0.020600338 |
| SUB2 | | 1.778303274 | | 0.015100865 |
| orf19.6883 | | 1.756402426 | | 0.030892345 |
| orf19.6539 | | 1.737090029 | | 0.013441301 |
| orf19.3223.1 | | 1.726764101 | | 0.046066687 |
| WAL1 | | 1.722218149 | | 0.030375355 |
| orf19.2111 | | 1.665563095 | | 0.036317773 |
| GAD1 | | 1.665232675 | | 0.018260154 |
| UGP1 | | 1.65866652 | | 0.043701376 |
| SPL1 | | 1.646789355 | | 0.030631616 |
| NOP1 | | 1.645847847 | | 0.012001981 |
| orf19.3932 | | 1.564641631 | | 0.024292677 |
| ALT1 | | 1.551495151 | | 0.027135777 |
| IFM3 | | 1.542241437 | | 0.047840579 |
| orf19.6359 | | 1.540369729 | | 0.023890208 |
| SCS7 | | 1.475881002 | | 0.035506186 |
| TIF11 | | 1.445588373 | | 0.027494581 |
| orf19.4246 | | 1.428078963 | | 0.023654058 |
| orf19.4597 | | 1.426003934 | | 0.031029777 |
| ECM25 | | 1.413541624 | | 0.048558275 |
| LYS12 | | 1.391365182 | | 0.041564384 |
| orf19.5114 | | 1.357169928 | | 0.046551075 |
| AUT7 | | 1.349993437 | | 0.047391522 |
| orf19.967 | | 1.301565921 | | 0.046423694 |
| ERG6 | | 1.251226534 | | 0.024279133 |
| MET15 | | 1.084521103 | | 0.04810563 |
| orf19.3928 | | -1.02991309 | | 0.04834037 |
| CLC1 | | -1.08486309 | | 0.029223736 |
| LTV1 | | -1.0906103 | | 0.036609854 |
| SNF7 | | -1.11005144 | | 0.02661238 |
| orf19.3173 | | -1.11266377 | | 0.042863408 |
| TBF1 | | -1.13829496 | | 0.047235891 |
| orf19.6984 | | -1.17294237 | | 0.028171707 |
| orf19.259 | | -1.17404574 | | 0.019031054 |
| orf19.5642 | | -1.18211111 | | 0.027927614 |
| PTR22 | | -1.20249858 | | 0.030652634 |
| orf19.1374 | | -1.24320343 | | 0.038656284 |
| SGA1 | | -1.26162948 | | 0.039500896 |
| orf19.7034 | | -1.28635585 | | 0.035548965 |
| orf19.1588 | | -1.31449374 | | 0.010577663 |
| EFB1 | | -1.3172736 | | 0.04869264 |
| FCR3 | | -1.31731993 | | 0.037028548 |
| MPRL36 | | -1.31818846 | | 0.022516484 |
| URA4 | | -1.3294113 | | 0.040117047 |
| orf19.7459 | | -1.33986398 | | 0.037589861 |
| orf19.4878 | | -1.35447928 | | 0.029017587 |
| PHO4 | | -1.3794601 | | 0.039706596 |
| VMA8 | | -1.41045517 | | 0.023190253 |
| HTA2 | | -1.44157381 | | 0.039724562 |
| KEX2 | | -1.44396572 | | 0.042431753 |
| orf19.2131 | | -1.46475525 | | 0.022279813 |
| CNB1 | | -1.46882654 | | 0.047432445 |
| VPS27 | | -1.47826426 | | 0.048728586 |
| orf19.934 | | -1.50262528 | | 0.011152659 |
| orf19.4172 | | -1.52730938 | | 0.011079619 |
| DQD1 | | -1.54020445 | | 0.045263685 |
| orf19.2426 | | -1.64877534 | | 0.031850116 |
| PRX1 | | -1.68546948 | | 0.007948557 |
| orf19.5609 | | -1.74168496 | | 0.006718545 |
| CDC19 | | -1.76744221 | | 0.040265438 |
| MCM3 | | -1.7844724 | | 0.023463656 |
| orf19.7341 | | -1.7878332 | | 0.006110847 |
| orf19.1137 | | -1.79938467 | | 0.028960484 |
| PEX14 | | -1.80396496 | | 0.024524647 |
| orf19.1815 | | -1.80994489 | | 0.031869926 |
| COI1 | | -1.81119033 | | 0.005160603 |
| HMS1 | | -1.82491867 | | 0.013134312 |
| RIB4 | | -1.85248912 | | 0.008131682 |
| MET6 | | -1.8938696 | | 0.027062667 |
| orf19.3887 | | -1.92469867 | | 0.037191185 |
| WOR3 | | -1.98969968 | | 0.007303671 |
| orf19.692 | | -2.04077101 | | 0.004543078 |
| LYS1 | | -2.10577009 | | 0.000796771 |
| orf19.7006 | | -2.11539725 | | 0.029249978 |
| HNT1 | | -2.12830977 | | 0.041285667 |
| orf19.7580 | | -2.12934017 | | 0.00271162 |
| NAM2 | | -2.14030893 | | 0.049915005 |
| SMT3 | | -2.15584898 | | 0.045236709 |
| GCV3 | | -2.19079527 | | 0.027188125 |
| orf19.1483 | | -2.20346613 | | 0.003746644 |
| HRT2 | | -2.29650531 | | 0.01080287 |
| orf19.1285 | | -2.3914517 | | 0.020165988 |
| orf19.4043 | | -2.52914392 | | 0.028592822 |
| orf19.1414 | | -2.60895554 | | 0.013822539 |
| orf19.2988 | | -2.6572942 | | 0.016907513 |
| CKS1 | | -2.9498215 | | 0.032298562 |
| orf19.3250 | | -3.13025001 | | 0.011514703 |
| orf19.4306 | | -3.39004292 | | 0.026044115 |
| SOD3 | | -3.42950656 | | 0.035417935 |
| AUR1 | | -3.67368045 | | 0.004925634 |
| orf19.6929 | | -3.96923186 | | 0.024604941 |
| orf19.5620 | | -4.3148478 | | 0.001682381 |
| ATS1 | | -4.65223479 | | 0.002211399 |
| ADP1 | | -6.65223479 | | 5.43834E-07 |
| ATP17 | | -6.65223479 | | 5.43834E-07 |
| BMT6 | | -6.65223479 | | 5.43834E-07 |
| BUD31 | | -6.65223479 | | 5.43834E-07 |
| CCC1 | | -6.65223479 | | 5.43834E-07 |
| CHS2 | | -6.65223479 | | 5.43834E-07 |
| CLB4 | | -6.65223479 | | 5.43834E-07 |
| CMK1 | | -6.65223479 | | 5.43834E-07 |
| CSR1 | | -6.65223479 | | 5.43834E-07 |
| CTM1 | | -6.65223479 | | 5.43834E-07 |
| DAD4 | | -6.65223479 | | 5.43834E-07 |
| DBP7 | | -6.65223479 | | 5.43834E-07 |
| DIP5 | | -6.65223479 | | 5.43834E-07 |
| DNM1 | | -6.65223479 | | 5.43834E-07 |
| ECM7 | | -6.65223479 | | 5.43834E-07 |
| GPX2 | | -6.65223479 | | 5.43834E-07 |
| GUT1 | | -6.65223479 | | 5.43834E-07 |
| GYP2 | | -6.65223479 | | 5.43834E-07 |
| HDA1 | | -6.65223479 | | 5.43834E-07 |
| HYS2 | | -6.65223479 | | 5.43834E-07 |
| IHD1 | | -6.65223479 | | 5.43834E-07 |
| IRS4 | | -6.65223479 | | 5.43834E-07 |
| ITR1 | | -6.65223479 | | 5.43834E-07 |
| MIR1 | | -6.65223479 | | 5.43834E-07 |
| MIT1 | | -6.65223479 | | 5.43834E-07 |
| MNL1 | | -6.65223479 | | 5.43834E-07 |
| MRPL19 | | -6.65223479 | | 5.43834E-07 |
| MUP1 | | -6.65223479 | | 5.43834E-07 |
| NMD5 | | -6.65223479 | | 5.43834E-07 |
| orf19.100 | | -6.65223479 | | 5.43834E-07 |
| orf19.1117 | | -6.65223479 | | 5.43834E-07 |
| orf19.1297 | | -6.65223479 | | 5.43834E-07 |
| orf19.1525 | | -6.65223479 | | 5.43834E-07 |
| orf19.1668 | | -6.65223479 | | 5.43834E-07 |
| orf19.1675 | | -6.65223479 | | 5.43834E-07 |
| orf19.1840 | | -6.65223479 | | 5.43834E-07 |
| orf19.1871 | | -6.65223479 | | 5.43834E-07 |
| orf19.1975 | | -6.65223479 | | 5.43834E-07 |
| orf19.2017 | | -6.65223479 | | 5.43834E-07 |
| orf19.2728 | | -6.65223479 | | 5.43834E-07 |
| orf19.2770 | | -6.65223479 | | 5.43834E-07 |
| orf19.3226 | | -6.65223479 | | 5.43834E-07 |
| orf19.3418 | | -6.65223479 | | 5.43834E-07 |
| orf19.3556 | | -6.65223479 | | 5.43834E-07 |
| orf19.3606 | | -6.65223479 | | 5.43834E-07 |
| orf19.3615 | | -6.65223479 | | 5.43834E-07 |
| orf19.3689 | | -6.65223479 | | 5.43834E-07 |
| orf19.3874 | | -6.65223479 | | 5.43834E-07 |
| orf19.3932.1 | | -6.65223479 | | 5.43834E-07 |
| orf19.4169 | | -6.65223479 | | 5.43834E-07 |
| orf19.4724 | | -6.65223479 | | 5.43834E-07 |
| orf19.4749 | | -6.65223479 | | 5.43834E-07 |
| orf19.4787 | | -6.65223479 | | 5.43834E-07 |
| orf19.4824 | | -6.65223479 | | 5.43834E-07 |
| orf19.4893 | | -6.65223479 | | 5.43834E-07 |
| orf19.5092 | | -6.65223479 | | 5.43834E-07 |
| orf19.515 | | -6.65223479 | | 5.43834E-07 |
| orf19.5160 | | -6.65223479 | | 5.43834E-07 |
| orf19.5269 | | -6.65223479 | | 5.43834E-07 |
| orf19.5376 | | -6.65223479 | | 5.43834E-07 |
| orf19.5555 | | -6.65223479 | | 5.43834E-07 |
| orf19.5684.1 | | -6.65223479 | | 5.43834E-07 |
| orf19.5688 | | -6.65223479 | | 5.43834E-07 |
| orf19.5905 | | -6.65223479 | | 5.43834E-07 |
| orf19.5929 | | -6.65223479 | | 5.43834E-07 |
| orf19.6023 | | -6.65223479 | | 5.43834E-07 |
| orf19.6185 | | -6.65223479 | | 5.43834E-07 |
| orf19.6233 | | -6.65223479 | | 5.43834E-07 |
| orf19.6342 | | -6.65223479 | | 5.43834E-07 |
| orf19.6487 | | -6.65223479 | | 5.43834E-07 |
| orf19.6597 | | -6.65223479 | | 5.43834E-07 |
| orf19.7058 | | -6.65223479 | | 5.43834E-07 |
| orf19.7316 | | -6.65223479 | | 5.43834E-07 |
| orf19.7425 | | -6.65223479 | | 5.43834E-07 |
| PEX6 | | -6.65223479 | | 5.43834E-07 |
| PGA53 | | -6.65223479 | | 5.43834E-07 |
| PHO91 | | -6.65223479 | | 5.43834E-07 |
| PLB1 | | -6.65223479 | | 5.43834E-07 |
| PWP2 | | -6.65223479 | | 5.43834E-07 |
| PZF1 | | -6.65223479 | | 5.43834E-07 |
| RAD9 | | -6.65223479 | | 5.43834E-07 |
| SCH9 | | -6.65223479 | | 5.43834E-07 |
| SCP1 | | -6.65223479 | | 5.43834E-07 |
| SEN1 | | -6.65223479 | | 5.43834E-07 |
| SLD5 | | -6.65223479 | | 5.43834E-07 |
| SLU7 | | -6.65223479 | | 5.43834E-07 |
| SMI1B | | -6.65223479 | | 5.43834E-07 |
| SPT20 | | -6.65223479 | | 5.43834E-07 |
| STP4 | | -6.65223479 | | 5.43834E-07 |
| TFC4 | | -6.65223479 | | 5.43834E-07 |
| TLO11 | | -6.65223479 | | 5.43834E-07 |
| TPO4 | | -6.65223479 | | 5.43834E-07 |
| TRP4 | | -6.65223479 | | 5.43834E-07 |
| WSC4 | | -6.65223479 | | 5.43834E-07 |
| YBL053 | | -6.65223479 | | 5.43834E-07 |
| YME1 | | -6.65223479 | | 5.43834E-07 |
| **Supplementary table 2. List of proteins that significantly changed their abundance pvalue <0.05 between SC5314_FLU and SC5314** | | | | |
| **Protein** | **log2FC** | | **p.value** | |
| ACS1 | 7.428889072 | | 1.78E-07 | |
| ADE4 | 7.428889072 | | 1.78E-07 | |
| AFP99 | 7.428889072 | | 1.78E-07 | |
| AGP2 | 7.428889072 | | 1.78E-07 | |
| APM1 | 7.428889072 | | 1.78E-07 | |
| ATO2 | 7.428889072 | | 1.78E-07 | |
| BMT4 | 7.428889072 | | 1.78E-07 | |
| BUR2 | 7.428889072 | | 1.78E-07 | |
| CCE1 | 7.428889072 | | 1.78E-07 | |
| CCR4 | 7.428889072 | | 1.78E-07 | |
| CDC42 | 7.428889072 | | 1.78E-07 | |
| CTN3 | 7.428889072 | | 1.78E-07 | |
| CYR1 | 7.428889072 | | 1.78E-07 | |
| DAG7 | 7.428889072 | | 1.78E-07 | |
| DUN1 | 7.428889072 | | 1.78E-07 | |
| ECM21 | 7.428889072 | | 1.78E-07 | |
| ENA2 | 7.428889072 | | 1.78E-07 | |
| EPL1 | 7.428889072 | | 1.78E-07 | |
| ERG11 | 7.428889072 | | 1.78E-07 | |
| ERG3 | 7.428889072 | | 1.78E-07 | |
| EXO84 | 7.428889072 | | 1.78E-07 | |
| FCR1 | 7.428889072 | | 1.78E-07 | |
| FLC2 | 7.428889072 | | 1.78E-07 | |
| FMP45 | 7.428889072 | | 1.78E-07 | |
| FOL1 | 7.428889072 | | 1.78E-07 | |
| GCN5 | 7.428889072 | | 1.78E-07 | |
| GTT12 | 7.428889072 | | 1.78E-07 | |
| GYP7 | 7.428889072 | | 1.78E-07 | |
| HGT2 | 7.428889072 | | 1.78E-07 | |
| IFR1 | 7.428889072 | | 1.78E-07 | |
| ISN1 | 7.428889072 | | 1.78E-07 | |
| IST1 | 7.428889072 | | 1.78E-07 | |
| JEM1 | 7.428889072 | | 1.78E-07 | |
| MNN9 | 7.428889072 | | 1.78E-07 | |
| MNT2 | 7.428889072 | | 1.78E-07 | |
| MRV2 | 7.428889072 | | 1.78E-07 | |
| NEP1 | 7.428889072 | | 1.78E-07 | |
| NOC2 | 7.428889072 | | 1.78E-07 | |
| ORC1 | 7.428889072 | | 1.78E-07 | |
| orf19.1057 | 7.428889072 | | 1.78E-07 | |
| orf19.1179 | 7.428889072 | | 1.78E-07 | |
| orf19.1584 | 7.428889072 | | 1.78E-07 | |
| orf19.1589.1 | 7.428889072 | | 1.78E-07 | |
| orf19.1608 | 7.428889072 | | 1.78E-07 | |
| orf19.1764 | 7.428889072 | | 1.78E-07 | |
| orf19.1772 | 7.428889072 | | 1.78E-07 | |
| orf19.1800 | 7.428889072 | | 1.78E-07 | |
| orf19.1856 | 7.428889072 | | 1.78E-07 | |
| orf19.1888 | 7.428889072 | | 1.78E-07 | |
| orf19.190 | 7.428889072 | | 1.78E-07 | |
| orf19.2030 | 7.428889072 | | 1.78E-07 | |
| orf19.213 | 7.428889072 | | 1.78E-07 | |
| orf19.2346 | 7.428889072 | | 1.78E-07 | |
| orf19.2473 | 7.428889072 | | 1.78E-07 | |
| orf19.2476 | 7.428889072 | | 1.78E-07 | |
| orf19.2485 | 7.428889072 | | 1.78E-07 | |
| orf19.2564 | 7.428889072 | | 1.78E-07 | |
| orf19.2663 | 7.428889072 | | 1.78E-07 | |
| orf19.2838 | 7.428889072 | | 1.78E-07 | |
| orf19.2848 | 7.428889072 | | 1.78E-07 | |
| orf19.2889 | 7.428889072 | | 1.78E-07 | |
| orf19.3045 | 7.428889072 | | 1.78E-07 | |
| orf19.3135 | 7.428889072 | | 1.78E-07 | |
| orf19.3163 | 7.428889072 | | 1.78E-07 | |
| orf19.3483 | 7.428889072 | | 1.78E-07 | |
| orf19.3679 | 7.428889072 | | 1.78E-07 | |
| orf19.3684 | 7.428889072 | | 1.78E-07 | |
| orf19.3804 | 7.428889072 | | 1.78E-07 | |
| orf19.3810 | 7.428889072 | | 1.78E-07 | |
| orf19.3831 | 7.428889072 | | 1.78E-07 | |
| orf19.4117 | 7.428889072 | | 1.78E-07 | |
| orf19.4262 | 7.428889072 | | 1.78E-07 | |
| orf19.4292 | 7.428889072 | | 1.78E-07 | |
| orf19.4382 | 7.428889072 | | 1.78E-07 | |
| orf19.4522 | 7.428889072 | | 1.78E-07 | |
| orf19.4589 | 7.428889072 | | 1.78E-07 | |
| orf19.4659 | 7.428889072 | | 1.78E-07 | |
| orf19.491 | 7.428889072 | | 1.78E-07 | |
| orf19.5247 | 7.428889072 | | 1.78E-07 | |
| orf19.5249 | 7.428889072 | | 1.78E-07 | |
| orf19.5418 | 7.428889072 | | 1.78E-07 | |
| orf19.5566 | 7.428889072 | | 1.78E-07 | |
| orf19.5680 | 7.428889072 | | 1.78E-07 | |
| orf19.5727 | 7.428889072 | | 1.78E-07 | |
| orf19.580 | 7.428889072 | | 1.78E-07 | |
| orf19.5814.1 | 7.428889072 | | 1.78E-07 | |
| orf19.5828 | 7.428889072 | | 1.78E-07 | |
| orf19.5852 | 7.428889072 | | 1.78E-07 | |
| orf19.6075 | 7.428889072 | | 1.78E-07 | |
| orf19.6152 | 7.428889072 | | 1.78E-07 | |
| orf19.6211 | 7.428889072 | | 1.78E-07 | |
| orf19.6396 | 7.428889072 | | 1.78E-07 | |
| orf19.6484 | 7.428889072 | | 1.78E-07 | |
| orf19.6552 | 7.428889072 | | 1.78E-07 | |
| orf19.6581 | 7.428889072 | | 1.78E-07 | |
| orf19.6601 | 7.428889072 | | 1.78E-07 | |
| orf19.6639 | 7.428889072 | | 1.78E-07 | |
| orf19.6769 | 7.428889072 | | 1.78E-07 | |
| orf19.6788 | 7.428889072 | | 1.78E-07 | |
| orf19.6790 | 7.428889072 | | 1.78E-07 | |
| orf19.6939 | 7.428889072 | | 1.78E-07 | |
| orf19.7183 | 7.428889072 | | 1.78E-07 | |
| orf19.7194 | 7.428889072 | | 1.78E-07 | |
| orf19.729 | 7.428889072 | | 1.78E-07 | |
| orf19.7326 | 7.428889072 | | 1.78E-07 | |
| orf19.7345 | 7.428889072 | | 1.78E-07 | |
| orf19.757 | 7.428889072 | | 1.78E-07 | |
| PEX1 | 7.428889072 | | 1.78E-07 | |
| PHO89 | 7.428889072 | | 1.78E-07 | |
| PIKA | 7.428889072 | | 1.78E-07 | |
| PKH3 | 7.428889072 | | 1.78E-07 | |
| PMR1 | 7.428889072 | | 1.78E-07 | |
| PRS5 | 7.428889072 | | 1.78E-07 | |
| RAC1 | 7.428889072 | | 1.78E-07 | |
| RBE1 | 7.428889072 | | 1.78E-07 | |
| RIX7 | 7.428889072 | | 1.78E-07 | |
| RMP1 | 7.428889072 | | 1.78E-07 | |
| RNH35 | 7.428889072 | | 1.78E-07 | |
| RTA2 | 7.428889072 | | 1.78E-07 | |
| SAM4 | 7.428889072 | | 1.78E-07 | |
| SAP4 | 7.428889072 | | 1.78E-07 | |
| SNF1 | 7.428889072 | | 1.78E-07 | |
| SPT7 | 7.428889072 | | 1.78E-07 | |
| SRP54 | 7.428889072 | | 1.78E-07 | |
| SUN41 | 7.428889072 | | 1.78E-07 | |
| SUR2 | 7.428889072 | | 1.78E-07 | |
| TCC1 | 7.428889072 | | 1.78E-07 | |
| TOM7 | 7.428889072 | | 1.78E-07 | |
| TPS1 | 7.428889072 | | 1.78E-07 | |
| TRS33 | 7.428889072 | | 1.78E-07 | |
| VMA5 | 7.428889072 | | 1.78E-07 | |
| YTA6 | 7.428889072 | | 1.78E-07 | |
| XOG1 | 5.428889072 | | 1.78E-05 | |
| orf19.4423 | 5.284511649 | | 0.013619391 | |
| SEN1 | 5.225261945 | | 0.003202716 | |
| ENA21 | 4.730786173 | | 0.001195555 | |
| PMC1 | 4.531185693 | | 3.59E-05 | |
| RHO3 | 4.215027564 | | 0.006552643 | |
| ARG1 | 3.942850533 | | 0.040386487 | |
| DDR48 | 3.804533417 | | 0.003539026 | |
| RPT4 | 3.799234496 | | 0.019055439 | |
| orf19.5250 | 3.672574521 | | 0.013609901 | |
| orf19.6311 | 3.320827826 | | 0.001905206 | |
| PNG2 | 3.318764125 | | 4.46E-05 | |
| CYB2 | 3.295065946 | | 0.022986123 | |
| orf19.2737 | 3.271336617 | | 0.038105502 | |
| POT1 | 3.179208041 | | 0.010260454 | |
| KRE6 | 3.178049729 | | 0.00092103 | |
| GAL10 | 3.170402231 | | 0.025929993 | |
| orf19.5295 | 3.057938316 | | 0.000516494 | |
| WAL1 | 2.966700705 | | 0.000670224 | |
| orf19.2125 | 2.869209293 | | 0.00059023 | |
| PHM7 | 2.828099558 | | 0.002925661 | |
| orf19.3932 | 2.690574693 | | 0.000430463 | |
| PHR2 | 2.659126888 | | 0.000250742 | |
| BZZ1 | 2.658892568 | | 0.004548574 | |
| NCR1 | 2.546382569 | | 0.007015762 | |
| ERG6 | 2.506098414 | | 0.000997383 | |
| SUR7 | 2.467495092 | | 0.031581031 | |
| RHO1 | 2.452898402 | | 0.014608507 | |
| PST3 | 2.404332104 | | 0.003686693 | |
| APN1 | 2.404326959 | | 0.012322559 | |
| MDH1 | 2.402493539 | | 0.011505546 | |
| orf19.3310 | 2.384911054 | | 0.006361084 | |
| RCT1 | 2.359305231 | | 0.0028405 | |
| orf19.6658 | 2.30868298 | | 0.03720667 | |
| orf19.3213 | 2.305006616 | | 0.018264913 | |
| HYR1 | 2.272110117 | | 0.026505038 | |
| LYS22 | 2.229450023 | | 0.04564118 | |
| orf19.6979 | 2.225314169 | | 0.004378866 | |
| orf19.4476 | 2.216368724 | | 0.007201662 | |
| SAP9 | 2.20375442 | | 0.000586149 | |
| VPS21 | 2.197471952 | | 0.019114638 | |
| DAP1 | 2.149555408 | | 0.00966097 | |
| orf19.5525 | 2.117338112 | | 0.035180241 | |
| SRO77 | 2.106704822 | | 0.023744572 | |
| orf19.6637 | 2.09822358 | | 0.009583014 | |
| CSP37 | 2.05066155 | | 0.033206671 | |
| AMO1 | 2.039946692 | | 0.02454123 | |
| CRH11 | 2.002147485 | | 0.003339127 | |
| orf19.4520 | 1.991571376 | | 0.016400053 | |
| MYO5 | 1.925870813 | | 0.019897169 | |
| orf19.4246 | 1.87554564 | | 0.009413424 | |
| APR1 | 1.849701831 | | 0.009490885 | |
| SCS7 | 1.814636322 | | 0.010472724 | |
| orf19.2113 | 1.798139469 | | 0.03082048 | |
| HGT1 | 1.790265665 | | 0.040141227 | |
| orf19.3848 | 1.781567078 | | 0.014393946 | |
| LYS4 | 1.774994412 | | 0.025721683 | |
| orf19.2111 | 1.733452968 | | 0.032598909 | |
| orf19.7085 | 1.712541921 | | 0.023365306 | |
| FRP3 | 1.704159495 | | 0.029567158 | |
| GDA1 | 1.684599721 | | 0.037591773 | |
| RET2 | 1.678375299 | | 0.038118508 | |
| CSH1 | 1.676074562 | | 0.009010299 | |
| PGA52 | 1.668879044 | | 0.009119713 | |
| orf19.6359 | 1.65861738 | | 0.034761028 | |
| GAR1 | 1.653956928 | | 0.032749137 | |
| PST1 | 1.642220987 | | 0.017145651 | |
| orf19.6554 | 1.640898803 | | 0.009997713 | |
| orf19.6816 | 1.621753969 | | 0.018043164 | |
| SEC3 | 1.621738444 | | 0.045132925 | |
| PRC2 | 1.615018401 | | 0.005959961 | |
| orf19.2296 | 1.597659344 | | 0.021193463 | |
| PMA1 | 1.553603778 | | 0.013285308 | |
| orf19.7531 | 1.538832628 | | 0.034413113 | |
| ARE2 | 1.514875684 | | 0.02414628 | |
| orf19.6148 | 1.423333374 | | 0.04313817 | |
| orf19.7357 | 1.402831549 | | 0.020587406 | |
| orf19.4888 | 1.36880817 | | 0.031413467 | |
| BGL2 | 1.339824087 | | 0.017337543 | |
| ECM331 | 1.322294368 | | 0.039935138 | |
| COX19 | 1.23338 | | 0.038188297 | |
| orf19.716 | 1.228090885 | | 0.026656244 | |
| TOS1 | 1.217010345 | | 0.017427978 | |
| LYS9 | 1.183158601 | | 0.046556692 | |
| PLB3 | 1.088564156 | | 0.033770607 | |
| orf19.5614 | 1.018206745 | | 0.04242945 | |
| orf19.1889 | -1.030025301 | | 0.04973683 | |
| orf19.1273 | -1.035326142 | | 0.027118135 | |
| LTV1 | -1.10858977 | | 0.037251711 | |
| orf19.6990 | -1.14218201 | | 0.040862365 | |
| orf19.3572.3 | -1.23591494 | | 0.038724805 | |
| RPL39 | -1.25090647 | | 0.046662587 | |
| CKS1 | -1.275174829 | | 0.036144608 | |
| PGA4 | -1.305129481 | | 0.02300771 | |
| orf19.3087.2 | -1.314996158 | | 0.044384062 | |
| orf19.4204 | -1.379631224 | | 0.021648664 | |
| CNB1 | -1.383087733 | | 0.011468205 | |
| orf19.2452 | -1.385904468 | | 0.013827533 | |
| orf19.1897 | -1.414315638 | | 0.034165457 | |
| DBP2 | -1.420142851 | | 0.013060354 | |
| MVD | -1.442952365 | | 0.035038366 | |
| orf19.5747 | -1.484605062 | | 0.049808954 | |
| RIP1 | -1.502841631 | | 0.015872088 | |
| RPS19A | -1.512417462 | | 0.032470484 | |
| orf19.7341 | -1.529707837 | | 0.019587116 | |
| PDX3 | -1.547201419 | | 0.018369397 | |
| KRE5 | -1.596572763 | | 0.047144151 | |
| WRS1 | -1.624181116 | | 0.006893757 | |
| GIS2 | -1.675680707 | | 0.041812829 | |
| SPE2 | -1.683225116 | | 0.04653011 | |
| HTS1 | -1.768897233 | | 0.022752143 | |
| MED9 | -1.802773058 | | 0.021565331 | |
| orf19.7580 | -1.84813246 | | 0.010389859 | |
| MED3 | -1.873757138 | | 0.017406521 | |
| FBA1 | -1.903821642 | | 0.017865909 | |
| orf19.2852 | -1.917510272 | | 0.031849298 | |
| RFG1 | -2.093500458 | | 0.015777869 | |
| orf19.5254 | -2.112645182 | | 0.017964726 | |
| FCR3 | -2.136117602 | | 0.00048903 | |
| CDC10 | -2.136198867 | | 0.009022657 | |
| RPL17B | -2.196256404 | | 0.012395863 | |
| TIM13 | -2.253872599 | | 0.008459244 | |
| RPL11 | -2.260036798 | | 0.011172547 | |
| orf19.1545 | -2.261293307 | | 0.034177034 | |
| RPL30 | -2.270341926 | | 0.045472788 | |
| RHR2 | -2.352556582 | | 0.002082487 | |
| orf19.6679 | -2.356008348 | | 0.010550177 | |
| orf19.1409.1 | -2.423577992 | | 0.000938589 | |
| GIN1 | -2.445162645 | | 0.043870657 | |
| RLI1 | -2.46673506 | | 0.004422775 | |
| ASM3 | -2.552851724 | | 0.005641991 | |
| BRE1 | -2.633492018 | | 0.001985059 | |
| orf19.5103 | -2.745737216 | | 0.03569348 | |
| ECE1 | -3.099412014 | | 0.002752056 | |
| NUP | -4.323941597 | | 0.000161117 | |
| orf19.6929 | -4.504846784 | | 0.005456655 | |
| RAT1 | -6.637705648 | | 0.0028556 | |
| AAH1 | -8.637705648 | | 1.78E-07 | |
| ABP140 | -8.637705648 | | 1.78E-07 | |
| ARC35 | -8.637705648 | | 1.78E-07 | |
| ATP17 | -8.637705648 | | 1.78E-07 | |
| BMT1 | -8.637705648 | | 1.78E-07 | |
| BMT6 | -8.637705648 | | 1.78E-07 | |
| CLB4 | -8.637705648 | | 1.78E-07 | |
| CSR1 | -8.637705648 | | 1.78E-07 | |
| CTA24 | -8.637705648 | | 1.78E-07 | |
| CTA8 | -8.637705648 | | 1.78E-07 | |
| CTF5 | -8.637705648 | | 1.78E-07 | |
| CTF8 | -8.637705648 | | 1.78E-07 | |
| CTM1 | -8.637705648 | | 1.78E-07 | |
| DBP7 | -8.637705648 | | 1.78E-07 | |
| DEF1 | -8.637705648 | | 1.78E-07 | |
| DIP2 | -8.637705648 | | 1.78E-07 | |
| GLN3 | -8.637705648 | | 1.78E-07 | |
| HAP43 | -8.637705648 | | 1.78E-07 | |
| HEM3 | -8.637705648 | | 1.78E-07 | |
| HGH1 | -8.637705648 | | 1.78E-07 | |
| HSP78 | -8.637705648 | | 1.78E-07 | |
| MIR1 | -8.637705648 | | 1.78E-07 | |
| MRPL19 | -8.637705648 | | 1.78E-07 | |
| NMD5 | -8.637705648 | | 1.78E-07 | |
| orf19.1360 | -8.637705648 | | 1.78E-07 | |
| orf19.1525 | -8.637705648 | | 1.78E-07 | |
| orf19.1849 | -8.637705648 | | 1.78E-07 | |
| orf19.2378 | -8.637705648 | | 1.78E-07 | |
| orf19.2519 | -8.637705648 | | 1.78E-07 | |
| orf19.2686 | -8.637705648 | | 1.78E-07 | |
| orf19.2853 | -8.637705648 | | 1.78E-07 | |
| orf19.3226 | -8.637705648 | | 1.78E-07 | |
| orf19.3353 | -8.637705648 | | 1.78E-07 | |
| orf19.3556 | -8.637705648 | | 1.78E-07 | |
| orf19.3571 | -8.637705648 | | 1.78E-07 | |
| orf19.3606 | -8.637705648 | | 1.78E-07 | |
| orf19.3689 | -8.637705648 | | 1.78E-07 | |
| orf19.4007 | -8.637705648 | | 1.78E-07 | |
| orf19.4080 | -8.637705648 | | 1.78E-07 | |
| orf19.4169 | -8.637705648 | | 1.78E-07 | |
| orf19.4185 | -8.637705648 | | 1.78E-07 | |
| orf19.4250 | -8.637705648 | | 1.78E-07 | |
| orf19.4316 | -8.637705648 | | 1.78E-07 | |
| orf19.4355 | -8.637705648 | | 1.78E-07 | |
| orf19.4787 | -8.637705648 | | 1.78E-07 | |
| orf19.4824 | -8.637705648 | | 1.78E-07 | |
| orf19.5049 | -8.637705648 | | 1.78E-07 | |
| orf19.5085 | -8.637705648 | | 1.78E-07 | |
| orf19.5090 | -8.637705648 | | 1.78E-07 | |
| orf19.515 | -8.637705648 | | 1.78E-07 | |
| orf19.5296 | -8.637705648 | | 1.78E-07 | |
| orf19.5376 | -8.637705648 | | 1.78E-07 | |
| orf19.5684.1 | -8.637705648 | | 1.78E-07 | |
| orf19.5688 | -8.637705648 | | 1.78E-07 | |
| orf19.5752 | -8.637705648 | | 1.78E-07 | |
| orf19.5929 | -8.637705648 | | 1.78E-07 | |
| orf19.6205 | -8.637705648 | | 1.78E-07 | |
| orf19.6528 | -8.637705648 | | 1.78E-07 | |
| orf19.6597 | -8.637705648 | | 1.78E-07 | |
| orf19.6989 | -8.637705648 | | 1.78E-07 | |
| orf19.7102 | -8.637705648 | | 1.78E-07 | |
| orf19.787.1 | -8.637705648 | | 1.78E-07 | |
| PDC2 | -8.637705648 | | 1.78E-07 | |
| PGA53 | -8.637705648 | | 1.78E-07 | |
| PZF1 | -8.637705648 | | 1.78E-07 | |
| RAD9 | -8.637705648 | | 1.78E-07 | |
| RMT2 | -8.637705648 | | 1.78E-07 | |
| RPN5 | -8.637705648 | | 1.78E-07 | |
| RSN1 | -8.637705648 | | 1.78E-07 | |
| RVS161 | -8.637705648 | | 1.78E-07 | |
| SLC1 | -8.637705648 | | 1.78E-07 | |
| SLD5 | -8.637705648 | | 1.78E-07 | |
| SPC34 | -8.637705648 | | 1.78E-07 | |
| SPT20 | -8.637705648 | | 1.78E-07 | |
| TLO11 | -8.637705648 | | 1.78E-07 | |
| TOM40 | -8.637705648 | | 1.78E-07 | |
| TOP2 | -8.637705648 | | 1.78E-07 | |
| TPO4 | -8.637705648 | | 1.78E-07 | |
| TYE7 | -8.637705648 | | 1.78E-07 | |
| URA1 | -8.637705648 | | 1.78E-07 | |
| VPS70 | -8.637705648 | | 1.78E-07 | |
| YHM1 | -8.637705648 | | 1.78E-07 | |
| YTH1 | -8.637705648 | | 1.78E-07 | |

| **Supplementary table 3. List of proteins that significantly changed their abundance pvalue <0.05 between PUJ256_FLU and PUJ256** | | |
| --- | --- | --- |
| **Protein** | **log2FC** | **p.value** |
| ADP1 | 5.43819897 | 4.98022E-05 |
| ATP17 | 5.43819897 | 4.98022E-05 |
| BMT4 | 5.43819897 | 4.98022E-05 |
| BUD31 | 5.43819897 | 4.98022E-05 |
| CCC1 | 5.43819897 | 4.98022E-05 |
| CHS2 | 5.43819897 | 4.98022E-05 |
| CHS8 | 5.43819897 | 4.98022E-05 |
| CSR1 | 5.43819897 | 4.98022E-05 |
| CTM1 | 5.43819897 | 4.98022E-05 |
| DAD4 | 5.43819897 | 4.98022E-05 |
| DAG7 | 5.43819897 | 4.98022E-05 |
| DBP7 | 5.43819897 | 4.98022E-05 |
| DUG3 | 5.43819897 | 4.98022E-05 |
| HOS3 | 5.43819897 | 4.98022E-05 |
| HYS2 | 5.43819897 | 4.98022E-05 |
| IRS4 | 5.43819897 | 4.98022E-05 |
| ITR1 | 5.43819897 | 4.98022E-05 |
| JEM1 | 5.43819897 | 4.98022E-05 |
| MIT1 | 5.43819897 | 4.98022E-05 |
| MNL1 | 5.43819897 | 4.98022E-05 |
| orf19.100 | 5.43819897 | 4.98022E-05 |
| orf19.1297 | 5.43819897 | 4.98022E-05 |
| orf19.1525 | 5.43819897 | 4.98022E-05 |
| orf19.1562 | 5.43819897 | 4.98022E-05 |
| orf19.1668 | 5.43819897 | 4.98022E-05 |
| orf19.1675 | 5.43819897 | 4.98022E-05 |
| orf19.1840 | 5.43819897 | 4.98022E-05 |
| orf19.1975 | 5.43819897 | 4.98022E-05 |
| orf19.2663 | 5.43819897 | 4.98022E-05 |
| orf19.2836 | 5.43819897 | 4.98022E-05 |
| orf19.3226 | 5.43819897 | 4.98022E-05 |
| orf19.3418 | 5.43819897 | 4.98022E-05 |
| orf19.354 | 5.43819897 | 4.98022E-05 |
| orf19.3606 | 5.43819897 | 4.98022E-05 |
| orf19.3874 | 5.43819897 | 4.98022E-05 |
| orf19.4169 | 5.43819897 | 4.98022E-05 |
| orf19.4390 | 5.43819897 | 4.98022E-05 |
| orf19.4589 | 5.43819897 | 4.98022E-05 |
| orf19.4724 | 5.43819897 | 4.98022E-05 |
| orf19.4824 | 5.43819897 | 4.98022E-05 |
| orf19.4893 | 5.43819897 | 4.98022E-05 |
| orf19.515 | 5.43819897 | 4.98022E-05 |
| orf19.5160 | 5.43819897 | 4.98022E-05 |
| orf19.5247 | 5.43819897 | 4.98022E-05 |
| orf19.5376 | 5.43819897 | 4.98022E-05 |
| orf19.580 | 5.43819897 | 4.98022E-05 |
| orf19.6023 | 5.43819897 | 4.98022E-05 |
| orf19.6185 | 5.43819897 | 4.98022E-05 |
| orf19.6342 | 5.43819897 | 4.98022E-05 |
| orf19.6396 | 5.43819897 | 4.98022E-05 |
| orf19.6487 | 5.43819897 | 4.98022E-05 |
| orf19.7316 | 5.43819897 | 4.98022E-05 |
| PHO100 | 5.43819897 | 4.98022E-05 |
| PHO91 | 5.43819897 | 4.98022E-05 |
| PIKA | 5.43819897 | 4.98022E-05 |
| PZF1 | 5.43819897 | 4.98022E-05 |
| RBE1 | 5.43819897 | 4.98022E-05 |
| SCH9 | 5.43819897 | 4.98022E-05 |
| SCP1 | 5.43819897 | 4.98022E-05 |
| SEN1 | 5.43819897 | 4.98022E-05 |
| SMI1B | 5.43819897 | 4.98022E-05 |
| SPT20 | 5.43819897 | 4.98022E-05 |
| STP4 | 5.43819897 | 4.98022E-05 |
| THG1 | 5.43819897 | 4.98022E-05 |
| TRP4 | 5.43819897 | 4.98022E-05 |
| TRS33 | 5.43819897 | 4.98022E-05 |
| WSC4 | 5.43819897 | 4.98022E-05 |
| YME1 | 5.43819897 | 4.98022E-05 |
| orf19.4068 | 3.33478797 | 0.04201614 |
| orf19.1285 | 2.70489688 | 0.005924604 |
| SIT4 | 2.46404898 | 0.03124203 |
| UBA4 | 2.43151832 | 0.034168936 |
| orf19.5003 | 2.28049276 | 0.013608845 |
| orf19.4423 | 2.2447504 | 0.044510926 |
| orf19.6227 | 2.09912443 | 0.007701858 |
| DDR48 | 2.09157657 | 0.006349444 |
| orf19.6503 | 2.05282316 | 0.049426416 |
| XOG1 | 1.95715942 | 0.004980216 |
| HNT1 | 1.91839996 | 0.045551863 |
| orf19.7589 | 1.85218791 | 0.025262384 |
| orf19.2988 | 1.83697326 | 0.042159751 |
| WOR3 | 1.53874632 | 0.013900562 |
| SCW11 | 1.39647886 | 0.034336892 |
| orf19.5295 | 1.32258479 | 0.010913964 |
| orf19.5607 | 1.16131241 | 0.02836351 |
| orf19.6268 | 1.13912515 | 0.043682127 |
| orf19.763 | -1.4192631 | 0.024044977 |
| ARO3 | -1.7234348 | 0.03549333 |
| orf19.5541 | -1.7371426 | 0.028044335 |
| GDS1 | -1.7825408 | 0.027466432 |
| LEU4 | -1.787546 | 0.022540112 |
| orf19.6392 | -1.8378476 | 0.046049644 |
| orf19.6360 | -1.9574721 | 0.009134074 |
| MED20 | -2.1111076 | 0.046462282 |
| BMT1 | -2.1801237 | 0.007100677 |
| FAS1 | -2.2031253 | 0.027429996 |
| HMS1 | -2.2250503 | 0.005835256 |
| orf19.1447 | -2.6788474 | 0.020889826 |
| orf19.3129 | -2.6890165 | 0.033105136 |
| orf19.5077 | -2.7652473 | 0.024012126 |
| BMS1 | -2.7891472 | 0.033072807 |
| orf19.7321 | -3.0234312 | 0.011772596 |
| AAH1 | -5.2830738 | 4.98022E-05 |
| ACC1 | -5.2830738 | 4.98022E-05 |
| ALD6 | -5.2830738 | 4.98022E-05 |
| ANP1 | -5.2830738 | 4.98022E-05 |
| ARC19 | -5.2830738 | 4.98022E-05 |
| ARF3 | -5.2830738 | 4.98022E-05 |
| ATO2 | -5.2830738 | 4.98022E-05 |
| COX2 | -5.2830738 | 4.98022E-05 |
| CTA4 | -5.2830738 | 4.98022E-05 |
| CTN3 | -5.2830738 | 4.98022E-05 |
| FLC2 | -5.2830738 | 4.98022E-05 |
| FMP45 | -5.2830738 | 4.98022E-05 |
| GAP1 | -5.2830738 | 4.98022E-05 |
| GPX3 | -5.2830738 | 4.98022E-05 |
| GTT12 | -5.2830738 | 4.98022E-05 |
| GYP1 | -5.2830738 | 4.98022E-05 |
| HAP43 | -5.2830738 | 4.98022E-05 |
| HYR1 | -5.2830738 | 4.98022E-05 |
| MEP1 | -5.2830738 | 4.98022E-05 |
| NAG6 | -5.2830738 | 4.98022E-05 |
| orf19.1034 | -5.2830738 | 4.98022E-05 |
| orf19.1054 | -5.2830738 | 4.98022E-05 |
| orf19.1113 | -5.2830738 | 4.98022E-05 |
| orf19.1114 | -5.2830738 | 4.98022E-05 |
| orf19.114 | -5.2830738 | 4.98022E-05 |
| orf19.1179 | -5.2830738 | 4.98022E-05 |
| orf19.1306 | -5.2830738 | 4.98022E-05 |
| orf19.1417 | -5.2830738 | 4.98022E-05 |
| orf19.1480 | -5.2830738 | 4.98022E-05 |
| orf19.1519 | -5.2830738 | 4.98022E-05 |
| orf19.1618.1 | -5.2830738 | 4.98022E-05 |
| orf19.1619 | -5.2830738 | 4.98022E-05 |
| orf19.1625 | -5.2830738 | 4.98022E-05 |
| orf19.1682 | -5.2830738 | 4.98022E-05 |
| orf19.1761 | -5.2830738 | 4.98022E-05 |
| orf19.1782 | -5.2830738 | 4.98022E-05 |
| orf19.1849 | -5.2830738 | 4.98022E-05 |
| orf19.1873 | -5.2830738 | 4.98022E-05 |
| orf19.216.1 | -5.2830738 | 4.98022E-05 |
| orf19.2168.3 | -5.2830738 | 4.98022E-05 |
| orf19.2208 | -5.2830738 | 4.98022E-05 |
| orf19.2228 | -5.2830738 | 4.98022E-05 |
| orf19.2318.1 | -5.2830738 | 4.98022E-05 |
| orf19.2472.1 | -5.2830738 | 4.98022E-05 |
| orf19.2673 | -5.2830738 | 4.98022E-05 |
| orf19.2848 | -5.2830738 | 4.98022E-05 |
| orf19.2863 | -5.2830738 | 4.98022E-05 |
| orf19.2954 | -5.2830738 | 4.98022E-05 |
| orf19.3057 | -5.2830738 | 4.98022E-05 |
| orf19.3087.1 | -5.2830738 | 4.98022E-05 |
| orf19.3128 | -5.2830738 | 4.98022E-05 |
| orf19.3250 | -5.2830738 | 4.98022E-05 |
| orf19.3272 | -5.2830738 | 4.98022E-05 |
| orf19.3366.1 | -5.2830738 | 4.98022E-05 |
| orf19.3406 | -5.2830738 | 4.98022E-05 |
| orf19.3483 | -5.2830738 | 4.98022E-05 |
| orf19.3552 | -5.2830738 | 4.98022E-05 |
| orf19.3659 | -5.2830738 | 4.98022E-05 |
| orf19.3684 | -5.2830738 | 4.98022E-05 |
| orf19.3782.2 | -5.2830738 | 4.98022E-05 |
| orf19.3843 | -5.2830738 | 4.98022E-05 |
| orf19.3965 | -5.2830738 | 4.98022E-05 |
| orf19.3970 | -5.2830738 | 4.98022E-05 |
| orf19.409 | -5.2830738 | 4.98022E-05 |
| orf19.4132 | -5.2830738 | 4.98022E-05 |
| orf19.4163 | -5.2830738 | 4.98022E-05 |
| orf19.4355 | -5.2830738 | 4.98022E-05 |
| orf19.4358 | -5.2830738 | 4.98022E-05 |
| orf19.4365 | -5.2830738 | 4.98022E-05 |
| orf19.446.2 | -5.2830738 | 4.98022E-05 |
| orf19.4522 | -5.2830738 | 4.98022E-05 |
| orf19.4563 | -5.2830738 | 4.98022E-05 |
| orf19.4643 | -5.2830738 | 4.98022E-05 |
| orf19.4676 | -5.2830738 | 4.98022E-05 |
| orf19.4792 | -5.2830738 | 4.98022E-05 |
| orf19.4886 | -5.2830738 | 4.98022E-05 |
| orf19.491 | -5.2830738 | 4.98022E-05 |
| orf19.4947 | -5.2830738 | 4.98022E-05 |
| orf19.5043 | -5.2830738 | 4.98022E-05 |
| orf19.5079.1 | -5.2830738 | 4.98022E-05 |
| orf19.5085 | -5.2830738 | 4.98022E-05 |
| orf19.5238 | -5.2830738 | 4.98022E-05 |
| orf19.5300 | -5.2830738 | 4.98022E-05 |
| orf19.5411 | -5.2830738 | 4.98022E-05 |
| orf19.5418 | -5.2830738 | 4.98022E-05 |
| orf19.556 | -5.2830738 | 4.98022E-05 |
| orf19.5660.1 | -5.2830738 | 4.98022E-05 |
| orf19.5727 | -5.2830738 | 4.98022E-05 |
| orf19.6035 | -5.2830738 | 4.98022E-05 |
| orf19.6039 | -5.2830738 | 4.98022E-05 |
| orf19.6062 | -5.2830738 | 4.98022E-05 |
| orf19.6189 | -5.2830738 | 4.98022E-05 |
| orf19.6198.1 | -5.2830738 | 4.98022E-05 |
| orf19.6247.1 | -5.2830738 | 4.98022E-05 |
| orf19.642 | -5.2830738 | 4.98022E-05 |
| orf19.643 | -5.2830738 | 4.98022E-05 |
| orf19.6474 | -5.2830738 | 4.98022E-05 |
| orf19.6565 | -5.2830738 | 4.98022E-05 |
| orf19.6607 | -5.2830738 | 4.98022E-05 |
| orf19.6639 | -5.2830738 | 4.98022E-05 |
| orf19.6769 | -5.2830738 | 4.98022E-05 |
| orf19.6923 | -5.2830738 | 4.98022E-05 |
| orf19.6980 | -5.2830738 | 4.98022E-05 |
| orf19.7078 | -5.2830738 | 4.98022E-05 |
| orf19.7088 | -5.2830738 | 4.98022E-05 |
| orf19.7111 | -5.2830738 | 4.98022E-05 |
| orf19.7183 | -5.2830738 | 4.98022E-05 |
| orf19.7202 | -5.2830738 | 4.98022E-05 |
| orf19.7326 | -5.2830738 | 4.98022E-05 |
| orf19.7344 | -5.2830738 | 4.98022E-05 |
| orf19.752 | -5.2830738 | 4.98022E-05 |
| orf19.7566 | -5.2830738 | 4.98022E-05 |
| orf19.757 | -5.2830738 | 4.98022E-05 |
| orf19.7604 | -5.2830738 | 4.98022E-05 |
| orf19.787.1 | -5.2830738 | 4.98022E-05 |
| orf19.813 | -5.2830738 | 4.98022E-05 |
| orf19.831 | -5.2830738 | 4.98022E-05 |
| PAM16 | -5.2830738 | 4.98022E-05 |
| PAM18 | -5.2830738 | 4.98022E-05 |
| PEX11 | -5.2830738 | 4.98022E-05 |
| PHO8 | -5.2830738 | 4.98022E-05 |
| PHO88 | -5.2830738 | 4.98022E-05 |
| PMT1 | -5.2830738 | 4.98022E-05 |
| PRP5 | -5.2830738 | 4.98022E-05 |
| QCR8 | -5.2830738 | 4.98022E-05 |
| RAC1 | -5.2830738 | 4.98022E-05 |
| RAX1 | -5.2830738 | 4.98022E-05 |
| RFC2 | -5.2830738 | 4.98022E-05 |
| RHO2 | -5.2830738 | 4.98022E-05 |
| RIA1 | -5.2830738 | 4.98022E-05 |
| RIM2 | -5.2830738 | 4.98022E-05 |
| RMT2 | -5.2830738 | 4.98022E-05 |
| RPC10 | -5.2830738 | 4.98022E-05 |
| RVS161 | -5.2830738 | 4.98022E-05 |
| SAM51 | -5.2830738 | 4.98022E-05 |
| SAP5 | -5.2830738 | 4.98022E-05 |
| SCO1 | -5.2830738 | 4.98022E-05 |
| SDH1 | -5.2830738 | 4.98022E-05 |
| SEN15 | -5.2830738 | 4.98022E-05 |
| SFT1 | -5.2830738 | 4.98022E-05 |
| SLN1 | -5.2830738 | 4.98022E-05 |
| SMF12 | -5.2830738 | 4.98022E-05 |
| SPC2 | -5.2830738 | 4.98022E-05 |
| SPP1 | -5.2830738 | 4.98022E-05 |
| SUR2 | -5.2830738 | 4.98022E-05 |
| TIM22 | -5.2830738 | 4.98022E-05 |
| TOM40 | -5.2830738 | 4.98022E-05 |
| TOM7 | -5.2830738 | 4.98022E-05 |
| URA1 | -5.2830738 | 4.98022E-05 |
| VPS70 | -5.2830738 | 4.98022E-05 |
| VTC4 | -5.2830738 | 4.98022E-05 |
| YHM1 | -5.2830738 | 4.98022E-05 |
| YIM1 | -5.2830738 | 4.98022E-05 |
| YPT72 | -5.2830738 | 4.98022E-05 |
| ZCF15 | -5.2830738 | 4.98022E-05 |

| **Supplementary table 4. List of proteins that significantly changed their abundance pvalue <0.05 between PUJ256_FLU and SC5314_FLU** | | |
| --- | --- | --- |
| **Protein** | **log2FC** | **p.value** |
| ACC1 | -7.338553484 | 1.00847E-07 |
| AGP2 | -7.338553484 | 1.00847E-07 |
| ALD6 | -7.338553484 | 1.00847E-07 |
| ATO2 | -7.338553484 | 1.00847E-07 |
| BUR2 | -7.338553484 | 1.00847E-07 |
| CCR4 | -7.338553484 | 1.00847E-07 |
| CDC42 | -7.338553484 | 1.00847E-07 |
| CMK1 | -7.338553484 | 1.00847E-07 |
| CTA4 | -7.338553484 | 1.00847E-07 |
| CTN3 | -7.338553484 | 1.00847E-07 |
| DIP5 | -7.338553484 | 1.00847E-07 |
| DNM1 | -7.338553484 | 1.00847E-07 |
| DUN1 | -7.338553484 | 1.00847E-07 |
| ECM7 | -7.338553484 | 1.00847E-07 |
| ENA2 | -7.338553484 | 1.00847E-07 |
| EXO84 | -7.338553484 | 1.00847E-07 |
| FCR1 | -7.338553484 | 1.00847E-07 |
| FLC2 | -7.338553484 | 1.00847E-07 |
| FMP45 | -7.338553484 | 1.00847E-07 |
| GAP1 | -7.338553484 | 1.00847E-07 |
| GPX2 | -7.338553484 | 1.00847E-07 |
| GTT12 | -7.338553484 | 1.00847E-07 |
| GUT1 | -7.338553484 | 1.00847E-07 |
| GYP1 | -7.338553484 | 1.00847E-07 |
| GYP2 | -7.338553484 | 1.00847E-07 |
| GYP7 | -7.338553484 | 1.00847E-07 |
| HDA1 | -7.338553484 | 1.00847E-07 |
| HGT2 | -7.338553484 | 1.00847E-07 |
| HYR1 | -7.338553484 | 1.00847E-07 |
| IFR1 | -7.338553484 | 1.00847E-07 |
| IHD1 | -7.338553484 | 1.00847E-07 |
| MEP1 | -7.338553484 | 1.00847E-07 |
| MRV2 | -7.338553484 | 1.00847E-07 |
| MUP1 | -7.338553484 | 1.00847E-07 |
| NOC2 | -7.338553484 | 1.00847E-07 |
| orf19.1117 | -7.338553484 | 1.00847E-07 |
| orf19.114 | -7.338553484 | 1.00847E-07 |
| orf19.1179 | -7.338553484 | 1.00847E-07 |
| orf19.1306 | -7.338553484 | 1.00847E-07 |
| orf19.1519 | -7.338553484 | 1.00847E-07 |
| orf19.1619 | -7.338553484 | 1.00847E-07 |
| orf19.1764 | -7.338553484 | 1.00847E-07 |
| orf19.1782 | -7.338553484 | 1.00847E-07 |
| orf19.1800 | -7.338553484 | 1.00847E-07 |
| orf19.1871 | -7.338553484 | 1.00847E-07 |
| orf19.2017 | -7.338553484 | 1.00847E-07 |
| orf19.213 | -7.338553484 | 1.00847E-07 |
| orf19.2168.3 | -7.338553484 | 1.00847E-07 |
| orf19.2208 | -7.338553484 | 1.00847E-07 |
| orf19.2476 | -7.338553484 | 1.00847E-07 |
| orf19.2728 | -7.338553484 | 1.00847E-07 |
| orf19.2770 | -7.338553484 | 1.00847E-07 |
| orf19.2838 | -7.338553484 | 1.00847E-07 |
| orf19.2848 | -7.338553484 | 1.00847E-07 |
| orf19.3087.1 | -7.338553484 | 1.00847E-07 |
| orf19.3250 | -7.338553484 | 1.00847E-07 |
| orf19.3272 | -7.338553484 | 1.00847E-07 |
| orf19.3406 | -7.338553484 | 1.00847E-07 |
| orf19.3483 | -7.338553484 | 1.00847E-07 |
| orf19.3615 | -7.338553484 | 1.00847E-07 |
| orf19.3684 | -7.338553484 | 1.00847E-07 |
| orf19.3831 | -7.338553484 | 1.00847E-07 |
| orf19.3932.1 | -7.338553484 | 1.00847E-07 |
| orf19.3970 | -7.338553484 | 1.00847E-07 |
| orf19.4117 | -7.338553484 | 1.00847E-07 |
| orf19.4163 | -7.338553484 | 1.00847E-07 |
| orf19.4292 | -7.338553484 | 1.00847E-07 |
| orf19.4522 | -7.338553484 | 1.00847E-07 |
| orf19.4563 | -7.338553484 | 1.00847E-07 |
| orf19.4643 | -7.338553484 | 1.00847E-07 |
| orf19.4676 | -7.338553484 | 1.00847E-07 |
| orf19.4749 | -7.338553484 | 1.00847E-07 |
| orf19.4792 | -7.338553484 | 1.00847E-07 |
| orf19.4886 | -7.338553484 | 1.00847E-07 |
| orf19.491 | -7.338553484 | 1.00847E-07 |
| orf19.5043 | -7.338553484 | 1.00847E-07 |
| orf19.5092 | -7.338553484 | 1.00847E-07 |
| orf19.5269 | -7.338553484 | 1.00847E-07 |
| orf19.5300 | -7.338553484 | 1.00847E-07 |
| orf19.5418 | -7.338553484 | 1.00847E-07 |
| orf19.5555 | -7.338553484 | 1.00847E-07 |
| orf19.5660.1 | -7.338553484 | 1.00847E-07 |
| orf19.5727 | -7.338553484 | 1.00847E-07 |
| orf19.5814.1 | -7.338553484 | 1.00847E-07 |
| orf19.5852 | -7.338553484 | 1.00847E-07 |
| orf19.5905 | -7.338553484 | 1.00847E-07 |
| orf19.6233 | -7.338553484 | 1.00847E-07 |
| orf19.643 | -7.338553484 | 1.00847E-07 |
| orf19.6484 | -7.338553484 | 1.00847E-07 |
| orf19.6581 | -7.338553484 | 1.00847E-07 |
| orf19.6601 | -7.338553484 | 1.00847E-07 |
| orf19.6639 | -7.338553484 | 1.00847E-07 |
| orf19.6769 | -7.338553484 | 1.00847E-07 |
| orf19.6790 | -7.338553484 | 1.00847E-07 |
| orf19.6923 | -7.338553484 | 1.00847E-07 |
| orf19.6980 | -7.338553484 | 1.00847E-07 |
| orf19.7058 | -7.338553484 | 1.00847E-07 |
| orf19.7078 | -7.338553484 | 1.00847E-07 |
| orf19.7111 | -7.338553484 | 1.00847E-07 |
| orf19.7183 | -7.338553484 | 1.00847E-07 |
| orf19.7194 | -7.338553484 | 1.00847E-07 |
| orf19.7326 | -7.338553484 | 1.00847E-07 |
| orf19.7425 | -7.338553484 | 1.00847E-07 |
| orf19.752 | -7.338553484 | 1.00847E-07 |
| orf19.757 | -7.338553484 | 1.00847E-07 |
| orf19.813 | -7.338553484 | 1.00847E-07 |
| orf19.831 | -7.338553484 | 1.00847E-07 |
| PEX1 | -7.338553484 | 1.00847E-07 |
| PEX6 | -7.338553484 | 1.00847E-07 |
| PHO8 | -7.338553484 | 1.00847E-07 |
| PHO88 | -7.338553484 | 1.00847E-07 |
| PHO89 | -7.338553484 | 1.00847E-07 |
| PLB1 | -7.338553484 | 1.00847E-07 |
| PMR1 | -7.338553484 | 1.00847E-07 |
| PRP5 | -7.338553484 | 1.00847E-07 |
| PWP2 | -7.338553484 | 1.00847E-07 |
| RAC1 | -7.338553484 | 1.00847E-07 |
| RAX1 | -7.338553484 | 1.00847E-07 |
| RIA1 | -7.338553484 | 1.00847E-07 |
| RIM2 | -7.338553484 | 1.00847E-07 |
| RPC10 | -7.338553484 | 1.00847E-07 |
| SAP5 | -7.338553484 | 1.00847E-07 |
| SDH1 | -7.338553484 | 1.00847E-07 |
| SEN15 | -7.338553484 | 1.00847E-07 |
| SLU7 | -7.338553484 | 1.00847E-07 |
| SMF12 | -7.338553484 | 1.00847E-07 |
| SPP1 | -7.338553484 | 1.00847E-07 |
| SUR2 | -7.338553484 | 1.00847E-07 |
| TCC1 | -7.338553484 | 1.00847E-07 |
| TFC4 | -7.338553484 | 1.00847E-07 |
| TOM7 | -7.338553484 | 1.00847E-07 |
| TPS1 | -7.338553484 | 1.00847E-07 |
| VTC4 | -7.338553484 | 1.00847E-07 |
| YBL053 | -7.338553484 | 1.00847E-07 |
| ZCF15 | -7.338553484 | 1.00847E-07 |
| HGT6 | -5.338553484 | 0.000136064 |
| PGA31 | -4.75691702 | 0.004572615 |
| SOD3 | -4.730356209 | 0.000710793 |
| orf19.1840 | -4.502226864 | 0.001367401 |
| RIX7 | -4.318299103 | 0.018782178 |
| ENA21 | -4.041836977 | 0.000678464 |
| SPT10 | -3.927995692 | 0.00168035 |
| orf19.6599.1 | -3.9045404 | 0.013387253 |
| AFP99 | -3.823072699 | 0.000749982 |
| orf19.5620 | -3.720600708 | 0.000102245 |
| SEN1 | -3.653714377 | 0.01584735 |
| RHO1 | -3.608175081 | 0.002149016 |
| RHO3 | -3.566314814 | 0.01072571 |
| HMS1 | -3.391193181 | 0.000282165 |
| XOG1 | -3.353131244 | 3.60863E-05 |
| orf19.2125 | -3.241248379 | 4.43866E-05 |
| orf19.4258 | -3.209828385 | 0.012929647 |
| VTC3 | -3.196710828 | 0.032872882 |
| orf19.7504 | -3.195011854 | 0.000314811 |
| PMC1 | -3.102802332 | 0.000522281 |
| PHO84 | -3.087155777 | 0.009558187 |
| orf19.7006 | -3.03366373 | 0.000572039 |
| PHM7 | -3.02490521 | 0.000278648 |
| orf19.1447 | -3.023166913 | 0.005054059 |
| PMA1 | -2.975456241 | 0.001059435 |
| orf19.1414 | -2.956649697 | 0.003542064 |
| GCA2 | -2.926573637 | 0.005815241 |
| KEX2 | -2.88653802 | 2.39839E-05 |
| NRP1 | -2.818658864 | 0.012173306 |
| orf19.993 | -2.767932279 | 0.000938292 |
| HGT1 | -2.690946007 | 0.004960194 |
| orf19.7321 | -2.665913249 | 0.015357555 |
| RCT1 | -2.652921574 | 0.000628736 |
| SEC4 | -2.605851458 | 0.032487503 |
| orf19.6311 | -2.595376665 | 0.001666784 |
| orf19.3418 | -2.572611031 | 0.028634657 |
| RPT4 | -2.494392027 | 0.00509372 |
| orf19.7459 | -2.486188869 | 0.000310752 |
| orf19.2442 | -2.44199668 | 0.035664786 |
| orf19.2867 | -2.420485778 | 0.010329583 |
| KRE6 | -2.397390445 | 0.001633313 |
| orf19.7296 | -2.351813582 | 0.023118811 |
| SEC7 | -2.34916831 | 0.01376457 |
| orf19.6637 | -2.341169395 | 0.000217386 |
| SNX4 | -2.337779953 | 0.016512664 |
| BZZ1 | -2.305768822 | 0.022497588 |
| orf19.7316 | -2.300577043 | 0.034189874 |
| orf19.3213 | -2.262586294 | 0.042908151 |
| orf19.2917.1 | -2.238985021 | 0.04370091 |
| SCP1 | -2.220441295 | 0.023695266 |
| orf19.5295 | -2.129271662 | 0.00083156 |
| RBT4 | -2.123587944 | 0.002253288 |
| HRT2 | -2.099608611 | 0.01953674 |
| GCD2 | -2.076002938 | 0.020435044 |
| FAS2 | -2.067659734 | 0.012684618 |
| orf19.7109 | -2.050173447 | 0.023522496 |
| SGA1 | -2.037596845 | 0.000359851 |
| orf19.3463 | -2.023192402 | 0.014562157 |
| orf19.7085 | -2.018100305 | 0.034074636 |
| orf19.2769 | -2.006286399 | 0.008734785 |
| orf19.4068 | -1.998678206 | 0.024873143 |
| orf19.592 | -1.994956424 | 0.003236221 |
| SPT7 | -1.957586376 | 0.04534687 |
| DAP1 | -1.899402334 | 0.004120246 |
| ASR1 | -1.893862102 | 0.015303403 |
| BUD21 | -1.885483031 | 0.023707358 |
| orf19.3932 | -1.873880759 | 0.002304235 |
| LYS1 | -1.872218361 | 0.007838623 |
| PRX1 | -1.863184199 | 0.01508286 |
| SAP10 | -1.844290499 | 0.024921041 |
| DDR48 | -1.842548457 | 0.004663148 |
| BEM2 | -1.823318665 | 0.003255472 |
| orf19.6360 | -1.759656766 | 0.001984976 |
| SRO77 | -1.748687673 | 0.042390371 |
| IDH1 | -1.748379687 | 0.002062347 |
| PUF3 | -1.741328704 | 0.006945921 |
| orf19.1336.2 | -1.720136995 | 0.009361737 |
| orf19.6789 | -1.709885166 | 0.007997049 |
| COQ5 | -1.695025934 | 0.008304321 |
| ECM4 | -1.63957977 | 0.028717717 |
| orf19.4825 | -1.62629072 | 0.030659755 |
| RPS12 | -1.606901881 | 0.010197209 |
| ARG5,6 | -1.603781115 | 0.041659095 |
| COX19 | -1.598991257 | 0.028480577 |
| TUB1 | -1.583900945 | 0.009386912 |
| orf19.4043 | -1.572009926 | 0.020639086 |
| orf19.4184 | -1.561542249 | 0.006105252 |
| SPB1 | -1.538891669 | 0.017882778 |
| LYS22 | -1.517838005 | 0.020554298 |
| IST2 | -1.515343637 | 0.033961646 |
| HSP70 | -1.501041698 | 0.012661825 |
| orf19.2368 | -1.480929602 | 0.007591571 |
| orf19.5516 | -1.480021024 | 0.035563383 |
| SYN8 | -1.464419485 | 0.012540694 |
| RLP24 | -1.463563576 | 0.027557824 |
| SAP9 | -1.433023102 | 0.015115095 |
| orf19.3007 | -1.431426607 | 0.045190944 |
| IDH2 | -1.431316818 | 0.007436637 |
| PNG2 | -1.422151096 | 0.008972506 |
| PHR2 | -1.415801492 | 0.00262319 |
| orf19.273 | -1.373430983 | 0.022668683 |
| orf19.1122 | -1.373310647 | 0.041476686 |
| LAG1 | -1.364343928 | 0.046404899 |
| orf19.716 | -1.351358075 | 0.031421492 |
| orf19.6628 | -1.350926899 | 0.04672636 |
| ASR3 | -1.34196312 | 0.029971641 |
| ATP18 | -1.33896108 | 0.017269359 |
| COX13 | -1.330894194 | 0.041829628 |
| ATP1 | -1.322154365 | 0.036773835 |
| orf19.1376 | -1.300861279 | 0.016326544 |
| orf19.2296 | -1.297902111 | 0.006933247 |
| ERG6 | -1.287192481 | 0.046878555 |
| SOD5 | -1.245694995 | 0.032133195 |
| CRH11 | -1.241610284 | 0.039342327 |
| BUD14 | -1.209125386 | 0.049043541 |
| LSC2 | -1.190844653 | 0.028399923 |
| RNR22 | -1.168902134 | 0.037748122 |
| orf19.3310 | -1.123804908 | 0.037794186 |
| RBD1 | -1.117592164 | 0.045171383 |
| orf19.3335 | -1.116968331 | 0.034098281 |
| COI1 | -1.107205202 | 0.03654961 |
| orf19.1609 | -1.094903825 | 0.021943977 |
| CHS3 | -1.051611699 | 0.031406782 |
| orf19.4839 | -1.051408298 | 0.047986263 |
| orf19.2008 | -1.047297047 | 0.03489816 |
| ATP16 | -1.007431008 | 0.028255382 |
| CNB1 | 1.021698031 | 0.017797129 |
| FCR3 | 1.042612611 | 0.049964026 |
| orf19.4639 | 1.052702153 | 0.042366534 |
| orf19.5394.1 | 1.056420072 | 0.037494141 |
| CDC37 | 1.066639987 | 0.019747102 |
| orf19.1421 | 1.080531438 | 0.036204906 |
| RPL29 | 1.088186879 | 0.034065213 |
| orf19.5833 | 1.090379344 | 0.026034288 |
| IPP1 | 1.175648882 | 0.023712654 |
| ARO4 | 1.179625427 | 0.025048303 |
| RPS17B | 1.179933582 | 0.039650441 |
| NMA111 | 1.191578986 | 0.042407641 |
| RPL5 | 1.225832917 | 0.02081803 |
| TKL1 | 1.227440104 | 0.014150139 |
| HMO1 | 1.238002784 | 0.026977166 |
| orf19.4888 | 1.274550539 | 0.021311887 |
| IMH3 | 1.282843579 | 0.022822963 |
| PGK1 | 1.296834711 | 0.013752244 |
| SPE2 | 1.29773748 | 0.018360454 |
| CAR1 | 1.3124461 | 0.017530168 |
| orf19.6604 | 1.312697641 | 0.025038147 |
| GRS1 | 1.321577763 | 0.006173788 |
| IDP1 | 1.326809971 | 0.011093693 |
| orf19.1697 | 1.33249865 | 0.012995039 |
| CMP1 | 1.336353201 | 0.025615249 |
| orf19.7199 | 1.338449795 | 0.005790345 |
| orf19.5158 | 1.346042501 | 0.010402826 |
| orf19.6596 | 1.36193812 | 0.018716369 |
| orf19.1626 | 1.366831563 | 0.015684096 |
| orf19.1085 | 1.373629373 | 0.010628269 |
| orf19.4078 | 1.375381005 | 0.029387085 |
| orf19.805 | 1.400708429 | 0.010486332 |
| orf19.4220 | 1.418552162 | 0.047454685 |
| CGR1 | 1.446412433 | 0.013942161 |
| orf19.3140.1 | 1.449315765 | 0.017777541 |
| TRY2 | 1.461407939 | 0.022291566 |
| orf19.7329 | 1.463067214 | 0.009929056 |
| GIG1 | 1.470934347 | 0.011505171 |
| RPL12 | 1.497246924 | 0.038197822 |
| orf19.3635 | 1.503201863 | 0.049591437 |
| RPL23A | 1.506617729 | 0.010083303 |
| orf19.5747 | 1.514570453 | 0.023250954 |
| SEC23 | 1.534544541 | 0.008483763 |
| ARD | 1.548442624 | 0.021345289 |
| GIS2 | 1.550517346 | 0.033271546 |
| orf19.3649 | 1.580871557 | 0.001597234 |
| orf19.1180 | 1.589470349 | 0.003541742 |
| ATC1 | 1.593129506 | 0.027110239 |
| orf19.7297 | 1.595535047 | 0.031146199 |
| orf19.2335 | 1.612115752 | 0.044896738 |
| orf19.3312 | 1.62530628 | 0.03343209 |
| FBA1 | 1.629105146 | 0.015952243 |
| orf19.6230 | 1.644568372 | 0.023980584 |
| orf19.1267.1 | 1.646352842 | 0.009319458 |
| GCD11 | 1.664132153 | 0.011978161 |
| SSZ1 | 1.665910298 | 0.007753219 |
| UGP1 | 1.699938864 | 0.021690098 |
| MED3 | 1.700014324 | 0.02670224 |
| orf19.6917 | 1.707919447 | 0.017703167 |
| orf19.1409.1 | 1.723193036 | 0.026734644 |
| orf19.81 | 1.738178014 | 0.029928392 |
| PBP2 | 1.746560455 | 0.038640631 |
| orf19.3342 | 1.747723947 | 0.040927001 |
| orf19.4532 | 1.752020014 | 0.040113288 |
| ARF2 | 1.77969263 | 0.030584158 |
| orf19.3235 | 1.795777192 | 0.006507122 |
| RPL6 | 1.800074842 | 0.018990351 |
| MDH1 | 1.813488963 | 0.006304205 |
| SUI2 | 1.835273209 | 0.004309655 |
| RPL38 | 1.8459295 | 0.039193018 |
| orf19.967 | 1.855017065 | 0.04734972 |
| BUD23 | 1.889271144 | 0.004304732 |
| orf19.2607 | 1.901934479 | 0.036584103 |
| orf19.6539 | 1.910727312 | 0.011991118 |
| GRX3 | 1.920891165 | 0.000699268 |
| BOI2 | 1.925990229 | 0.016051671 |
| WRS1 | 1.929641733 | 0.002904566 |
| MTR2 | 1.932783398 | 0.004585434 |
| UGA11 | 1.948469561 | 0.012069368 |
| ADE5,7 | 1.949712572 | 0.010389833 |
| orf19.5607 | 1.952269738 | 0.001797305 |
| orf19.3156 | 1.953650229 | 0.011727745 |
| LEU1 | 1.97115599 | 0.017613707 |
| XKS1 | 1.983042875 | 0.001243873 |
| RPF2 | 2.006791373 | 0.016585355 |
| RSM22 | 2.015165177 | 0.005591632 |
| EBP1 | 2.026842129 | 0.003383356 |
| PHA2 | 2.058482662 | 0.026815413 |
| RPL11 | 2.061846774 | 0.017677468 |
| orf19.1545 | 2.066098353 | 0.041103417 |
| MED9 | 2.068382846 | 0.005808011 |
| orf19.6392 | 2.094845239 | 0.049987329 |
| orf19.1796 | 2.116701248 | 0.002570773 |
| PTC7 | 2.128994986 | 0.019888437 |
| ELC1 | 2.131596397 | 0.000832202 |
| orf19.1026 | 2.14224705 | 0.00190278 |
| orf19.376 | 2.148763169 | 0.009066231 |
| UBC15 | 2.149420522 | 0.002436677 |
| GUK1 | 2.163859845 | 0.014066054 |
| PRP3 | 2.16988435 | 0.020607855 |
| orf19.86 | 2.191188808 | 0.000118853 |
| orf19.5987 | 2.199450018 | 0.020806427 |
| orf19.6509 | 2.20524052 | 0.037106217 |
| MDH1-3 | 2.207269314 | 0.002596351 |
| MCM6 | 2.212614776 | 0.033646827 |
| orf19.6227 | 2.229475726 | 0.02510734 |
| BRE1 | 2.239642021 | 0.002718902 |
| orf19.7531 | 2.254552864 | 0.001057357 |
| orf19.6355 | 2.255139025 | 0.004658187 |
| GAL10 | 2.263576668 | 0.025959748 |
| orf19.4167 | 2.273944213 | 0.01456424 |
| SPL1 | 2.301616824 | 0.002105135 |
| FRS1 | 2.316489241 | 0.043625626 |
| OYE23 | 2.359345889 | 0.007557247 |
| orf19.2965 | 2.381041967 | 0.024465764 |
| RFA1 | 2.399590431 | 0.017381693 |
| PRT1 | 2.425529625 | 0.023888972 |
| orf19.5682 | 2.439272557 | 0.009425656 |
| orf19.6883 | 2.452147374 | 0.011729655 |
| orf19.4904 | 2.474903049 | 0.010803085 |
| IQG1 | 2.478889893 | 0.02837127 |
| orf19.2671 | 2.49118672 | 0.023758246 |
| ECM38 | 2.518835198 | 0.00151672 |
| orf19.185 | 2.527467411 | 0.017391547 |
| orf19.6739 | 2.532407875 | 0.014696461 |
| RPL17B | 2.538205816 | 0.037289486 |
| orf19.1460 | 2.567123071 | 0.000118828 |
| orf19.500 | 2.57499801 | 0.007645623 |
| orf19.6054 | 2.586346475 | 0.013848011 |
| orf19.4953 | 2.609457638 | 0.002522099 |
| orf19.7263 | 2.66237194 | 0.000707423 |
| KRE5 | 2.713075234 | 0.002198207 |
| PTC5 | 2.724447881 | 0.004522132 |
| MRT4 | 2.736936276 | 0.002751174 |
| orf19.6056 | 2.768157505 | 0.002453041 |
| MRP17 | 2.800686489 | 0.021948186 |
| IPK2 | 2.8359567 | 0.014044036 |
| RHR2 | 2.846905089 | 0.000759803 |
| orf19.3482 | 2.87682015 | 0.00035301 |
| orf19.7234 | 2.877884909 | 0.017926479 |
| MUQ1 | 2.901051884 | 0.008520006 |
| TAF145 | 3.01992191 | 0.009460632 |
| RPL30 | 3.11154893 | 0.014481719 |
| orf19.1355 | 3.900638177 | 0.000757419 |
| RLI1 | 3.924301188 | 0.003284677 |
| orf19.5783 | 4.188924187 | 0.001609213 |
| orf19.577 | 4.190008084 | 0.045284837 |
| MEX67 | 4.243874504 | 0.000231922 |
| AGO1 | 4.665168301 | 0.001714264 |
| RAT1 | 4.736997658 | 0.021434389 |
| orf19.7345 | 5.088532435 | 0.000433285 |
| orf19.6306 | 5.106989955 | 1.00847E-05 |
| NUP | 5.405937191 | 3.69547E-05 |
| orf19.4474 | 6.230224349 | 0.013694496 |
| ABP140 | 8.230224349 | 1.00847E-07 |
| ARC35 | 8.230224349 | 1.00847E-07 |
| ATP17 | 8.230224349 | 1.00847E-07 |
| BMT1 | 8.230224349 | 1.00847E-07 |
| BNA4 | 8.230224349 | 1.00847E-07 |
| CCT7 | 8.230224349 | 1.00847E-07 |
| CEF1 | 8.230224349 | 1.00847E-07 |
| CHS8 | 8.230224349 | 1.00847E-07 |
| COX1 | 8.230224349 | 1.00847E-07 |
| CSR1 | 8.230224349 | 1.00847E-07 |
| CTA24 | 8.230224349 | 1.00847E-07 |
| CTA8 | 8.230224349 | 1.00847E-07 |
| CTF5 | 8.230224349 | 1.00847E-07 |
| CTF8 | 8.230224349 | 1.00847E-07 |
| CTM1 | 8.230224349 | 1.00847E-07 |
| DAO1 | 8.230224349 | 1.00847E-07 |
| DBP7 | 8.230224349 | 1.00847E-07 |
| DEF1 | 8.230224349 | 1.00847E-07 |
| DIP2 | 8.230224349 | 1.00847E-07 |
| DRG1 | 8.230224349 | 1.00847E-07 |
| DUG3 | 8.230224349 | 1.00847E-07 |
| ERG2 | 8.230224349 | 1.00847E-07 |
| GLN3 | 8.230224349 | 1.00847E-07 |
| HEM3 | 8.230224349 | 1.00847E-07 |
| HGH1 | 8.230224349 | 1.00847E-07 |
| HOS3 | 8.230224349 | 1.00847E-07 |
| HPA2 | 8.230224349 | 1.00847E-07 |
| HSP78 | 8.230224349 | 1.00847E-07 |
| MED20 | 8.230224349 | 1.00847E-07 |
| MET3 | 8.230224349 | 1.00847E-07 |
| OCA1 | 8.230224349 | 1.00847E-07 |
| orf19.102 | 8.230224349 | 1.00847E-07 |
| orf19.1360 | 8.230224349 | 1.00847E-07 |
| orf19.1525 | 8.230224349 | 1.00847E-07 |
| orf19.1562 | 8.230224349 | 1.00847E-07 |
| orf19.1940 | 8.230224349 | 1.00847E-07 |
| orf19.2378 | 8.230224349 | 1.00847E-07 |
| orf19.2400 | 8.230224349 | 1.00847E-07 |
| orf19.2519 | 8.230224349 | 1.00847E-07 |
| orf19.2686 | 8.230224349 | 1.00847E-07 |
| orf19.2749 | 8.230224349 | 1.00847E-07 |
| orf19.279 | 8.230224349 | 1.00847E-07 |
| orf19.2836 | 8.230224349 | 1.00847E-07 |
| orf19.2853 | 8.230224349 | 1.00847E-07 |
| orf19.2963 | 8.230224349 | 1.00847E-07 |
| orf19.2995 | 8.230224349 | 1.00847E-07 |
| orf19.3030 | 8.230224349 | 1.00847E-07 |
| orf19.3226 | 8.230224349 | 1.00847E-07 |
| orf19.3353 | 8.230224349 | 1.00847E-07 |
| orf19.354 | 8.230224349 | 1.00847E-07 |
| orf19.3571 | 8.230224349 | 1.00847E-07 |
| orf19.3606 | 8.230224349 | 1.00847E-07 |
| orf19.3991 | 8.230224349 | 1.00847E-07 |
| orf19.4007 | 8.230224349 | 1.00847E-07 |
| orf19.4080 | 8.230224349 | 1.00847E-07 |
| orf19.4086 | 8.230224349 | 1.00847E-07 |
| orf19.4169 | 8.230224349 | 1.00847E-07 |
| orf19.4185 | 8.230224349 | 1.00847E-07 |
| orf19.4250 | 8.230224349 | 1.00847E-07 |
| orf19.4316 | 8.230224349 | 1.00847E-07 |
| orf19.4370 | 8.230224349 | 1.00847E-07 |
| orf19.4390 | 8.230224349 | 1.00847E-07 |
| orf19.4612 | 8.230224349 | 1.00847E-07 |
| orf19.4824 | 8.230224349 | 1.00847E-07 |
| orf19.4914 | 8.230224349 | 1.00847E-07 |
| orf19.5049 | 8.230224349 | 1.00847E-07 |
| orf19.5090 | 8.230224349 | 1.00847E-07 |
| orf19.5134 | 8.230224349 | 1.00847E-07 |
| orf19.515 | 8.230224349 | 1.00847E-07 |
| orf19.5296 | 8.230224349 | 1.00847E-07 |
| orf19.5376 | 8.230224349 | 1.00847E-07 |
| orf19.5752 | 8.230224349 | 1.00847E-07 |
| orf19.5935 | 8.230224349 | 1.00847E-07 |
| orf19.6027 | 8.230224349 | 1.00847E-07 |
| orf19.6205 | 8.230224349 | 1.00847E-07 |
| orf19.6341 | 8.230224349 | 1.00847E-07 |
| orf19.6528 | 8.230224349 | 1.00847E-07 |
| orf19.6732 | 8.230224349 | 1.00847E-07 |
| orf19.6989 | 8.230224349 | 1.00847E-07 |
| orf19.7067 | 8.230224349 | 1.00847E-07 |
| orf19.7102 | 8.230224349 | 1.00847E-07 |
| orf19.7260 | 8.230224349 | 1.00847E-07 |
| orf19.7478 | 8.230224349 | 1.00847E-07 |
| PDC2 | 8.230224349 | 1.00847E-07 |
| PDR17 | 8.230224349 | 1.00847E-07 |
| PHO100 | 8.230224349 | 1.00847E-07 |
| PZF1 | 8.230224349 | 1.00847E-07 |
| RPN5 | 8.230224349 | 1.00847E-07 |
| RRP42 | 8.230224349 | 1.00847E-07 |
| RSN1 | 8.230224349 | 1.00847E-07 |
| SAP2 | 8.230224349 | 1.00847E-07 |
| SLC1 | 8.230224349 | 1.00847E-07 |
| SPC34 | 8.230224349 | 1.00847E-07 |
| SPT20 | 8.230224349 | 1.00847E-07 |
| THG1 | 8.230224349 | 1.00847E-07 |
| TOP2 | 8.230224349 | 1.00847E-07 |
| TYE7 | 8.230224349 | 1.00847E-07 |
| UBP1 | 8.230224349 | 1.00847E-07 |
| YTH1 | 8.230224349 | 1.00847E-07 |

**Supplementary Table 5**. **Functional classification of proteins showing significantly different abundance (p-value < 0.05) between PUJ256 and SC5314 in the presence of fluconazole.** Proteins are grouped according to Gene Ontology (GO) biological processes based on the Candida Genome Database (CGD). The number in parentheses indicates the total number of proteins significantly more abundant in each strain within the indicated functional category. Proteins exclusively identified in one strain under fluconazole treatment are highlighted in grey.

| **PUJ256 Fluconazole (253)** | **SC5314 Fluconazole (261)** |
| --- | --- |
| **Response to stress** | |
| orf19.3140.1, orf19.6917, orf19.3649, Elc1, Hsp78, orf19.4914, Cta8 (7) | Swr1, Gpx2, Ecm4, Spt10, Rpt4, orf19.4522, Hda1, Mep1, orf19.2208, orf19.7425, Fcr1 (11) |
| **Filamentous growth** | |
| orf19.4953, Rpl6, Nnf1, Hmo1, Arf2, Bna4, Cef1, Ctm1, Top2, Drg1, Dip2 (11) | Hgt1, Rbd1, Ecm7, Gap1, Ctn3 (5) |
| **RNA metabolic process** | |
| orf19.4078, Rvb2, Rsc8, Rpl17b, Rpf2, Mex67, Wrs1, orf19.5987, Bud23, Ssz1, Mrt4, Prp3, Grs1, orf19.3482, Rat1, Ctr9, Frs1, Spl1, Rpl30, orf19.1026, Mtr2, orf19.500, Rrp46, orf19.6230, Taf145, Abp140, Dbp7, orf19.2400, orf19.5090, orf19.1525, orf19.4080, Pzf1, Rrp42, Thg1, Yth1, (36) | orf19.6360, Prp40, Puf3, orf19.3831, Tfc4, orf19.2017, Bur2, Sen15, Prp5, Slu7, orf19.6923, Rpc10 (12) |
| **Interaction between organisms** | |
| orf19.967, Cmp1, Imh3, Cnb1, Ade5,7, Def1, Gln3, Hem3, Sap2, Tye7, Pho100 (11) | Kex2, Pmc1, Rbt4, Phm7, Xog1, Tcc1, orf19.7194 (7) |
| **Cell cycle** | |
| Mcm6, Rfa1, Ctf5, Ctf8, (4) | orf19.6628, orf19.7006, orf19.6311, Rax1 (4) |
| **Response to chemical** | |
| Nma111, orf19.4474, Gpx31, Ptc5, orf19.6596, Gis2, Hpa2, Cta24, Oca1 (9) | Sod5, Pho84, Ddr48, Hms1, Hsp70, orf19.4258, orf19.7321, Sod3, Prx1, orf19.2476, Cta4, Pex1 (12) |
| **Vesicle-mediated transport** | |
| Boi2, orf19.2965, Sec23, Osh4, Arc35 (5) | Snx4, orf19.4184, orf19.2867, Pep12, Sec4, Bzz1, Sso2, Sro77, Syn8, orf19.4839, Ypp1, Sec7, Exo84, Gyp1, Gyp7, orf19.5418, orf19.2168.3 (17) |
| **Lipid metabolic process** | |
| Ipk2, Ebp1, Muq1, Mdh1-3, orf19.7478, Erg2, Slc1 (7) | Slp3, Dap1, Nsg2, Fas2, Lag1, Erg6, orf19.3684, Smf12, Acc1, Sur2, Flc2, Pex6, Plb1, orf19.3483 (15) |
| **Signal transduction** | |
| Ecs3, Bre1, orf19.6528, Spt20 (4) | orf19.5620, Ccr4, Ybl053, orf19.2728 (4) |
| **Cell wall organization** | |
| Ubc7 (1) | Sap10, Crh11, Chs3, Sap9, Pga31, Phr2, Kre6, Cmk1, Rac1, Fmp45 (10) |
| **Ribosome biogenesis** | |
| Rpl5, Rpl11, Rli1, Rps17b, Rpl12, orf19.6355 (6) | Rix7, Spb1, Bud21, orf19.3463, Sen1, orf19.1609, Rps12, Rlp24, Ria1, Pwp2, Noc2, orf19.3970 (12) |
| **Carbohydrate metabolic process** | |
| Xks1, Gal10, Fba1, orf19.6739, Rhr2, orf19.1355, Pgk1, Atc1, Mdh1, Ard, Ugp1, Kre5, orf19.5376, Bmt1 (14) | Snf6, Rho1, Sga1, Tps1, Pmr1, Gut1 (6) |
| **Protein modification process** | |
| orf19.5783, orf19.1626, Ptc7, orf19.3342, Ubp1, orf19.4007, Rpn5, orf19.3030 (8) | Png2, Rtk1, orf19.592, orf19.7326, Dun1, orf19.3615, Gtt12, orf19.1619, orf19.6581 (9) |
| **Cytoskeleton organization** | |
| Iqg1, Grx3, Cdc37, orf19.3235, Spc34 (5) | Bir1, Tub1, Bud14, Bem2, Rho3, orf19.6789, Scp1, Cdc42 (8) |
| **Biofilm formation** | |
| Med3, Fcr3, orf19.7199, Try2, Med20, Csr1 (6) | Atg13, orf19.7459, Gca2, Zcf15, Hyr1, Sap5 (6) |

#### 2 Supplementary Figures

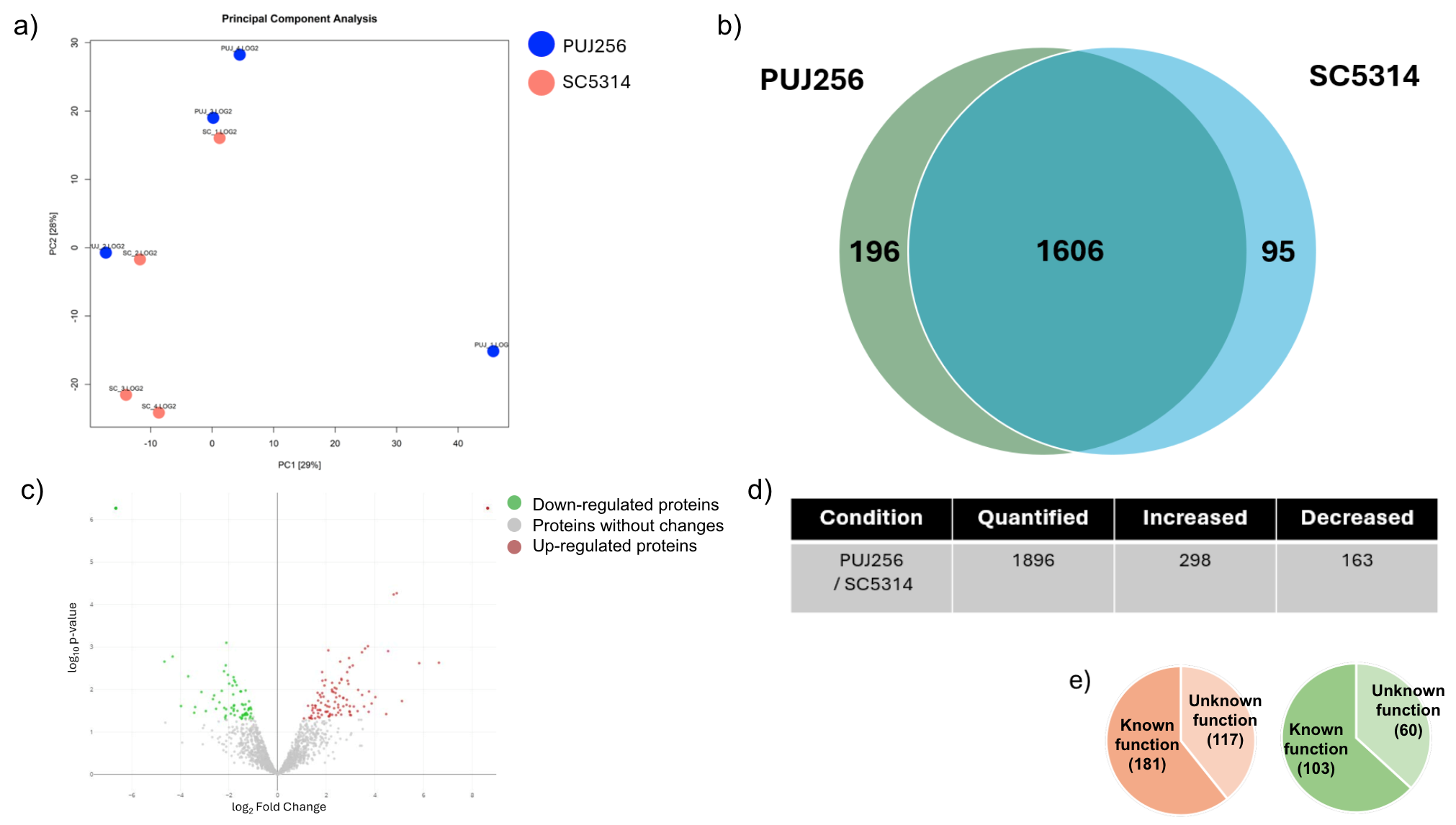

**Supplementary Figure 1.** a) Principal Component Analysis of SC5314 and PUJ256 proteome. b) Venn diagram showing common and non-common proteins with significant changes in abundance between strains in response to the treatment. c) Volcano plots representing proteins with significant changes in abundance. Significant changes in the protein abundance (−log10 p-value > 1.3) after treatment are presented in red for increase or green for decrease. d) Number of quantified proteins and proteins showing significant differences in abundance between both strains. e) Number of quantified proteins with significant differences in abundance whose function is unknown.

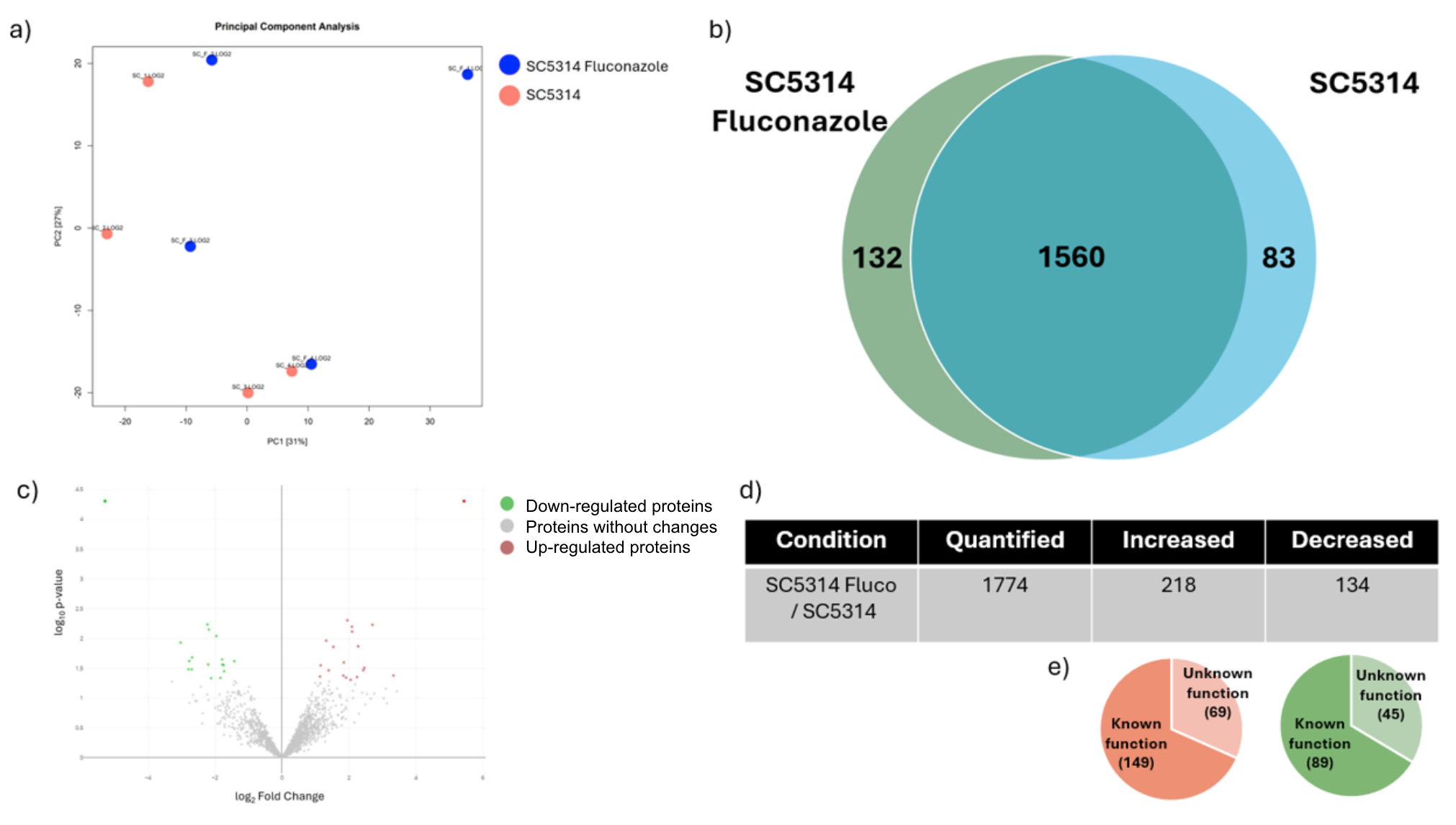

**Supplementary Figure 2.** . a) Principal Component Analysis of SC5314 with fluconazole (1.1 μg/mL) and SC5314 proteome . b) Venn diagram showing common and non-common proteins with significant changes in abundance between strains in response to the treatment. c) Volcano plots representing proteins with significant changes in abundance. Significant changes in the protein abundance (−log10 p-value > 1.3) after treatment are presented in red for increase or green for decrease. d) Number of quantified proteins and proteins showing significant differences in abundance between both conditions with fluconazole (1.1 μg/mL). e) Number of quantified proteins with significant differences in abundance whose function is unknown.

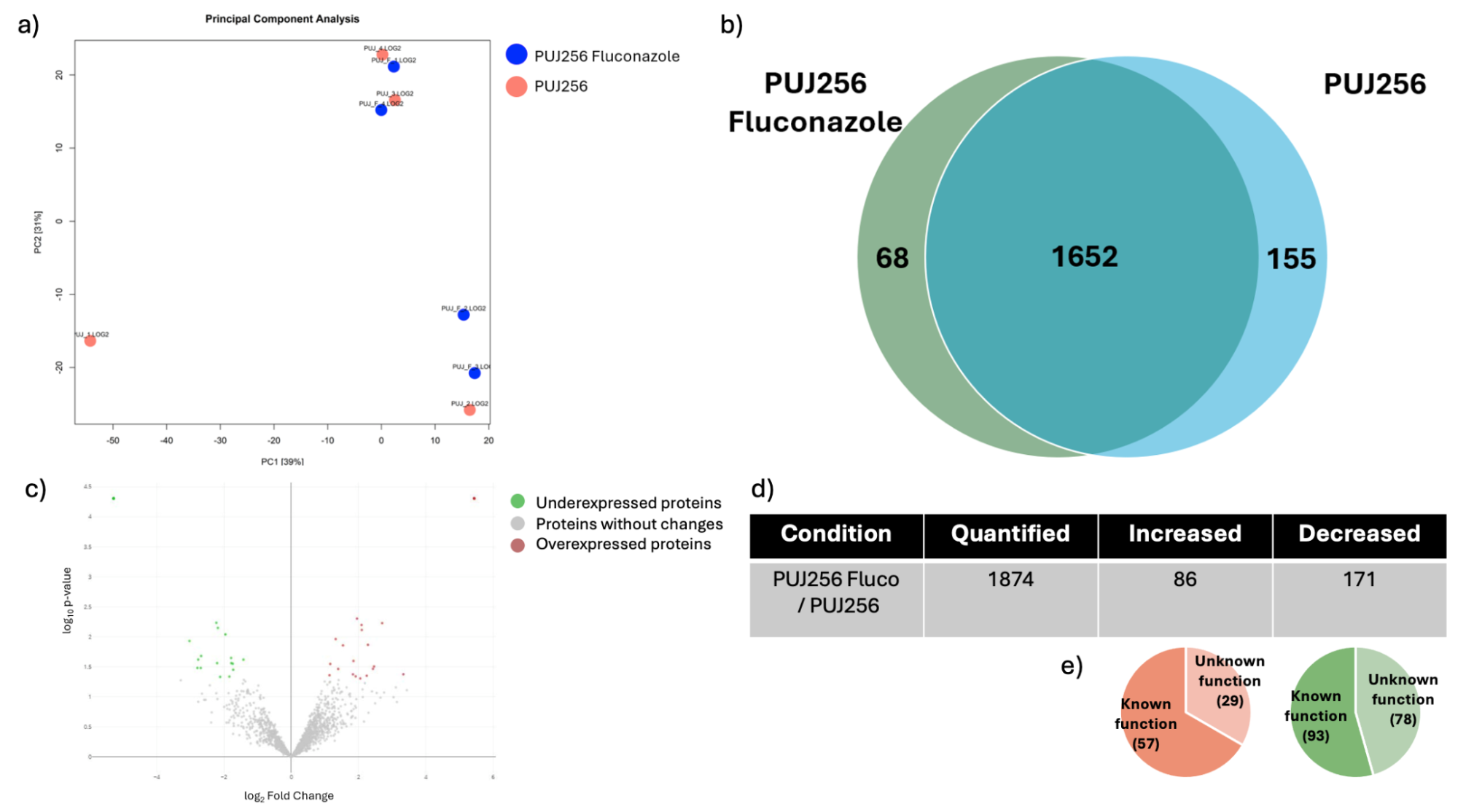

**Supplementary Figure 3.** a) Principal Component Analysis of PUJ256 with fluconazole (1.1 μg/mL) and PUJ256 proteome. b) Venn diagram showing common and non-common proteins with significant changes in abundance between strains in response to the treatment. c) Volcano plots representing proteins with significant changes in abundance. Significant changes in the protein abundance (−log10 p-value > 1.3) after treatment are presented in red for increase or green for decrease. d) Number of quantified proteins and proteins showing significant differences in abundance between both conditions. e) Number of quantified proteins with significant differences in abundance whose function is unknown.

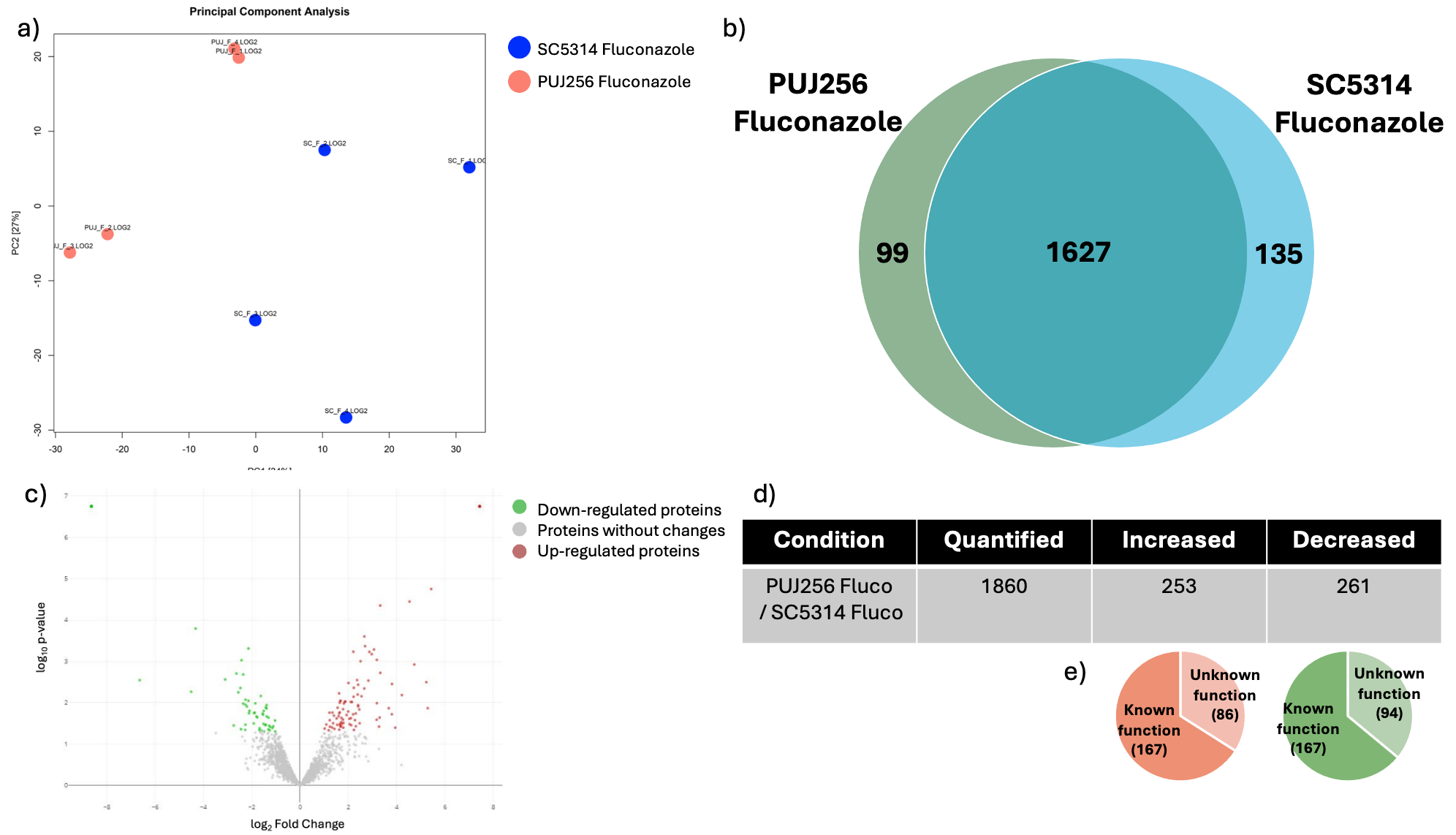

**Supplementary Figure 4.** a) Principal Component Analysis of SC5314 and PUJ256 proteome with fluconazole (1.1 μg/mL). b) Venn diagram showing common and non-common proteins with significant changes in abundance between strains in response to the treatment. c) Volcano plots representing proteins with significant changes in abundance. Significant changes in the protein abundance (−log10 p-value > 1.3) after treatment are presented in red for increase or green for decrease. d) Number of quantified proteins and proteins showing significant differences in abundance between both strains with fluconazole (11 μg/mL). e) Number of quantified proteins with significant differences in abundance whose function is unknown.
